# Mixed evidence for species diversity affecting ecological forecasts in constant versus declining light

**DOI:** 10.1101/2024.04.30.591860

**Authors:** Romana Limberger, Uriah Daugaard, Yves Choffat, Anubhav Gupta, Martina Jelić, Sabina Jyrkinen, Rainer M. Krug, Seraina Nohl, Frank Pennekamp, Sofia J. van Moorsel, Xue Zheng, Debra Zuppinger-Dingley, Owen L. Petchey

## Abstract

Accurate forecasts of ecological dynamics are critical for ecosystem management and conservation, yet the drivers of forecastability are poorly understood. Environmental change and diversity are considered major challenges to ecological forecasting. This assumption, however, has never been tested experimentally because forecasts have high data requirements. In a long-term microcosm experiment, we manipulated species richness of 30 experimental protist communities and exposed them to constant or gradually decreasing light levels. We collected finely-resolved time series (123 sampling dates over 41 weeks) of species abundances, community biomass, and oxygen concentrations. We then employed data-driven forecasting methods to forecast these variables. We found that species richness and light had a weak interactive effect on forecasts of species abundances: richness tended to reduce forecast accuracy in constant light but tended to increase forecast accuracy in declining light. These effects could partially be explained by differences among time series in variability and autocorrelation. Forecasts of aggregate properties (community biomass, oxygen), however, were unaffected by richness and light, and were not more accurate than those of species abundances. Our forecasts were based on time series that were detrended and standardized. Since real-world forecasting applications require predictions at the original scale of the forecasted variable, it is important to note that the results were qualitatively identical when back-transforming the forecasts to the original scale. Taken together, we found no strong evidence that higher diversity results in lower forecastability. Rather, our results imply that promoting diversity could make populations more predictable when environmental conditions change. From a conservation and management perspective, our findings suggest that diversity conservation might have beneficial effects on decision-taking by increasing the forecastability of species abundances in changing environments.

## Introduction

Forecasting the future state of populations, communities, and ecosystems is challenging but necessary to make well-informed decisions in ecosystem conservation and management (Deyle *et al*. 2022; Dietze *et al*. 2024, 2018; Tulloch *et al*. 2020). Forecasts are quantitative predictions of the future (Dietze 2017), i.e. they are based on models fitted to data from the past and present. Forecasts are evaluated by measuring the difference between observed and predicted values, referred to as forecast error, while its inverse is termed forecast accuracy or forecastability.

Ecological dynamics are difficult to forecast because organisms are embedded in complex networks of species interactions, respond simultaneously to many different drivers, often in a nonlinear way, and can display stochastic or chaotic dynamics (Beckage *et al*. 2011; Doak *et al*. 2008). However, global change and diversity loss have made ecological forecasting all the more relevant, resulting in rising numbers of published forecasts (Lewis *et al*. 2022). Typically, forecasts are based on observational data of organisms that are relevant in conservation and management (Lewis *et al*. 2022). Examples are forecasts of species that are harvested or hunted (Deyle *et al*. 2013; Henden *et al*. 2020), of pest outbreaks (Piou & Marescot 2023), or of harmful algal blooms (Rousso *et al*. 2020). Likewise, observational data has been used to compare forecast accuracy across different species and ecosystem properties (Lewis *et al*. 2022; Ward *et al*. 2014), in an attempt to gain a more general understanding about the constraints of forecastability.

Only few studies, however, investigated potential drivers of forecastability with experimental approaches (Benincà *et al*. 2008; Daugaard *et al*. 2022; Dumandan *et al*. 2024; Fujita *et al*. 2023). Forecasts of ecological dynamics benefit from long and finely-resolved time series (Dietze 2017), which can be challenging to accomplish with experiments. Yet, controlled experiments have been a fundamental tool in many fields of ecology (Jessup *et al*. 2004; Jochum *et al*. 2020; Petchey *et al*. 1999) and could be useful to advance our understanding about the potential and the limits of ecological forecasting.

One potential driver of forecastability is the diversity (i.e. species richness) of communities. This idea, however, has never been tested with an experimental manipulation of diversity. Previous work focused on comparing forecast accuracy among species within a community (Daugaard *et al*. 2022), rather than among communities with different properties (e.g. diversity). The diversity of a community could have both negative and positive effects on the forecastability of the species embedded in the community, as well as on aggregate ecosystem properties (e.g. community biomass). First, diversity could reduce forecastability by making the species interactions network more complex: communities with higher species richness have the potential for more direct and indirect interactions among species (Polis & Strong 1996; Yodzis 2000), making the outcome of species interactions less predictable (Montoya *et al*. 2009; Novak *et al*. 2011). Second, diversity could affect forecastability by influencing the temporal variability of populations and aggregate ecosystem properties. Higher diversity can both increase or reduce the temporal variability of populations (Xu *et al*. 2021), but usually reduces the temporal variability of aggregate ecosystem properties (Xu *et al*. 2021). The dampening effect of diversity on the temporal variability of aggregate properties could be particularly pronounced when environmental conditions change (Yachi & Loreau 1999), because diversity can buffer aggregate ecosystem properties against environmental changes (Baert *et al*. 2016; Isbell *et al*. 2015), and delay abrupt shifts to new ecosystem states (Limberger *et al*. 2023). Provided that lower variability results in higher forecastability, higher diversity could therefore increase the forecastability of community biomass (in particular when the environment changes), but reduce or increase the forecastability of species abundances. In general, aggregate properties are expected to be easier to forecast than species abundances because they might integrate over environmental variability (Lewis *et al*. 2023).

When environmental conditions change, ecological forecasting is particularly relevant but could also be particularly difficult. Of particular importance in a management context are near-term forecasts (Dietze *et al*. 2024, 2018), that is, forecasts of near-term variability rather than of long-term trends. Directional environmental change might alter such near-term variability if different environmental conditions lead to different population dynamics (Fussmann *et al*. 2014). Previous work showed that random environmental fluctuations can reduce forecastability (Daugaard *et al*. 2022) but we do not know if the same applies when the environment changes directionally to novel conditions, as is characteristic of many global change pressures (Tilman *et al*. 2001). Data-driven forecasting methods (e.g. machine learning) perform better when past time points hold more information about the future (Pennekamp *et al*. 2019). Population dynamics can differ among environmental conditions (Fussmann *et al*. 2000, 2014), and changes to novel environmental conditions could therefore reduce forecastability due to greater dissimilarity of past and future dynamics. In addition, abiotic change often alters species interactions (Blois *et al*. 2013; Gilman *et al*. 2010), which could further constrain forecastability (Dumandan *et al*. 2024). On the other hand, environmental change can reduce stochasticity (Blasius *et al*. 2020) and increase autocorrelation of population time series (Drake & Griffen 2010; Pace *et al*. 2017). Consequently, future states of such systems depend more on past states, and should thus be more predictable (Pennekamp *et al*. 2019). Despite the importance of ecological forecasting in changing environments, we lack empirical studies that test how directional change to novel environmental conditions influences forecastability.

Here we investigated how diversity and environmental change affect the forecastability of ecological dynamics in a controlled and simplified system. We conducted a microcosm experiment with a unique combination of high numbers of time points (123), experimental units (30), and experimental taxa (18). We used aquatic protists as model organisms because they have short generation times (∼ 0.5-3 generations per day under ideal growth conditions (Leary & Petchey 2009; Tabi *et al*. 2019)), which allows to collect time series of population dynamics within a relatively short time frame. We assembled communities with three different levels of species richness and exposed the communities to constantly high light levels or to light that declined gradually and then remained constantly at low levels. We collected time series of species abundances and aggregate ecosystem properties (community biomass and oxygen) and used data-driven forecasting models to forecast these variables. With this dataset we tested four hypotheses (Supplement Figure S2 a, b): (1) higher diversity leads to higher forecast error of species abundances (Hypothesis 1, H1), (2) historical environmental change to novel conditions increases forecast error of species abundances (H2), (3) forecast error of aggregate ecosystem properties decreases with diversity, in particular in changing environments (H3), and (4) forecast error of aggregate ecosystem properties is lower than forecast error of species abundances (H4).

Our experimental communities and treatments are simplified versions of real-world scenarios. Our manipulation of light is meant to represent anthropogenic pressures that are changing gradually and directionally (Tilman *et al*. 2001). We chose light because it is relevant in a global change context (Leech *et al*. 2018; Winder & Sommer 2012) but also because we expected that light would affect the growth of many species in the community, in particular that of primary producers (algae), but indirectly also that of consumers. However, we did not intend to mimic the complex characteristics of human-induced light regime changes in natural aquatic systems: these changes include both increases in light availability (due to prolonged lake stratification (Winder & Sommer 2012)) and decreases (e.g. due to the browning of lakes (Leech *et al*. 2018)), and they can be associated with concurrent changes in other factors (e.g. browning resulting from increased input of dissolved organic matter). Likewise, our manipulation of species richness (maximum: 14 species) is a simplified (i.e. shorter) version of diversity gradients observed in natural systems. To allow high-frequent samplings, the size of our species pool was limited because we only used species that were distinguishable with automatic classification. Although our communities are simpler than their natural counterparts, they represent a model of an important community type, that is, of a food web dominated by consumer-resource interactions. With this simple and tractable system we aimed to provide first experimental insights into the effects of diversity and environmental change on ecological forecasts.

## Methods

### Experimental design

In a nine-month experiment, we manipulated light (2 levels) and species richness (3 levels) in a full factorial design. At each of the three richness levels, we assembled five different community compositions. The composition × light combinations were not replicated, i.e. from each of the 15 compositions we set up two microcosms, one for each of the two light levels, resulting in a total of 30 microcosms. Light intensity was either constantly high or declined gradually. To manipulate light, we used eight incubators (IPP260plus, Memmert), four per light level. The declining light treatment involved three phases: (1) three months of constantly high light (30% of the maximum incubator light intensity, ∼ 22,500 Lux), (2) three months of gradual light decline (from 30% to 1%, ∼ 800 Lux), and (3) three months at the final low light level (Supplement Figure S3). Throughout the entire experiment, all incubators were set to a light:dark cycle of 16:8 hours and a temperature of 18°C.

We manipulated species diversity by assembling protist communities with different numbers of species (7, 10, and 14 species, respectively). We used a pool of 18 species of algae and ciliates, comprising six functional groups: algae which were edible for ciliates (two species), inedible algae (5), bacterivorous ciliates (4), omnivorous ciliates (4), mixotrophic ciliates (2), and a predatory ciliate (1) (Supplement Table S1, Figure S1). The three diversity levels were generated by manipulating diversity within the functional groups of inedible algae, bacterivores, and omnivores (one, two, or three species per functional group), and in the mixotrophs (one, one, or two species) (Supplement Table S2). To avoid confounding effects of species richness and species identity/composition (Huston 1997; Schmid *et al*. 2017), we assembled five different compositions at each diversity level. The compositions were as dissimilar as possible, and species occurred with similar frequency both within and across diversity levels (see Supplement Section S1.3 for computation of community compositions). Each of the 15 compositions comprised all six functional groups.

### Implementation of the experiment

Over a period of four weeks, we inoculated 30 2L-bottles from stock cultures growing at high densities. Each bottle was filled with 1 L of medium (75% filtered protist pellet medium, i.e. 0.55 g protist pellet in 1L Chalkley’s medium, and 25% WC medium), 20 wheat seeds for slow release of nutrients, and a magnetic stirrer for homogenization during samplings. Diversity was manipulated in a substitutive design, e.g. the culture volume added of inedible algae was divided among one to three species depending on the diversity level. During inoculation, we homogenized bacteria associated with algae and ciliate cultures so that communities did not differ in their initial bacterial composition. See Supplement Section S1.5 for further details on inoculation.

After the inoculation phase, we sampled the 30 bottles three times per week (Monday, Wednesday, Friday) for nine months (41 weeks), resulting in 123 sampling days. On sampling days, we removed 50 mL from each bottle after homogenization on a magnetic stirrer and then added 50 mL of fresh medium to the bottle. Samples were taken under sterile conditions using 50-mL glass pipettes.

To allow species to re-colonize bottles if they had gone extinct, we added the species from the stock cultures every three weeks, mimicking a very low rate of immigration. In protist microcosms, species often go extinct through competitive or trophic interactions and would have no chance of re-colonizing the microcosm when conditions become favourable. At these immigration events, we added a very low number (10 individuals) of each species that was part of the community of a given bottle. Soon after the start of the experiment, two species went extinct in the stock cultures (*Stylonychia mytilus* and *Didinium nasutum*) and could therefore not be re-immigrated. Stock cultures were cultivated in high light conditions (i.e. 30% light).

### Quantification of species abundances and ecosystem properties

We measured abiotic ecosystem properties (dissolved oxygen, carbon, and nitrogen), species abundances of ciliates and algae, and total abundance of bacteria. To quantify dissolved oxygen, we used a hand-held oxygen meter and two optical oxygen sensors attached to the inner wall of each bottle. Oxygen was measured in the morning prior to sampling. The light:dark cycle of the incubators was set such that the oxygen measurements would be made towards the end of the 16-hour light phase, specifically after 13.5 hours of light. We measured dissolved organic carbon (DOC) and dissolved nitrogen with a TOC/TN analyzer, bacterial abundance with a flowcytometer, algae abundances and morphological traits with a FlowCAM, and ciliate abundances and movement and morphological traits with video microscopy (R-package *bemovi* (Pennekamp *et al*. 2015)). The predatory ciliate *Didinium* was counted manually under a dissecting microscope when present. See Supplement Section S2 for details on measurements and instruments used.

We identified the recorded algae and ciliate individuals by using support vector machine (SVM) classifiers (R-package e1071 (Meyer *et al*. 2021)). We trained these classifiers with trait data collected from monocultures cultivated at different light levels and, if necessary, with manually annotated data from the experiment. The trait data to train the classifiers was collected with the same methods used in the experiment, i.e. video microscopy for ciliate traits and FlowCAM for algae traits. See Supplement Section S2.6 for further details on classifications.

Lastly, we computed community biomass by summing the biomass of all algae, ciliates, and bacteria in a sample. To this end, we estimated the volume of each particle using its extracted traits and three-dimensional shape. Further details are given in Supplement Section S4.3.

### Processing of recorded time series

After data collection and classification, we carried out several quality controls and made corrections if needed (for a complete description see Supplement Section S3). Then, we averaged over technical replicates of density measurements made with flowcytometry and video microscopy, respectively, and of oxygen concentration measurements made with the oxygen meter.

The recorded data included some missing data (flowCAM: 0.81%, flowcytometer: 4.84%, videos: 0.84%, oxygen meter: 1.67%, TOC/TN analyzer: 0.92%, Supplement Table S5). We used cubic hermite splines to impute the missing data and to interpolate the time series such that all time points were equidistant (time step of 2.3 days). To make the time series stationary, we detrended and standardized them, as is commonly done in time series analyses (Benincà *et al*. 2008) by taking the standardized residuals of the regressions of the recorded values against time. That is, the purpose of our forecasts was to predict the variation around a trend, not the trend itself. Since forecasts of detrended time series can be back-transformed to the original scale of the data, detrending does not preclude the use of forecasts when the actual values are needed (e.g. in conservation and management). To test the robustness of our results, we also evaluated the forecasts after back-transforming them to the original scale of the forecasted variables (Supplement Section S7.5).

We detrended and standardized the entire time series, i.e. the test data was included in the processing of the time series. This approach allows for the best possible time series transformation and is therefore often used in theoretical studies that compare forecast accuracy among treatments or species (Benincà *et al*. 2008; Daugaard *et al*. 2022; Munch *et al*. 2023). In real-world settings, however, the test data is not available prior to making forecasts. We therefore checked if our results changed when using only the historic part of the time series (i.e. the 111 time points of the training dataset) to detrend and standardize the time series (Supplement Section S7.4).

The untransformed and transformed time series are displayed in Supplement Figures S7-S14. Note that the log-transformation in figures of untransformed time series (Figures S7-S9) is for better visibility of the time series. However, the data was not log-transformed prior to detrending.

### Forecasting

We forecasted species abundances and two aggregate ecosystem properties (total community biomass and oxygen concentration). The forecasted species (i.e. forecast targets) included the ciliate and algae species, bacteria (considered one group), and additional morphotypes (“*Chlamydomonas* clumps”, “*Desmodesmus* clumps”, “Dividing *Chlamydomonas*” and “Small cells”, i.e. small unidentified cells), 14 targets in total. However, we did not forecast the abundance of a target if it went extinct during the experiment (Supplement Section S3.2). Across all bottles, we forecasted 253 taxa time series.

We used different forecast methods to ensure that the results were independent of the chosen methodology. We used one linear model (Auto-Regressive Integrated Moving Average, i.e. ARIMA), two nonlinear approaches based on Empirical Dynamic Modeling (Simplex EDM and Multiview EDM (Sugihara & May 1990; Ye & Sugihara 2016)), and two machine learning methods (Random Forest RF and Recurrent Neural Networks RNN). Simplex and Multiview EDM differ insofar as the former allows better parameter optimization, while the latter is designed for high-dimensional systems. See Supplement Section S5 for a more detailed description of the forecast models and their parametrization.

We used the first 111 time points of the time series to train the models and then forecasted the remaining 12 time points (i.e. ∼10% of the data, corresponding to the last 4 weeks of sampling) with one-step-ahead forecasts. Two of the forecast methods (ARIMA, Simplex EDM) included only the forecast target itself as predictor, whereas the other three methods included additional predictors. As predictors we used all forecast targets of a given bottle as well as the dissolved carbon and nitrogen concentrations. The predictor data did not include the forecasted time step. We quantified the forecast error (i.e. the disagreement between forecasted and recorded values) as the root mean square error (RMSE). Because we forecasted standardized time series, an RMSE value below one indicates a better performing model than the average value of the time series (which has an expected RMSE of one).

### Analyzing the effects of species richness and light on forecast error

To analyze the effects of the experimental treatments on forecast error, we computed linear mixed models (R-package *lme4* (Bates *et al*. 2015)), following common practice in similarly designed experiments (Pennekamp *et al*. 2018; Steiner *et al*. 2006). In separate models, we used forecast error of either taxa abundances, community biomass, or oxygen as response variable. As explanatory variables, we used the experimental treatments, i.e. light condition (binary), species richness (continuous), and their interaction. We used type III ANOVA to assess the importance of covariates and interactions. If the interaction was significant or marginally significant (i.e. *p*-values below 0.1), we kept the interaction in the model and used type III ANOVA. Otherwise, we removed the interaction from the model and used type II ANOVA to investigate the importance of the main fixed effects. In the case of a significant interaction, we analyzed the effect of species richness with a separate linear mixed model at each of the two light levels (Quinn & Keough 2024), and the effect of light at the lowest and highest richness level with a post-hoc analysis that estimated the marginal means.

Observed species richness deviated from planned richness because not all species persisted throughout the experiment (Supplement Section S3.1). Furthermore, in one bottle the alga *Desmodesmus* was introduced by cross-contamination. We therefore used realized richness (i.e. the median of the observed richness values) rather than planned richness in our models.

Realized species richness was unaffected by the light conditions and was positively correlated with planned species richness (for more information and alternative realized species richness metrics see Supplement Section S4.1). We also evaluated if forecast error was related to further diversity indices (Supplement Section S9).

In all models, we included random intercepts following the experimental design. That is, models to investigate treatment effects on the forecast error of taxa abundances included random intercepts for taxon, bottle, and incubator, whereas models to investigate effects on the forecast error of aggregate properties included random intercepts for composition and incubator.

We also investigated if treatment effects depended on the level of aggregation of the response variable by including the mean forecast error of taxa abundances and the forecast error of community biomass in the response variable. As explanatory variables, we used realized richness, light condition, a binary variable that specified the level of aggregation (taxa abundances vs. community biomass), and all possible pairwise interaction terms. Bottle and incubator were included as random effects. We assessed the importance of model covariates with type III or type II ANOVA depending on the *p*-value of the interaction, as described above. We repeated this model for the forecast error of oxygen instead of community biomass.

We repeated the analyses for each of the five forecasting methods. As treatment effects on forecast error were similar among the five approaches, we chose one method (Simplex EDM) to describe the treatment effects in the main text and refer to the supplementary material for results regarding the other four methods.

### Explaining the treatment effects on forecast error

To explore possible mechanisms behind observed treatment effects on forecast error, we investigated how forecast error was related with time series properties. For each time series, we calculated three metrics: the coefficient of variation, autocorrelation, and permutation entropy. The coefficient of variation is a measure of temporal variability and was calculated as the ratio of the standard deviation of the detrended time series and the mean of the original time series. Autocorrelation describes the correlation of current and past values and was calculated for a lag of 5 (i.e. correlation with values 5 time points ago); the choice of lag did not affect the results. Permutation entropy is a measure of time series complexity, with higher permutation entropy usually being associated with higher forecast error (Pennekamp *et al*. 2019). We calculated autocorrelation and permutation entropy based on the transformed time series. See Supplement Section S4.2 for further details.

We used two approaches to investigate the relationship of the time series metrics with the experimental treatments and with forecast error of species abundances. First, we computed separate linear mixed models to test (i) how the experimental treatments (realized richness, light, and their interaction) affected each of the three time series metrics, and (ii) how each of the three metrics, light condition, and their interaction affected forecast error. Again, we included random intercepts for taxon, bottle, and incubator in the models, and assessed the importance of model covariates with type III or type II ANOVA as described above. As a second approach, we computed a structural equation model (SEM) to further clarify the relations found with the first approach, using the R-packages *lavaan* (Rosseel 2012) and *lavaan.survey* (Oberski 2014). *Lavaan.survey* allowed us to include random effects according to our experimental design. We set up the model to estimate the relations between the variables in both light conditions by using a multiple group analysis, and tested the significance of SEM path coefficients with two-sided t-tests. All data analyses and forecasts were done in R (version 4.3, package information in Supplement Section S10).

## Results

### Effects of species richness and light on forecast error

Richness and light had a weak interactive effect on the forecast error of taxa abundances (richness x light: *p*-value= 0.0363; Figure 1a, Table 1, forecast method: Simplex EDM). The statistical support for this effect was somewhat limited (Figure 1a: overlapping confidence intervals), potentially due to high amounts of variance in the data. However, the interactive effect was consistent among forecast method, detrending approach, and scale of forecast error calculation (see below). The interaction of the two treatments resulted from opposite effects of species richness in constant versus declining light: forecast error tended to increase with species richness in constant light but tended to decrease with richness in declining light (Figure 1a). Specifically, when median richness increased by one species, forecast error increased by 5.6% (95% CI: −0.8% – 12.1%) in constant light and decreased by 4.6% (95% CI: −11.0% – 1.0%) in declining light relative to the baseline forecast error of 1 (i.e. the expected forecast error when the average value of a time series is used as the prediction). Note that the 95% confidence intervals of the estimates include zero despite the interaction term p-value < 0.05. A discrepancy such as this can occur when the statistical evidence of an effect is not very strong (e.g., *p*-value = 0.0363) and when different methods are used to calculate *p*-values and confidence intervals. Specifically, approximate *p*-values were calculated with Wald tests, whereas confidence intervals were calculated with profile likelihoods due to the greater precision of this approach.

**Figure 1:**
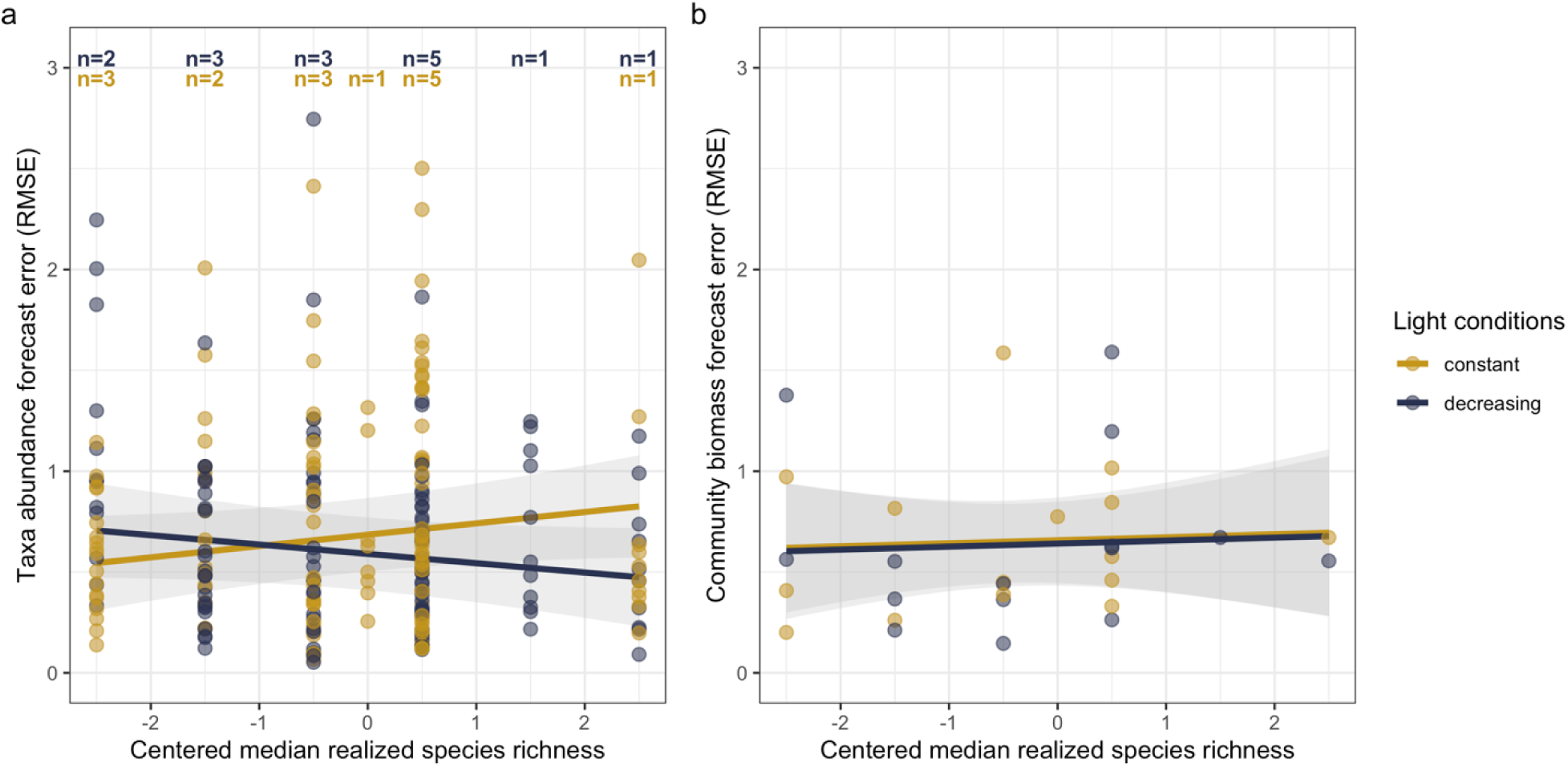
Effects of species richness and light on forecast error. The two treatments had interactive effects on forecast error of taxa abundances (a) but no effect on the forecast error of community biomass (b). Forecasts were computed with Simplex Empirical Dynamic Modeling (EDM), and forecast error was calculated as root mean square error (RMSE). RMSE is standardized (unitless). Observed richness deviated from planned richness because not all species persisted throughout the experiment. We therefore used realized richness (i.e. the median of the observed richness values) as explanatory variable. Median richness was centered; a centered median richness of zero corresponds to a median richness of 5.5 (minimum median richness: 3; maximum median richness: 8). Lines display the fit of the linear mixed model; shaded areas denote the 95% confidence intervals. In (a), we forecasted 253 taxa time series (across all bottles); each point represents a forecasted taxon in a bottle. Taxon, bottle, and incubator were included as random intercepts in the model. Inserts at the top indicate the number of bottles for the respective richness level (yellow: constant light, blue: decreasing light). In (b), we forecasted community biomass in 30 bottles. Composition and incubator were included as random intercepts in the model.

**Table 1:**
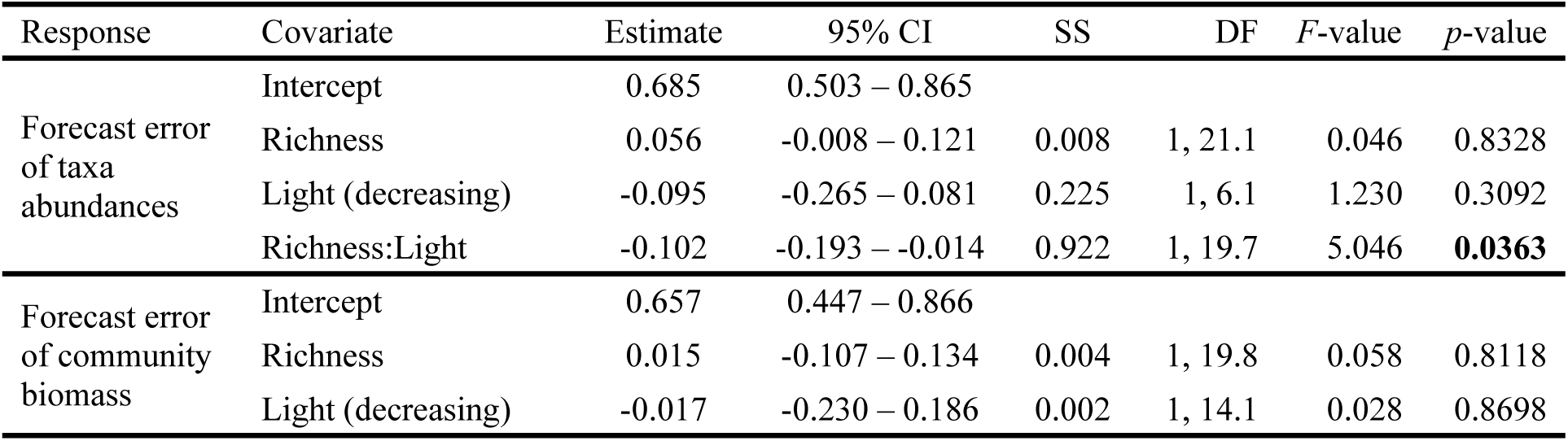
Results of two linear mixed models investigating the effects of species richness and light on the forecast error of taxa abundances and community biomass, respectively. We forecasted 253 taxa time series (across all bottles) and community biomass in 30 bottles. In both models, median centered richness and light treatment were fixed effects. The first model (response: forecast error of taxa abundances) included taxon, bottle, and incubator as random intercepts. The second model (response: forecast error of community biomass) included composition and incubator as random intercepts. Forecast error is based on forecasts computed with Simplex Empirical Dynamic Modeling (EDM). In the second model, the interaction was not significant (*p*-value > 0.1) and was therefore removed from the model. The last four columns report the ANOVA results. P-values < 0.05 in bold. See Supplement Table S6 for further details.

Separate analyses at constant and declining light conditions also indicated opposite richness effects on taxa abundance forecasts in the two light conditions, as the slopes of the two relationships had opposite signs and neither of their 95% confidence intervals overlapped with zero (Supplement Table S7). Furthermore, we investigated the effect of the light decline on the forecast error of taxa abundances at the low and high end of the diversity gradient. The light decline reduced forecast error at high species richness, but had no effect at low species richness (tested with a post-hoc analysis; Supplement Table S9).

The interactive effect of richness and light on the forecast error of taxa abundances persisted in several robustness analyses (Supplement Section 7). Both the direction and the size of the treatment effects were similar among the five forecasting methods (Supplement Table S8, Figure S18, S21). Likewise, treatment effects were similar among richness variables (including planned richness) and detrending method and were not driven by individual species (Supplement Figures S21-S24). Furthermore, treatment effects were similar as above when using only the historical data to detrend and standardize the time series (Supplement Figure S26) and when evaluating the forecasts after back-transforming the forecasts to the original scale (Supplement Figure S28).

Forecastability of aggregate ecosystem properties was largely unaffected by the experimental treatments. Light and richness had no effect on the forecast error of total community biomass (Table 1, Figure 1b) except when biomass was forecasted with RNN (Supplement Table S10, Supplement Figure S19). In this case, treatments had similar effects as on forecast error of taxa abundances. Forecast error of oxygen was unaffected by light and richness (Supplement Table S11, Supplement Figure S20). For most forecasting methods, forecast error of aggregate properties (community biomass, oxygen) did not differ from average forecast error of species abundances (Supplement Tables S12-S14).

### Explanations of the treatment effects on forecast error

To investigate possible reasons for differences in forecast error, we analyzed how the forecast error of species abundances was related with three time series properties: autocorrelation, coefficient of variation (as a measure of temporal variability), and permutation entropy (as a measure of time series complexity (Pennekamp *et al*. 2019)). We expected that greater autocorrelation and lower permutation entropy would be associated with lower forecast error, while greater variability (i.e. higher coefficient of variation) could be associated with lower or higher forecast error, depending on whether higher variability is associated with more or less determinism (Supplement Figure S2 c-e).

Forecast error decreased with increasing autocorrelation and coefficient of variation, and increased with increasing permutation entropy (Figure 2 a-c, Supplement Table S15). The negative relationship between forecast error and the coefficient of variation suggests that higher temporal variability reflected higher deterministic variability (rather than higher stochastic variability). The experimental treatments affected the three time series metrics, albeit not in an interactive way: autocorrelation and coefficient of variation were lower in declining light than in constant light, whereas permutation entropy was unaffected by light but increased with richness (Figure 2 d-f, Supplement Table S16).

**Figure 2:**
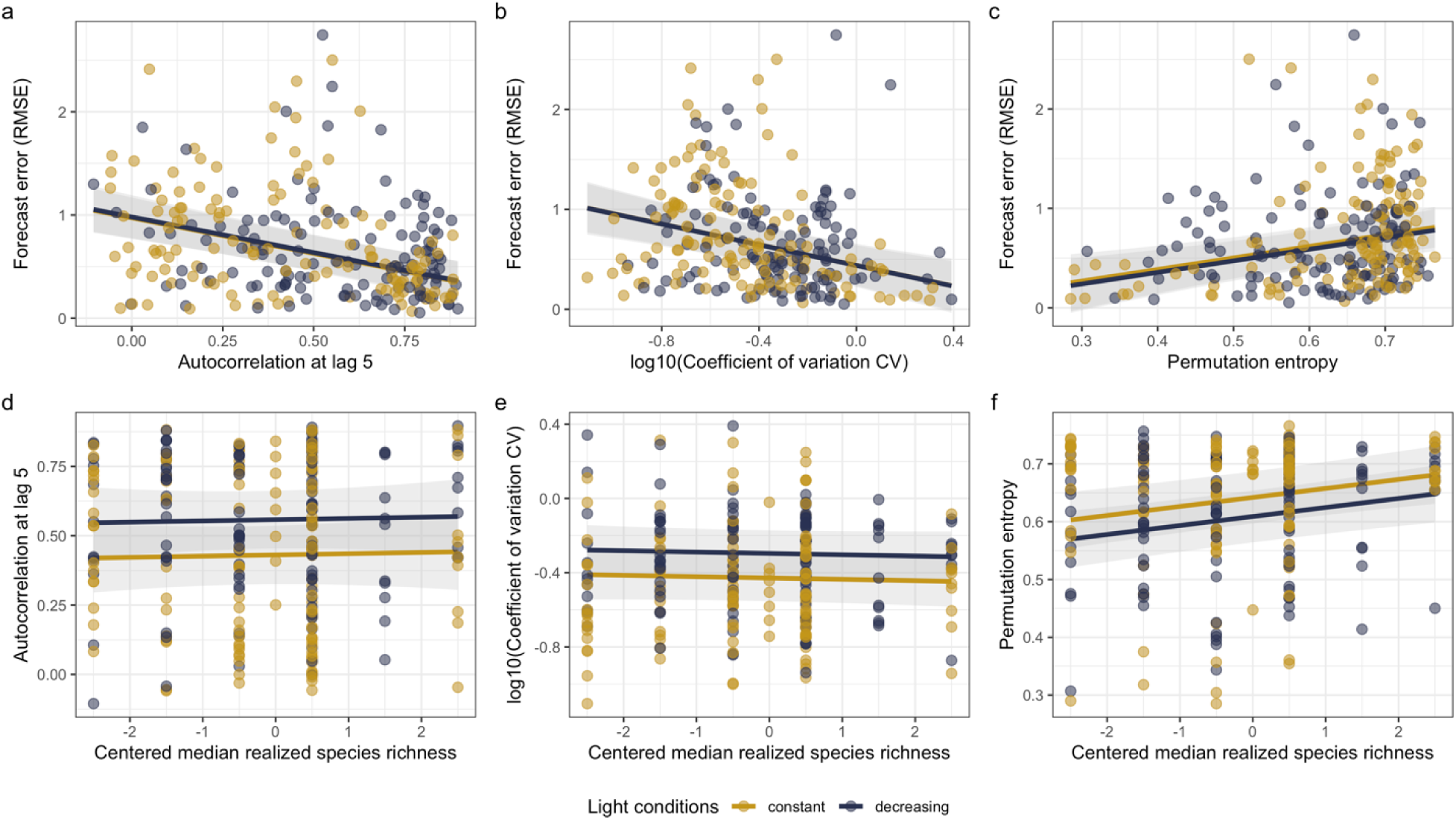
Relationship of time series metrics with forecast error (a-c), and with experimental treatments (d-f). Time series metrics are (a, d) autocorrelation at lag 5, (b, e) coefficient of variation as a measure of temporal variability, and (c, f) permutation entropy as a measure of time series complexity. All variables are unitless. Time series metrics were calculated for 253 taxa time series (across all bottles). Lines display the fit of the corresponding linear mixed model, shaded areas denote the 95% confidence intervals. Taxon, bottle, and incubator were included as random effects in the model. Forecast error is based on forecasts computed with Simplex EDM. Note that in (b) and (e) the coefficient of variation was log_10_-transformed so that linear model assumptions were met.

We used a structural equation model (SEM) to disentangle direct and indirect effects of richness and time series metrics on forecast error of taxa abundances (Figure 3, Supplement Table S17). A multi-group model provided a good match between model-implied and observed covariance (*χ*^2^ = 3.219, DF = 4, *p* = 0.522, i.e., no evidence against a good model fit). The model explained 32.2% of the variation in forecast error in constant light, and 9.7% of the variation in forecast error in declining light (R^2^, Table S18). In line with the bivariate correlation analyses, the SEM showed in both light conditions that forecast error declined with increasing autocorrelation, declined with an increasing coefficient of variation, and increased with increasing permutation entropy (Figure 3; note that not all of these effects were statistically significant).

**Figure 3:**
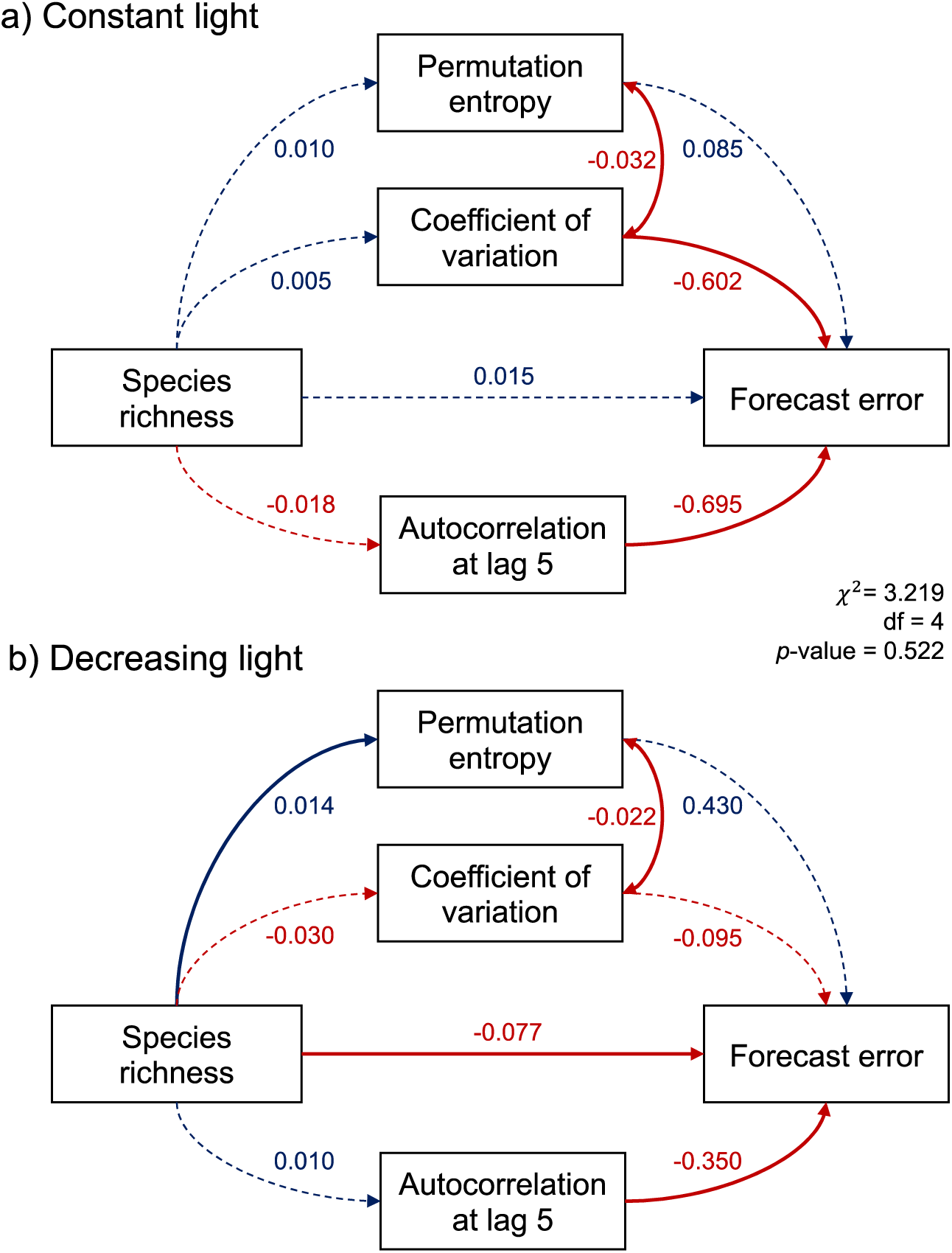
Structural equation model (SEM) showing the effects of species richness and time series metrics on forecast error of species abundances in (a) constant light, and (b) decreasing light. Solid and dashed arrows indicate significant and non-significant effects, respectively (see Supplement Table S17). Values next to arrows are the path coefficients. The color matches the sign of the coefficients (blue=positive, red=negative). Bidirectional arrows are covariances between the error terms of the involved variables.

Similarly as the linear mixed model analysis, the SEM showed that the effect of richness on forecast error of taxa abundances depended on the light treatment. Richness had a positive (albeit not significant) effect on forecast error in constant light but significantly reduced forecast error in decreasing light (Figure 3, Supplement Table S17). Furthermore, the effect of species richness on the coefficient of variation and on autocorrelation was reversed between the two light treatments (Figure 3). Although not significant, these reversed effects of richness on coefficient of variation and autocorrelation in the two light treatments might explain why increasing richness had opposite effects on forecast error in the two light treatments.

## Discussion

Across the diversity gradient used in this experiment, diversity had only weak effects on the forecastability of species abundances. Diversity tended to increase forecastability (i.e. reduce forecast error) of species abundances in declining light but tended to reduce forecastability in constant light. These effects, although weak, were consistent among different forecast methods, detrending approaches, and scales of forecast error calculation. In summary, we found only weak support for the hypothesis that higher diversity makes population dynamics hard to forecast (H1), and this (weak) negative effect of diversity on forecastability was only observed in constant light. Our hypothesis that environmental change reduces forecastability (H2) was not supported. On the contrary, light decline increased forecastability when richness was high but had no significant effect when richness was low. Forecastability of aggregate properties (community biomass, oxygen) was unaffected by our experimental treatments (H3) and was not higher than that of species abundances (H4). Taken together, we found no strong evidence that higher diversity leads to low forecastability, which could be partly due to the comparatively short diversity gradient. However, our results suggest preliminary support for higher diversity increasing the forecastability of species abundances in changing environments. Diversity effects on forecastability could be stronger across broader diversity gradients, which calls for further research in more complex systems with larger diversity gradients to explore the generality of our findings.

Light decline increased the forecastability of species abundances at the upper end of the diversity gradient. This positive effect of light decline on forecastability potentially resulted from higher autocorrelation in changing conditions (Figure 2d). Higher correlation between successive time points means that past states have higher information content, resulting in higher accuracy of forecasts (Ward *et al*. 2014). Accordingly, higher autocorrelation was associated with lower forecast error (Figure 2a, Figure 3). Higher autocorrelation in changing conditions was due to both direct and indirect effects of the environmental driver. Decreasing light put some species on deterministic trajectories of population decline, which indirectly led to abundance increases of inferior competitors and prey organisms (Supplement Figures S29, S30). Such indirect effects of environmental change via altered species interactions are widespread (Amundrud & Srivastava 2016; Bell *et al*. 2019; Martin & Maron 2012), and may be more frequent than direct effects (Ockendon *et al*. 2014). As such, autocorrelation can spread through the community even if only few species are directly affected by environmental change. Taken together, our results illustrate that forecastability of species abundances can be high in changing environments if direct and indirect effects of environmental change result in higher autocorrelation of time series, and therefore in greater information content of past states.

In the declining light treatment, forecastability of species abundances tended to decrease as diversity decreased. We can only speculate about a possible mechanism behind this pattern. Possibly, high forecast error at the low end of the diversity gradient might have resulted from declining light leading to previously unobserved dynamics. Specifically, in the two communities with lowest diversity, an entire functional group (consumers of algae, i.e. omnivores + mixotrophs) was represented by only one species, which declined or suddenly increased in response to the environmental driver, resulting in pronounced changes in prey abundances (Supplement Section S8.1). In more diverse communities, however, the consumer functional group thrived across the entire light gradient and did not change suddenly as conditions changed. For example, the decline of the dominant consumer in the decreasing light treatment was compensated by increases in other consumer species (Supplement Section S8.1). This process is known as the insurance effect of diversity (Loreau *et al*. 2021; Yachi & Loreau 1999), i.e. the loss of species due to environmental change is compensated by other species that perform similar functions but tolerate and potentially thrive in the new environmental conditions (Bestion *et al*. 2020; Hong *et al*. 2022; Isbell *et al*. 2015). This mechanism might have contributed to the positive effect of diversity on forecastability in the changing light treatment by maintaining functional groups in the community, thereby preventing pronounced changes in population dynamics.

The statistical support for the treatment effects was somewhat limited, possibly reflecting high variance in the data. Having more replicates of each richness × composition × light combination would be useful in future research to gain stronger statistical support for treatment effects. In the case of the light treatment, high variance might have resulted from counteracting effects of light on forecastability. As discussed above, it seems that environmental change had contrasting effects on forecastability, that is, light decline increased forecastability by increasing autocorrelation but reduced forecastability by leading to previously unobserved dynamics. The first of these effects (higher autocorrelation) dominated when diversity was high, whereas the second effect (change in dynamics) dominated when diversity was low. Still, however, both effects likely occurred along the entire diversity gradient, with the importance of these effects likely differing among species and compositions. These contrasting effects of environmental change on forecastability might have contributed to the high variation in the data.

The limited statistical support for the diversity effect on taxa abundance forecasts could be partly due to the rather short diversity gradient. To enable high-frequent samplings, we only used species that were distinguishable by automatic classification, which limited the size of the species pool. This limitation is typical for studies on ecological forecasting: the high number of required time points constrains studies to focusing either on comparatively few species (Daugaard *et al*. 2022; Dumandan *et al*. 2024) or on few experimental units (Benincà *et al*. 2008). Despite this limitation, our diversity gradient was in the range of previous studies that used experimental protist communities to draw important conclusions about the effect of diversity on the stability of communities (Pennekamp et al. 2018; Steiner et al. 2005). In contrast to stability, however, forecastability is associated with more sources of error resulting from the training of forecast models. Consequently, it is to be expected that variance in the data is larger for forecastability than for response variables such as stability or species abundances.

Although diversity explained only a part of the variation in forecastability of taxa abundances (Supplement Section S6.1), our results nevertheless indicate that diversity tends to increase forecastability in changing environments. From an ecosystem management perspective, our results therefore suggest that diversity conservation could have a positive effect on the accuracy of population forecasts in changing environments. More-diverse communities might be less prone to ecological surprises as environmental conditions change because diversity can promote the resistance of functional groups (Baert *et al*. 2016; Isbell *et al*. 2015). As implied by our findings (see discussion above and Supplement Section S8.1), the loss or rise of functional groups can have large impacts if it concerns species with strong ecosystem-effects. For example, the decline of keystone predators or consumers can have cascading effects on other groups (Amundrud & Srivastava 2016; Estes *et al*. 2011), potentially leading to novel dynamics of many species in the community, and therefore to overall low forecastability. Such species with strong ecosystem-effects might thus be of particular concern in conservation and management, not only because of their effects on ecosystem functioning but potentially also on population forecasts. To confirm that our findings are transferable to a management context, we conducted robustness analyses which showed that treatment effects on forecastability were similar when we back-transformed the forecasts to the original scale of the time series (Supplement Section 7.5). Taken together, diversity maintenance might reduce ecological surprises in changing environments, thereby increasing the forecastability of species abundances.

In contrast to our hypothesis, the forecastability of aggregate properties (i.e. community biomass, oxygen) was largely unaffected by our experimental treatments. Trends of treatment effects went in the same rather than opposite direction as effects on forecastability of species abundances. We had expected that contrasting effects of diversity on the temporal variability of aggregate and non-aggregate properties (Tilman 1996; Xu *et al*. 2021) may translate into contrasting effects on forecastability. However, in our experiment, species richness affected neither the temporal variability of population abundances (Figure 2e) nor that of community biomass or oxygen (Supplement Figure S31). In general, the experimental treatments had similar effects on time series properties irrespective of the level of aggregation of the response variable, possibly explaining the similarity in treatment effects on forecastability. Overall, our results suggest that diversity and environmental change are less influential in affecting the forecastability of aggregate ecosystem properties.

Also in contrast to our expectations, forecastability of community biomass did not differ from average forecastability of species abundances (Supplement Table S12). Forecastability is expected to increase with the level of aggregation because aggregate properties might integrate over environmental variability (Lewis *et al*. 2023). Indeed, community biomass had lower temporal variability than the biomass of individual species (Supplement Figure S32). However, our results suggests that lower temporal variability does not necessarily translate into higher forecastability. Rather, we observed the opposite pattern: time series with higher variability were forecasted better, indicating that higher temporal variability did not reflect more stochastic but more deterministic variation (e.g. abundance changes in response to the light decline or to changes in competitor abundances). At the level of the community, compensatory dynamics among species might have led to reduced variability of community biomass without increasing its forecastability because biomass integrated over primarily deterministic (not stochastic) variation. In conclusion, our results do not support the prospect that forecastability increases by aggregating individual ecosystem components.

Our study illustrates that experiments with microbial organisms are a useful tool to investigate potential drivers of forecastability. Data-driven forecasting methods require long time series (Dietze 2017), which can be collected within a comparatively short time frame for microbial organisms because of their short generation times. Our experimental system is a model of a food web dominated by consumer-resource interactions. Investigating how diversity and environmental change influence forecastability using other systems (e.g. mutualistic networks) and other environmental factors would be a promising avenue of future research to come to a more general understanding about the drivers of forecastability. Likewise, manipulation of different aspects of community structure could shed further light on the potential and limits of ecological forecasting. We manipulated species richness and attempted to keep the number of functional groups constant across bottles. However, since not all species persisted, as is common in such experiments (Steiner *et al*. 2006), our communities differed in the number of functional groups (Supplement Section S9). While the number of trophic levels was the same in all but one bottle, the number of functional groups was positively correlated with median realized richness (Supplement Figure S33) and had similar effects on forecast error as realized richness (Supplement Figure S34). Our experiment therefore does not allow to disentangle whether the diversity effects were due to differences in the number of species or in the number of functional groups. Teasing apart such effects could be a useful next step in future experiments.

Our findings give novel insights into the drivers of forecastability. Previous work has focused on comparing forecastability among different species and ecosystem properties (Lewis *et al*. 2022; Ward *et al*. 2014). For example, observational research showed that small, fast-growing species are less forecastable than large, slow-growing species (Ward *et al*. 2014), and experimental research found that species with fewer (and stronger) interactions are less forecastable than species with more (and weaker) interactions (Daugaard *et al*. 2022). Time series properties have been identified as a further driver of forecastability, with higher time series complexity and stronger non-linearity leading to lower forecastability (Pennekamp *et al*. 2019). Previous work has also explored how transferrable forecasting models are to novel biotic contexts and showed that models are less transferrable for a species that is more impacted by a change in biotic context than for a species that is less impacted (Dumandan *et al*. 2024). Our findings advance our understanding about ecological forecasting by focusing on how properties of the community (i.e. diversity) and of the environment (constant versus changing) affect the forecastability of the species embedded in these communities. We found only weak support for the expectation that higher diversity reduces the forecastability of populations. Whether a longer diversity gradient would alter this finding is an open question. If species in very diverse communities have more (but weaker) interactions, a longer diversity gradient might not necessarily result in lower forecastability, given that more (and weaker) interactions can lead to higher forecastability (Daugaard *et al*. 2022). Future work on forecastability could also explore the effect of other community properties (e.g. number of trophic levels), of other global change drivers, and of the number of drivers (single versus multiple drivers). While our study gave first experimental insights, more work is needed to further improve our understanding about the drivers of forecastability.

In conclusion, our results suggest that diversity does not necessarily reduce the forecastability of ecological systems. Rather, when environmental conditions change, diversity might promote forecastability by buffering against ecological surprises that can result from a decline or rise of functional groups. In contrast, low diversity in combination with environmental change could make ecosystems more unpredictable. Our results therefore caution against assuming high forecastability of ecosystems with low diversity. Forecasting efforts for management or conservation purposes might require particularly finely-resolved time series in ecosystems of low diversity, so as not to miss unexpected changes in ecological dynamics as environmental conditions change. Considering anthropogenic pressures on diversity, reduced forecastability of less-diverse systems may pose an additional challenge to conserve populations, communities, and ecosystems in changing environments.

## Acknowledgements

We thank Fabian Avesani for help in the lab, Bernhard Schmid for discussions on the experimental design, and the Predictive Ecology Group of the University of Zurich for feedback on the manuscript. We thank Frederik Hammes for input on flowcytometry settings and for access to a backup flowcytometer, with help from Jürg Sigrist. This project was part of SNF Project 310030_188431.

## Data availability statement

All data and code used in this study will be made **publicly** available on Zenodo upon publication.

## Contributions

**Conceptualization**: U.D., R.L., O.L.P.; **Data curation**: U.D., R.M.K., R.L., O.L.P.; **Formal analysis**: U.D., R.L.; **Funding acquisition**: O.L.P.; **Investigation**: Y.C., U.D., A.G., M.J., S.J., R.L., S.N., F.P., S.J.v.M, X.Z., D.Z-D.; **Methodology**: U.D., R.M.K., R.L., O.L.P; **Project administration**: R.L., O.L.P.; **Resources**: O.L.P.; **Software**: U.D., R.M.K.; **Supervision**: R.L., O.L.P.; **Validation**: U.D., R.M.K., R.L., F.P., O.L.P.; **Visualization**: U.D.; **Writing – original draft**: U.D., R.L.; **Writing – review & editing:** all authors.

