## Supplementary information for "Mixed evidence for species diversity affecting ecological forecasts in constant versus declining light"

**S9 Other diversity indices 58**

**S10 Versions of R and of the R-packages used 58**

**References 60**

### S1 Implementation of the experiment

#### S1.1 Organisms

We used a pool of 18 species of algae and ciliates in our experiment (Table S1, Figure S1). Based on edibility of algae and prey preferences of ciliates we classified the species into six functional groups: edible algae (two species), inedible algae (5), bacterivorous ciliates (4), omnivorous ciliates (4), mixotrophic ciliates (2), and a predatory ciliate (1). Most species were ordered from culture collections, except for some of the ciliates which were isolated from a pond at the University of Zurich. To remove contaminating algae, all ciliates were cleaned by transferring up to ten individuals through drops of sterile medium. Ciliate cultures established from field isolates were started with one individual to ensure cultures were monospecific.

Stock cultures of algae were cultivated in flasks filled with 100 mL of WC medium and transferred to flasks with fresh medium every three weeks. Stock cultures of ciliates were maintained by serial transfer in cell culture flasks filled with 40 mL of a mixture of 75% protist pellet medium (0.55 g protist pellets $L^{-1}$ Chalkley's medium) and 25% WC medium. Omnivorous and mixotrophic ciliates were fed with algae (*Chlamydomonas* or *Cryptomonas*), the predatory ciliate *Didinium* was fed with *Paramecium caudatum*, and all ciliates were fed with a mixture of three bacteria species (*Bacillus subtilis*, *Serratia fonticola* and *Brevibacillus brevis*).

#### S1.2 Experimental design

In a nine-month experiment, we factorially manipulated light and biodiversity in 30 microcosms. The hypotheses tested with the experiment are illustrated in Figure S2. To manipulate light, we used eight incubators (IPP260plus, Memmert), four per light level. Light was either constantly maintained at 30% of the maximum incubator light intensity or gradually decreased from 30% (~ 22,500 Lux) to 1% (~ 800 Lux). Prior to the light decline, all microcosms were maintained at 30% light for three months to allow slow-growing species to reach the stationary phase, and to produce time series of population and ecosystem variables. We then decreased light for three months with a weekly decrease of 2% relative to the maximum

Table S1: Algae and ciliates used in the experiment.

| Species | Functional group | Origin | Strain number | Prey | Indistinguishable from |
|---|---|---|---|---|---|
| *Chlamydomonas reinhardtii* | Edible algae | SAG[a] | 11-32b | | |
| *Cryptomonas* sp. | Edible algae | SAG | 26.80 | | |
| *Cosmarium botrytis* | Inedible algae | SAG | 136.80 | | |
| *Desmodesmus armatus* | Inedible algae | SAG | 276-4d | | |
| *Monoraphidium obtusum* | Inedible algae | SAG | 55.81 | | |
| *Staurastrum gracile* | Inedible algae | CCAP[b] | 679/3 | | |
| *Staurastrum polytrichum* | Inedible algae | SAG | 20.98 | | |
| *Colpidium striatum* | Bacterivores | Carolina Biological Supply | | Bacteria | |
| *Dexiostoma campylum* | Bacterivores | CCAP | 1611/1 | Bacteria | *Tetrahymena* |
| *Loxocephalus* sp. | Bacterivores | Carolina Biological Supply | | Bacteria | |
| *Tetrahymena thermophila* | Bacterivores | CCAP | 1630/1U | Bacteria | *Dexiostoma* |
| *Coleps* sp. | Omnivores | Pond University Zurich | | *Cryptomonas*, Bacteria | |
| *Paramecium caudatum* | Omnivores | Carolina Biological Supply | | *Chlamydomonas*, Bacteria | |
| *Stylonychia mytilus* | Omnivores | Pond University Zurich | | *Chlamydomonas*, Bacteria | *Stylonychia* sp. |
| *Stylonychia* sp. | Omnivores | Pond University Zurich | | *Chlamydomonas*, Bacteria | *Stylonychia mytilus* |
| *Euplotes daidaleos* | Mixotrophs | Sciento | P229 | *Cryptomonas*, Bacteria | |
| *Paramecium bursaria* | Mixotrophs | SAG | 27.96 | *Chlamydomonas*, Bacteria | |
| *Didinium nasutum* | Predator | Sciento | P220 | *Paramecium caudatum* | |

[a] SAG: Culture Collection of Algae at Göttingen University

[b] CCAP: Culture Collection of Algae and Protozoa

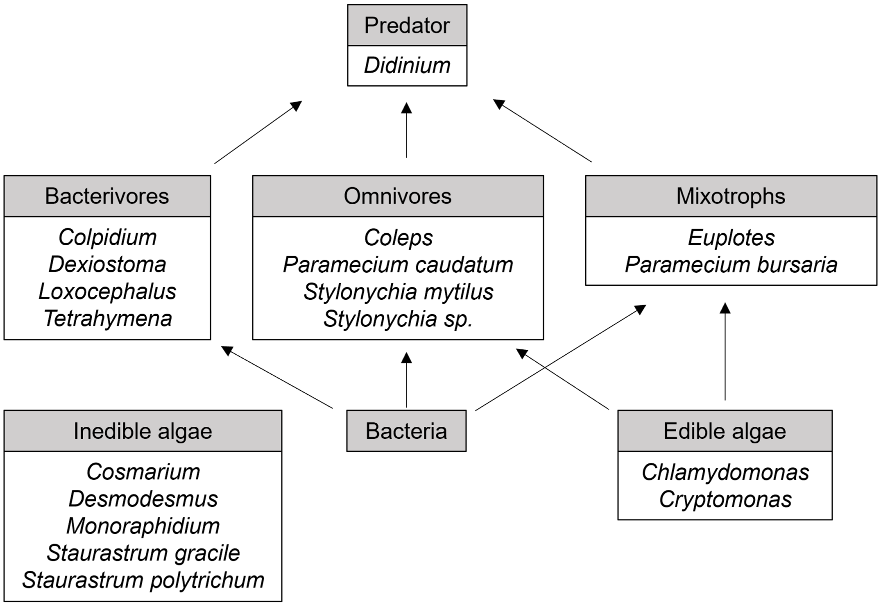

Figure S1: Experimental food web. The boxes display the different functional groups, the arrows denote feeding interactions. The basal level of the food web comprised bacteria, edible algae (i.e. consumable by ciliates), and inedible algae (not consumable by the ciliates). The middle trophic level comprised bacterivores (i.e. ciliates feeding on bacteria), omnivores (ciliates feeding on bacteria and edible algae), and mixotrophs (ciliates containing endosymbiotic algae). The top trophic level consisted of a predator which fed on ciliate consumers. Note that in the experimental communities the predator did not establish or went extinct quickly.

Table S2: Species richness per functional group in the four diversity levels. Numbers in brackets are the number of available species.

| Total (18) | Edible algae (2) | Inedible algae (5) | Bacterivores (4) | Omnivores (4) | Mixotrophs (2) | Predator (1) |
|---|---|---|---|---|---|---|
| 7 | 2 | 1 | 1 | 1 | 1 | 1 |
| 10 | 2 | 2 | 2 | 2 | 2 | 1 |
| 14 | 2 | 3 | 3 | 3 | 2 | 1 |

incubator light, and a final 1% decline. In the last three months of the experiment, all microcosms were maintained constantly at the final light conditions (i.e. 30 and 1%, respectively). Throughout the entire experiment, all incubators were set to a light:dark cycle of 16:8 hours (Figure S3).

We generated three levels of biodiversity by increasing the number of species within the functional groups of inedible algae, bacterivores, and omnivores (one, two, or three species), and in the mixotrophs (one, one, or two species), see Table S2. Because we only had one predator, we could not manipulate diversity at the highest trophic level. We also did not manipulate diversity of edible algae because preliminary tests showed that some consumers grew poorly when communities contained only one of the two edible algae species. The total planned species richness at the three diversity levels was thus 7, 10, and 14 species, respectively (Table S2). An overview of the experimental design is given in Table S3.

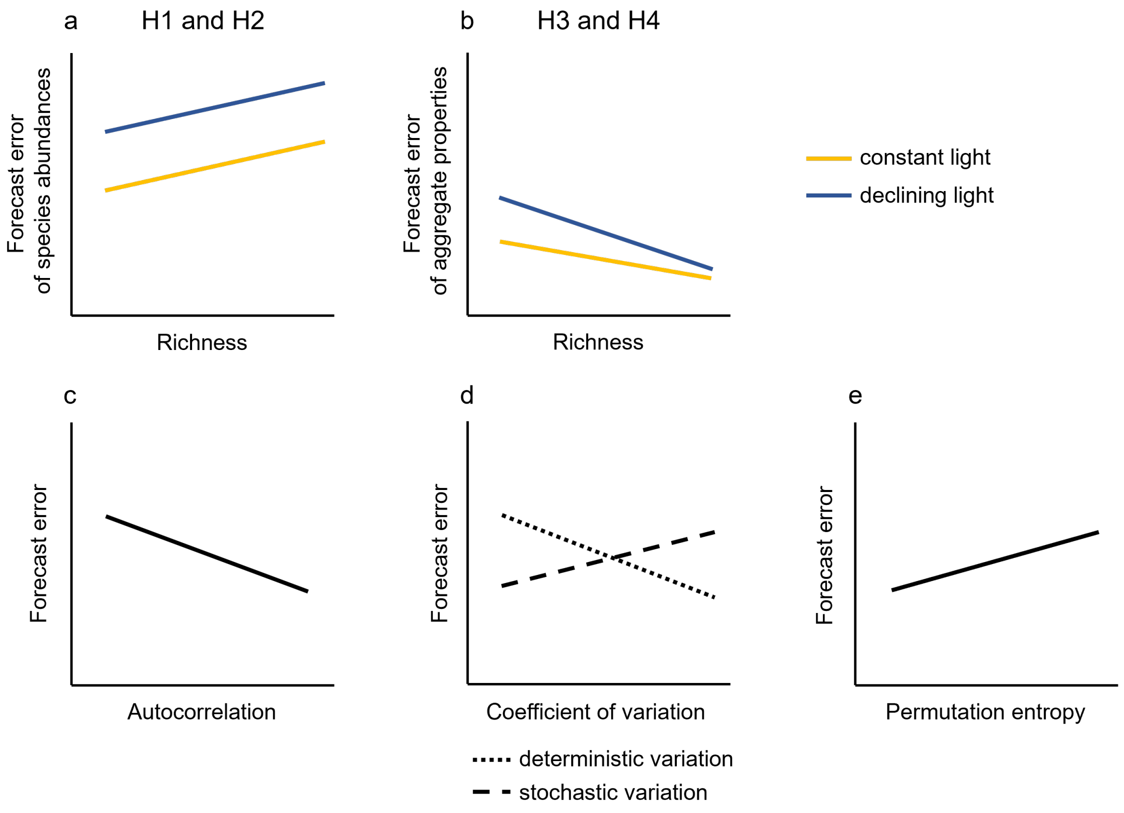

Figure S2: Hypotheses. The figure illustrates the hypotheses regarding the treatment effects on (a) forecast error of species abundances and (b) forecast error of aggregate ecosytem properties, and hypotheses about the relationship between forecast error and time series properties (c-e). (a) We expected that higher diversity leads to higher forecast error of species abundances (Hypothesis 1, H1), and historical change in environmental conditions (i.e. declining light) increases forecast error because forecasts are made for less-often observed conditions. (b) We expected that forecast error of aggregate ecosystem properties decreases with diversity, in particular in changing environments (H3), and (a vs. b) forecast error of aggregate ecosystem properties is lower than forecast error of species abundances (H4). To investigate possible reasons for differences in forecast error, we analyzed how the forecast error of species abundances was related with three time series properties. We expected that (c) higher autocorrelation would be associated with lower forecast error, (d) a higher coefficient of variation would be associated with higher forecast error if higher variation was due to more stochastic variation, but with lower forecast error if higher variation was due to more deterministic variation, and (e) higher permutation entropy would be associated with higher forecast error.

Table S3: The summarized experimental design.

| Bottle | Composition | Planned richness | Light conditions | Incubator |
|---|---|---|---|---|
| b__01 | c__08 | 10 | Constant | A |
| b__02 | c__02 | 7 | Constant | A |
| b__03 | c__11 | 14 | Constant | A |
| b__04 | c__05 | 7 | Constant | A |
| b__05 | c__13 | 14 | Decreasing | B |
| b__06 | c__11 | 14 | Decreasing | B |
| b__07 | c__06 | 10 | Decreasing | B |
| b__08 | c__01 | 7 | Decreasing | B |
| b__09 | c__02 | 7 | Decreasing | C |
| b__10 | c__07 | 10 | Decreasing | C |
| b__11 | c__08 | 10 | Decreasing | C |
| b__12 | c__14 | 14 | Decreasing | C |
| b__13 | c__06 | 10 | Constant | D |
| b__14 | c__03 | 7 | Constant | D |
| b__15 | c__14 | 14 | Constant | D |
| b__16 | c__15 | 14 | Constant | D |
| b__17 | c__05 | 7 | Decreasing | E |
| b__18 | c__04 | 7 | Decreasing | E |
| b__19 | c__12 | 14 | Decreasing | E |
| b__20 | c__09 | 10 | Decreasing | E |
| b__21 | c__07 | 10 | Constant | F |
| b__22 | c__10 | 10 | Constant | F |
| b__23 | c__04 | 7 | Constant | F |
| b__24 | c__13 | 14 | Constant | F |
| b__25 | c__03 | 7 | Decreasing | G |
| b__26 | c__15 | 14 | Decreasing | G |
| b__27 | c__10 | 10 | Decreasing | G |
| b__28 | c__09 | 10 | Constant | H |
| b__29 | c__12 | 14 | Constant | H |
| b__30 | c__01 | 7 | Constant | H |

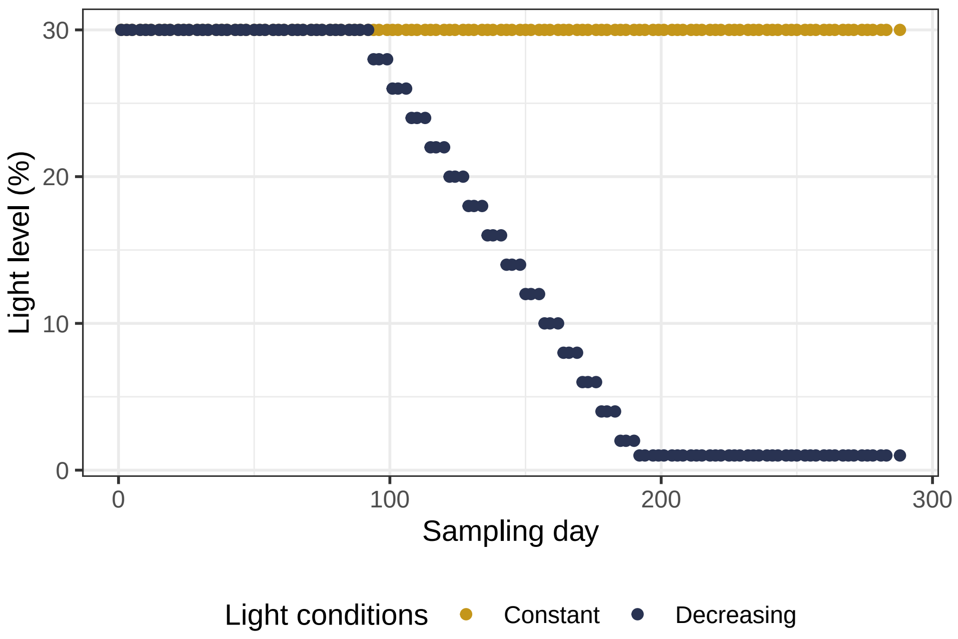

Figure S3: Light levels over the course of the experiment. After 3 months of constant environmental conditions, bottles either remained at the initial light level (30% light) or experienced a gradual decline in light for three months. Then, bottles remained three more months at the final light levels (30 or 1% light intensity). Each point represents a sampling day.

#### S1.3 Selection of compositions

To disentangle effects of species richness and species identity/composition, we assembled five different compositions at each diversity level such that compositions were as dissimilar as possible and species occurred with similar frequency both within and across diversity levels. An additional criterion for community assembly was that species could not co-occur if they were not distinguishable with our methods of quantification (Table S1). We first constructed all species combinations that met the criteria regarding distinguishability of species and had appropriate species numbers in each functional group (Table S2). From this pool of possible compositions, we randomly selected five compositions for each diversity level. To quantify dissimilarity of compositions within diversity levels, we calculated the mean pairwise Sorensen distance of compositions. However, a high mean pairwise Sorensen distance does not necessarily preclude that pairs of communities are very similar. For example, when of four communities two each are identical in composition but do not share species with the other two communities, mean pairwise Sorensen distance is high because two low distances are outweighed by four high distances. To quantify such effects, we also calculated the standard deviation of the pairwise Sorensen distances, with low standard deviation indicating absence of highly similar pairs of compositions.

As a further measure of dissimilarity, we checked if the five compositions were unique within functional groups. This was a useful criterion for the groups of inedible algae, bacterivores, and omnivores when species richness was 10. In the mixotrophs, however, the number of available species was too low to assemble five unique compositions. Similarly, at the lowest and highest diversity level, we could not avoid duplicates or triplicates of compositions within bacterivores, omnivores, and mixotrophs because the number of available species was not high enough. However, by maximizing dissimilarity (see below) we ensured that compositions were unique across functional groups.

To calculate if species occurred with similar frequency within diversity levels, we first computed the expected occurrence of each species under the assumption of equal frequency (i.e. sum of occurrences of species of functional group $i$ at a given diversity level / number of available species of functional group $i$). We then quantified the average per species deviation from equal occurrence by calculating the mean absolute difference between observed and expected occurrence. As a second metric we calculated deviation from equal occurrence per functional group to give equal weight to functional groups rather than to species. We first calculated the deviation from equal occurrence per functional group by calculating the sum of squared differences between observed and expected occurrences in functional group $i$. We then averaged over all functional groups after taking the square root of the per functional group deviation and dividing by the number of species in the respective functional group.

To maximize dissimilarity of compositions and minimize deviation from equal occurrence of species, we repeated the random draws of compositions 300,000 times for each diversity level, and then selected the sets of compositions that had maximum pairwise Sorensen distance. These sets of compositions simultaneously had minimum deviation from equal species occurrence. We then removed sets of compositions with species richness 10 unless all five compositions were unique within the groups of inedible algae, bacterivores, and omnivores. Of the remaining sets of compositions, we chose the ones with minimum standard deviation of the pairwise Sorensen distances. Except for one diversity level, this procedure resulted in more than one possible set of compositions.

Finally, we maximized dissimilarity of compositions and minimized deviation from equal species occurrence across diversity levels. From the sets of compositions with maximum dissimilarity within diversity levels, we randomly selected one set of compositions from each diversity level. For the resulting set of 15 communities, we again calculated the four metrics described above (mean and standard deviation of pairwise Sorensen distances, average per species deviation from equal occurrence, average per functional group deviation from equal occurrence). We repeated the random selection of 15 communities 300,000 times and then chose the 15-community sets with maximum pairwise Sorensen distance; these sets also had lowest deviation from equal species occurrence. Of these sets of combinations we selected those with lowest standard deviation of the pairwise Sorensen distances. From the remaining sets of combinations, we randomly chose one. The final community compositions that we used in the experiment are listed in Table S4.

Table S4: The 15 community compositions used in the experiment. A 1 indicates that a species was used in a specific composition, while a 0 indicates that is was not.

| | Community composition | | | | | | | | | | | | | | |
|---|---|---|---|---|---|---|---|---|---|---|---|---|---|---|---|
| Species | c_01 | c_02 | c_03 | c_04 | c_05 | c_06 | c_07 | c_08 | c_09 | c_10 | c_11 | c_12 | c_13 | c_14 | c_15 |
| *Chlamydomonas reinhardtii* | 1 | 1 | 1 | 1 | 1 | 1 | 1 | 1 | 1 | 1 | 1 | 1 | 1 | 1 | 1 |
| *Cryptomonas* sp. | 1 | 1 | 1 | 1 | 1 | 1 | 1 | 1 | 1 | 1 | 1 | 1 | 1 | 1 | 1 |
| *Monoraphidium obtusum* | 0 | 0 | 1 | 0 | 0 | 1 | 0 | 0 | 1 | 0 | 0 | 1 | 1 | 0 | 1 |
| *Cosmarium botrytis* | 0 | 0 | 0 | 1 | 0 | 0 | 0 | 1 | 0 | 1 | 1 | 1 | 1 | 0 | 0 |
| *Staurastrum gracile* | 0 | 1 | 0 | 0 | 0 | 0 | 1 | 0 | 1 | 0 | 1 | 1 | 0 | 1 | 0 |
| *Staurastrum polytrichum* | 1 | 0 | 0 | 0 | 0 | 0 | 1 | 0 | 0 | 1 | 0 | 0 | 1 | 1 | 1 |
| *Desmodesmus armatus* | 0 | 0 | 0 | 0 | 1 | 1 | 0 | 1 | 0 | 0 | 1 | 0 | 0 | 1 | 1 |
| *Tetrahymena thermophila* | 1 | 1 | 0 | 0 | 0 | 0 | 0 | 1 | 1 | 0 | 1 | 0 | 1 | 0 | 0 |
| *Colpidium striatum* | 0 | 0 | 0 | 0 | 1 | 0 | 1 | 0 | 1 | 1 | 1 | 1 | 1 | 1 | 1 |
| *Loxocephalus* sp. | 0 | 0 | 1 | 0 | 0 | 1 | 1 | 1 | 0 | 0 | 1 | 1 | 1 | 1 | 1 |
| *Dexiostoma campylum* | 0 | 0 | 0 | 1 | 0 | 1 | 0 | 0 | 0 | 1 | 0 | 1 | 0 | 1 | 1 |
| *Paramecium caudatum* | 0 | 1 | 0 | 0 | 0 | 1 | 1 | 1 | 0 | 0 | 1 | 1 | 1 | 1 | 1 |
| *Stylonychia mytilus* | 1 | 0 | 0 | 1 | 0 | 1 | 0 | 0 | 1 | 0 | 0 | 0 | 1 | 1 | 0 |
| *Stylonychia* sp. | 0 | 0 | 1 | 0 | 0 | 0 | 1 | 0 | 0 | 1 | 1 | 1 | 0 | 0 | 1 |
| *Coleps* sp. | 0 | 0 | 0 | 0 | 1 | 0 | 0 | 1 | 1 | 1 | 1 | 1 | 1 | 1 | 1 |
| *Paramecium bursaria* | 1 | 0 | 0 | 0 | 1 | 0 | 1 | 1 | 1 | 0 | 1 | 1 | 1 | 1 | 1 |
| *Euplotes daidaleos* | 0 | 1 | 1 | 1 | 0 | 1 | 0 | 0 | 0 | 1 | 1 | 1 | 1 | 1 | 1 |
| *Didinium nasutum* | 1 | 1 | 1 | 1 | 1 | 1 | 1 | 1 | 1 | 1 | 1 | 1 | 1 | 1 | 1 |
| **Planned richness** | **7** | **7** | **7** | **7** | **7** | **10** | **10** | **10** | **10** | **10** | **14** | **14** | **14** | **14** | **14** |

#### S1.4 Assignment of treatments to microcosms and incubators

To assign treatments to microcosms and incubators, we grouped incubators into pairs based on their position in the lab, and randomly assigned one incubator per pair to one of the two light levels. We then assigned the 15 communities to the four incubators of a given light level by randomly selecting one community per diversity level for each incubator, and then randomly distributing the remaining three communities (i.e. one per diversity level) over incubators. Finally, we randomly assigned the (three or) four communities of a given incubator to microcosms. To reduce effects of inhomogeneous light distribution in the incubators, we moved the microcosms every other week to a new position in their respective incubator.

#### S1.5 Inoculation

At the start of the experiment, we filled 30 2L-bottles (Duran) with 670 mL of medium, added 20 autoclaved wheat seeds for slow release of nutrients, and a magnetic stirrer for homogenization during samplings. During the four-week inoculation phase, we added a total of 330 mL of stock cultures, which were growing at high densities. In week one, we added 40 mL of prey (10 mL of a mixture of the bacteria *Bacillus subtilis*, *Serratia fonticola* and *Brevibacillus brevis*, 10 mL of *Chlamydomonas*, and 20 mL *Cryptomonas*), and 40 mL of inedible algae. The volume assigned to inedible algae was split among one to three species depending on the diversity level (i.e. 40 mL per species when diversity was one, 20 mL per species when diversity was two, and 13.3. mL per species when diversity was three). In week two, we repeated the inoculation of bacteria and algae to increase the probability of successful establishment of all species. In week two, we added the consumers, with 80 mL of volume split among three to eight ciliate species depending on the diversity level. Inoculation of consumers was repeated a week later. Finally, we added 10 individuals of the predator *Didinium* to each bottle in week three and 20 individuals of *Didinium* in week four. The lower number of *Didinium* added at the first *Didinium*-inoculation was due to unexpectedly low *Didinium* densities in the stock cultures. After completion of inoculation, the composition of the medium in the bottles was 75% protist pellet medium and 25% WC medium.

During the inoculation phase, we mixed and homogenized the bacterial communities associated with the algae and ciliate stock cultures to avoid that differences among our experimental communities were due to differences in associated bacteria rather than to differences in protist composition. To this end, we prepared a mix of all inedible algae at their first inoculation, and filtered the mixture through a filter with 1.2 $\mu$m mesh size (Acrodisc Syringe Filters, Pall) to remove all algae but maintain the majority of associated bacteria in the filtrate. We then added 5 mL of the filtrate to each bottle. We repeated this approach with the ciliate species at their first inoculation.

##### S1.6 Immigration

To allow species to re-colonize bottles if they had gone extinct, we re-immigrated the species from the stock cultures every three weeks. At these immigration events, we added 10 individuals of each species that was part of the community of a given bottle. Soon after the start of the experiment, two species went extinct in the stock cultures (*Stylonychia mytilus* and *Didinium nasutum*); we therefore could not re-immigrate these two species.

#### S2 Sampling of the experiment

##### S2.1 Sampling

After the four-week inoculation phase, we sampled the 30 bottles three times per week (Monday, Wednesday, Friday) for nine months (41 weeks). This resulted in 123 sampling days. We were able to always maintain this sampling schedule except for the last sampling event which had to be postponed from a Friday to the following Monday.

On each sampling day, we removed 50 mL from each bottle after homogenization on a magnetic stirrer, and then added 50 mL of fresh medium to the bottle. Samples were taken under sterile conditions using autoclaved 50-mL glass pipettes.

##### S2.2 Ciliates

We quantified ciliate abundances with video microscopy. We took six videos per sample, three videos at 16x magnification, three at 25x magnification. To this end, we used two identical video-microscopy set-ups. Each consisted of a dissecting microscopes (Leica M250 C) with dark-field illumination, each equipped with a Hamamatsu C11440 camera. From each sample, we set up two counting chambers filled with 0.65 mL of sample and made three videos per chamber. We used the software HCImage Live to take videos of five seconds length and 25 frames per second, with field delay set to 40 ms, exposure to 10 ms, and field depth to 8 bit. We used the R-package `bemovi` (Pennekamp, Schtickzelle, and Petchey 2015), to track moving particles and quantify morphological and movement traits. We extended this package slightly (package name: `LEEF.measurement.bemovi`) so that it is cloud-compatible (Krug 2024). The three small ciliate species (i.e. *Tetrahymena*, *Dexiostoma* and *Loxocephalus*) were only tracked at the higher magnification.

Because we expected the predator *Didinium nasutum* to occur in low abundances, we counted *Didinium* manually in up to 3 mL of sample using a dissecting microscope. However, *Didinium* established only in few bottles and soon went extinct both in the experiment and in the stock cultures. We continued checking the samples under a dissecting microscope but with reduced effort (1 mL volume, once per week).

#### S2.3 Algae

We quantified algae abundances and traits with a FlowCAM (VS1; Yokogawa Fluid Imaging Technologies, Scarborough, Maine, USA). This instrument counts and images the particles in a sample and reports 40 morphological properties (i.e. traits) of each particle. The sample was pumped with a flowrate of 0.3 mL per minute through a flowcell of 100 µm diameter and 700 $\mu$m width (pump: 5-mL C70 syringe; flowcell: FC100-FV). The sample was imaged using an objective with 10x magnification and a camera (Sony SX90CR), an auto-image rate of 20 frames per second, and a run time of 1 minute. We used the software Visual Spreadsheet (version 3.4.11) to count and analyze the particles in the sample. The segmentation threshold was set to 20 for dark pixels, i.e pixels were considered part of particles if they were at least by a value of 20 darker than the corresponding pixels of the calibration image.

The samples were measured alive within 4-5 hours of sampling. After a few sampling days, some bottles developed large clumps that clogged the flowcell. From the $14^{\text{th}}$ sampling onwards, we therefore filtered all samples through filters (pluriSelect) of 200 $\mu$m mesh size to remove large clumps. On one sampling day, we measured all samples unfiltered and filtered through 200 $\mu$m and found no difference in species abundances (Figure S4).

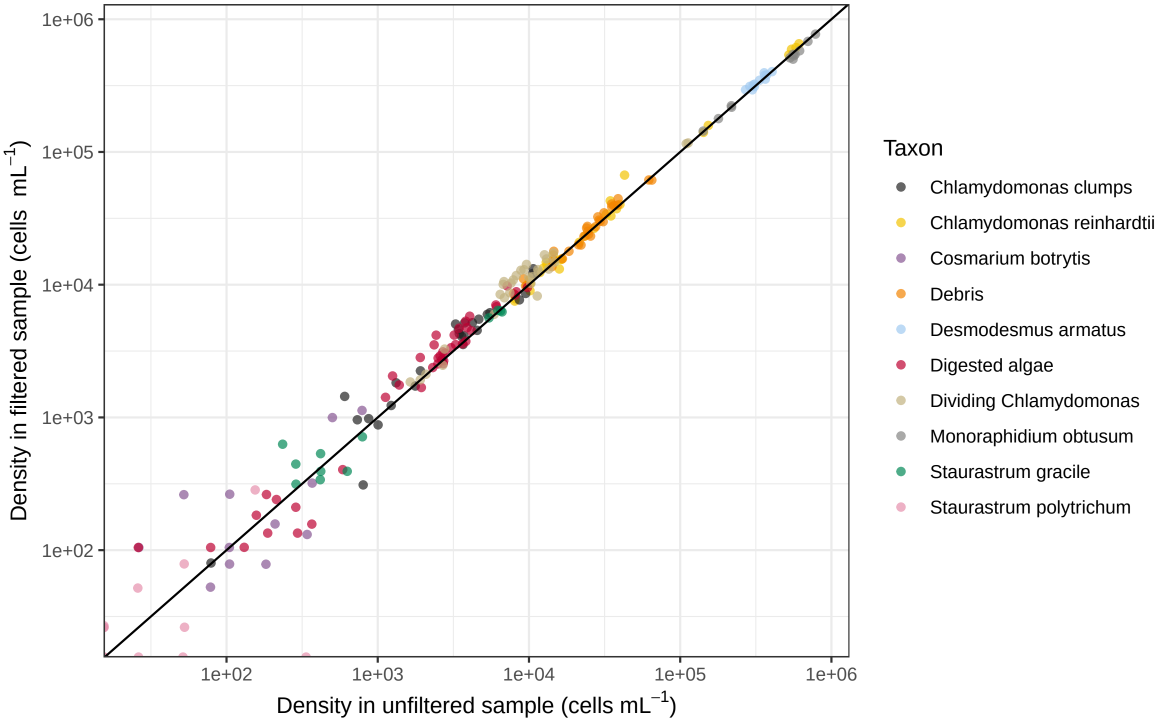

Figure S4: Comparison of filtered and unfiltered algae samples. On one sampling day, we made two flowcam measurements for each of the 30 samples, one from an unfiltered sample and one from a sample that had been filtered through a 200 $\mu$m mesh. The black line is the identity line.

##### S2.4 Bacteria

To measure abundances of bacteria, we used a flowcytometer (BD Accuri C6 Plus; BD Biosciences) equipped with an autosampler. At the majority of sampling days, we measured three replicates per sample, two with a 10-fold dilution and one with a 30-fold dilution. We diluted the samples in a 0.005 M EDTA solution, stained the samples with SYBR Green I (Invitrogen), and incubated the samples for 20 minutes at 37°C. Samples were set up in 96-well plates with 150 $\mu$l of diluted sample per well. In addition, we measured three replicate blanks of ultrapure water and of medium, respectively. We measured 35 $\mu$l of the diluted samples with a flowrate of 35 $\mu$l min$^{-1}$ and a threshold of 800 on the green fluorescence channel (FL1).

Some of the flowcytometry measurements deviated from this standard procedure. Specifically, from day 197 to 220, the flowcytometer normally used was broken. In this period, we either measured the samples on a different machine of the same type (BD Accuri C6; six sampling dates) or did not measure the samples (five sampling dates). We skipped never more than two consecutive samplings. At some samplings, we measured only one or two replicates (e.g. when the autoloader was broken). After 200 days, we started to pre-filter the samples through filters with 200 $\mu$m mesh size (pluriSelect) to avoid clogging of the flowcytometer. A comparison of unfiltered and filtered samples indicated that filtration did not alter bacterial abundances (Figure S5).

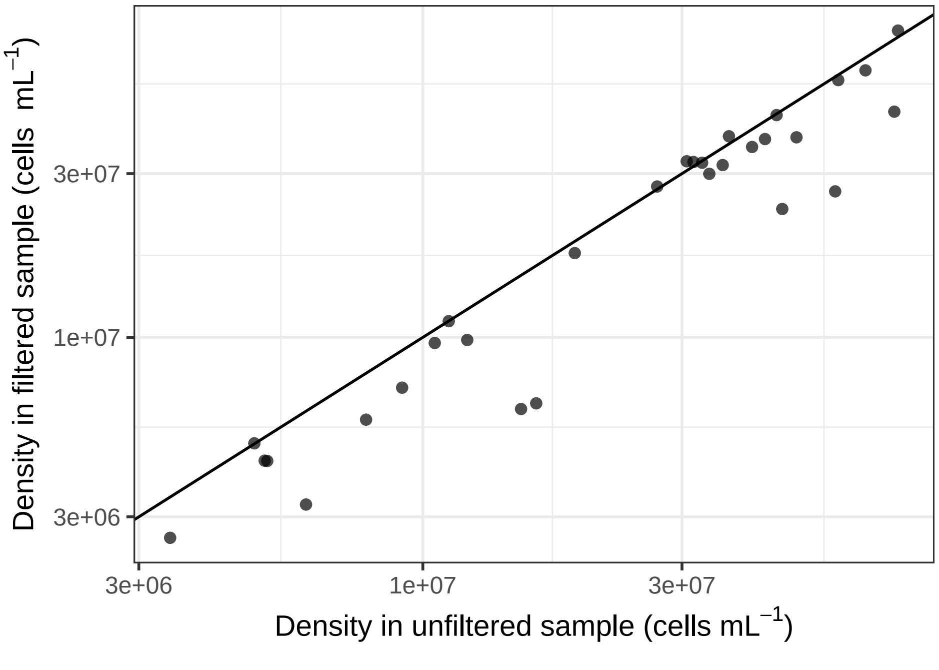

Figure S5: Comparison of filtered and unfiltered bacteria samples. On one sampling day, we made two flowcytometer measurements for each of the 30 samples: one from an unfiltered sample and one from a sample that had been filtered through 200 $\mu$m. The black line is the identity line.

To differentiate between bacteria, noise, and algae, we used a gate on the plot of green fluorescence (FL1) versus red fluorescence (FL3). We applied the same gate coordinates to all samples, except when we used a different flowcytometer (BD Accuri C6; six sampling dates); in that case, the gate started at a slightly higher FL-1 fluorescence (2500 rather than 2000) to avoid incorporation of background noise. We used the R-package `flowCore` (Ellis et al. 2023) to apply the gate to the flowcytometry samples. The general gating strategy is shown in Figure S6.

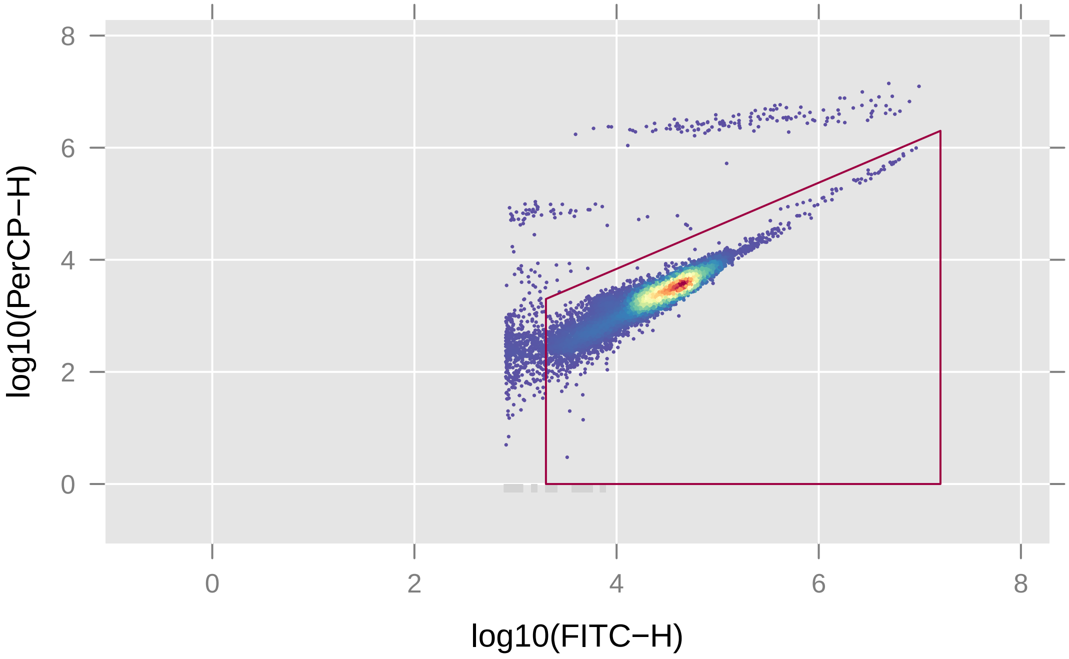

Figure S6: Example of a sample measured on the flowcytometer to illustrate the gating strategy. Bacteria were gated on the plot of FL-1 (i.e. FITC-H) versus FL-3 (i.e. PerCP-H). In this example, the bacteria population contained 178,063 cells. A random subsample of 10% of the measured particles is displayed to avoid over-plotting.

#### S2.5 Abiotic variables

To measure dissolved oxygen, we attached two optical oxygen sensors (SP-PSt3-YAU, PreSens Precision Sensing GmbH, Germany) on the inner wall of each bottle, one at the level of 400 mL and one at the level of 900 mL, and measured % oxygen with a hand-held oxygen meter (Fibox-4; PreSens Precision Sensing GmbH, Germany) equipped with a polymer optical fiber.

To measure dissolved carbon and nitrogen, we used a TOC/TN analyzer (Formacs HT). Samples were stored at -20°C prior to processing. After thawing, samples were centrifuged (4,000 rpm; 5 minutes) and the supernatant was analyzed with the TOC/TN analyzer. Dissolved organic carbon (DOC) was calculated by subtracting dissolved inorganic carbon from total dissolved carbon.

#### S2.6 Taxa classification and abundance estimation

After data collection, we identified the tracked particles (algae and ciliates) based on their movement (videos) and morphological (videos and FlowCAM) traits. For this, we trained composition-specific automated classifiers, separately for the algae and for the ciliates. The classifiers were support vector machines (SVM). The preparation of the data used to train the classifiers involved several steps. First, we collected trait data from monocultures cultivated at different light levels (30, 18, 10, 6 and 1% light). In addition, the training data also included several additional classes of other potentially present particles. For the FlowCAM, these classes were "Air bubbles", "*Chlamydomonas* clumps", "*Coleps*", "*Colpidium*", "*Colpidium* vacuoles", "Debris", "*Desmodesmus* clumps", *Dexiostoma*, "Digested algae", "Dividing *Chlamydomonas*", "*Loxocephalus*", "Other ciliate", "Small cells" (i.e. cells present in all bottles but that we did not further identify), "Small unidentified" (small, non-cell particles, probably debris), and "*Tetrahymena*". Trait data on these additional classes was collected from the experimental communities after manually categorizing particles into one of these classes. For the videos, the additional classes were "*Cryptomonas*" and "Debris and other things". Further, because the species *Stylonychia mytilus* went extinct everywhere immediately it was not included as a class in the classifiers.

Next, to ensure that the tracked particles were representative of the respective species we cleaned the training data. Specifically, for the ciliates we only kept particles that were tracked for more than 10 frames, and we also excluded (almost) static particles. Further, in all cases we excluded particles that showed clear outlier values in several of their traits. As not all species are equally abundant, to improve classifier performance we then sub-sampled the training data so that we had a more balanced amount of data for each species from each of the data sources (i.e. different light levels). In a final step, we then split the prepared data into the actual training data and into the evaluation data, and then trained the classifiers, using most of the recorded trait variables. For this we used the R-package `e1071` (Meyer et al. 2021) and the function `tune()` which we used to do a grid search for the hyperparameters of the chosen kernel (radial). Note that we did not train different classifiers for different light conditions. Instead, we aimed at producing general classifiers capable of distinguishing the different species across the entire light gradient. In this way, the resulting species abundance time series are smoother (i.e. without artificial changes introduced by using different classifiers) and the ranges of the recorded trait values of the species were not limited by the classifiers (because of their generality).

We tested the classifiers based on the evaluation data and based on several performance metrics: the positive predictive value (PPV), the negative predictive value, the sensitivity and the specificity. However, among these metrics the most important is the PPV. We aimed at having PPV values of at least 90% for all classes, but accepted lower values (but above 80%) in rare cases. Further, for the FlowCAM we manually annotated data from the experiment itself (from all bottles) and then compared the manual classification with the automated classification (same metrics as before). If the classification was not satisfactory, we improved the training data (with additional monoculture data and the annotated experiment data) and repeated the process until we were satisfied with the classifier performance.

In the case of the videos, manual annotation is more difficult. Instead, we assessed all automated classifiers (i.e. also the algae classifiers) with visual comparisons/classifications and manually improved the classification when necessary, which was the case in one instance: The density estimation of *Staurastrum gracile* was inadequate as particles in most cases belonging to the classes "Debris" and "*Chlamydomonas* clumps" were classified as *Staurastrum gracile* with some frequency. Normally, the classification works such that a particle is assigned to the class to which it belongs with the highest probability. In the case of *Staurastrum gracile* only, we found that the additional constraint that this probability had to be at least 90% solved the issue (we excluded particles assigned to *Staurastrum gracile* with lower probabilities). Further, note that we did additional quality controls (not limited to classification quality) that are described in the next section (section "Quality controls and time series processing").

In a last step, we then used the identified individuals to estimate the densities of the different taxa, based on the imaged volume. For the videos, this entailed averaging the counted particles over the three recorded videos that we took per bottle, magnification and sampling day.

### S3 Quality controls and time series processing

#### S3.1 Quality controls and missing data percentages

To ensure that the time series were of the highest quality possible we did several controls which are described in this section. The first control was of video quality, as over the course of the experiment we took more than 20000 videos. For this, we first randomly selected 1000 videos across the entire experiment and across all bottles at each of the two magnifications used. We then inspected every selected video for possible problems. In this way, at magnification 25x we found that 986 videos (98.6%) showed no problems, 10 (1.0%) were faulty because of shaking (making it impossible to track particles), 3 (0.3%) were faulty because of moving background (i.e. the liquid in the counting chamber was moving, interfering with tracking of particles) and 1 was faulty (0.1%) because of other reasons (e.g. the counting chamber was not properly captured, or something else interfered with the video). For the 16x magnification, the respective percentages were 99.2, 0.4, 0.1 and 0.3%. With the vast majority of videos passing the quality control, we concluded that the videos in general were adequate to be used.

The next quality control involved excluding time series of extinct species. At regular intervals throughout the experiment, we did manual occurrence checks in all bottles. The automated classification is, inevitably, imperfect and even if a species has gone extinct as long as the corresponding class is present in the classifier, particles can be, and will be identified as this species at a low frequency. By comparing the recorded time series with the manual checks we excluded the time series of species that we knew were never present in the bottles (i.e. they went extinct right at the start of the experiment), while we kept time series of species that went extinct or re-established themselves later into the experiment. We also excluded time series of species that never established themselves and were consistently below the detection limit, which resulted in abundance time series with many missing values (this was only the case for *Staurastrum polytrichum* in bottles 5, 12, 15, 16, 24 and 26).

Specifically, the following are the time series that we excluded based on the above criteria. *Cryptomonas*: all bottles. *Colpidium*: bottles 3, 4, 5, 6, 10, 12, 15, 16, 19, 21, 22, 24, 26, 28, 29. *Dexiostoma*: bottles 7, 12, 13, 15, 16, 19, 22, 26, 27, 29. *Euplotes*: bottles 2, 3, 5, 6, 7, 9, 12, 13, 14, 15, 16, 19, 22, 23, 24, 25, 26, 27, 29. *Loxocephalus*: bottles 1, 3, 5, 6, 7, 10, 11, 12, 13, 14, 15, 16, 19, 21, 24, 26, 29. *Staurastrum gracile*: bottles 3 and 15. *Staurastrum polytrichum*: bottles 5, 12, 15, 16, 24, 26. *Stylonychia* sp.: bottles 3, 6, 16, 22, 26. *Tetrahymena* all bottles.

Finally, measurements on the TOC/TN analyzer were influenced by measurement position. Specifically, samples measured later in a run had higher values than samples measured earlier in the run. However, as we found the relation between sampling order and measured values to be linear it proved to be sufficient to remove the linear trend (also note that we did not forecast the carbon and nitrogen concentrations).

Regarding missing data, no videos are available for sampling day 20220622 at the 25x magnification. The 16x videos were available, however, and allowed sampling of all species except for the small ciliates, i.e. *Tetrahymena*, *Dexiostoma* and *Loxocephalus*, which we only counted at magnification 25x. Further, and as mentioned in Section S2.4, the flowcytometer was not always available, which resulted in 4.8% of the bacteria density time points to be missing. Table S5 further lists the missing time points percentages for all measurement methods, bottles and experimental treatments. As can be seen, only very few time points were missing (1.67% for the oxygen, below 1% for the other measurements), with missing data generally being caused either by problems with the measurement instrument or by sampling mistakes. The missing data was imputed (see the next subsection for more details).

#### S3.2 The recorded time series and their processing

The recorded time series as recorded are depicted here as follows. The biotic times series are shown in Figure S7 (forecast targets) and Figure S8 (not forecast targets). Further, Figure S9 depicts the time series of community biomass (i.e. the total biomass in the bottles), Figure S10 the oxygen concentration time series and Figure S11 the time series of the water chemistry (dissolved carbon and nitrogen concentrations).

We processed the time series so that the various forecast methods were appropriate to be used (i.e. we made the time series stationary). The next few figures show the time series after their processing (i.e. the transformed time series) if we used them as forecast targets. The biotic time series are shown in Figure S12, the community biomass time series in Figure S13 and the mean oxygen time series in Figure S14.

Table S5: Percentage of missing data for different sampling methods and grouping variables.

| | Percentage of missing data for different sampling methods | | | | |
|---|---|---|---|---|---|
| Grouping | FlowCam | Flowcytometer | Video | Oxygen | TOC/TN analyzer |
| Bottle 01 | 0.81 | 4.84 | 0.81 | 2.42 | 0.81 |
| Bottle 02 | 0.81 | 4.84 | 0.81 | 1.61 | 0.81 |
| Bottle 03 | 0.81 | 4.84 | 0.81 | 1.61 | 0.81 |
| Bottle 04 | 0.81 | 4.84 | 0.81 | 1.61 | 0.81 |
| Bottle 05 | 0.81 | 4.84 | 0.81 | 1.61 | 0.81 |
| Bottle 06 | 0.81 | 4.84 | 0.81 | 1.61 | 0.81 |
| Bottle 07 | 0.81 | 4.84 | 0.81 | 1.61 | 1.21 |
| Bottle 08 | 0.81 | 4.84 | 0.81 | 1.61 | 0.81 |
| Bottle 09 | 0.81 | 4.84 | 0.81 | 1.61 | 1.01 |
| Bottle 10 | 0.81 | 4.84 | 0.81 | 1.61 | 1.01 |
| Bottle 11 | 0.81 | 4.84 | 0.81 | 1.61 | 1.01 |
| Bottle 12 | 0.81 | 4.84 | 0.81 | 1.61 | 1.01 |
| Bottle 13 | 0.81 | 4.84 | 0.81 | 1.61 | 1.01 |
| Bottle 14 | 0.81 | 4.84 | 0.81 | 1.61 | 1.41 |
| Bottle 15 | 0.81 | 4.84 | 0.81 | 1.61 | 0.81 |
| Bottle 16 | 0.81 | 4.84 | 0.81 | 1.61 | 0.81 |
| Bottle 17 | 0.81 | 4.84 | 0.81 | 1.61 | 1.21 |
| Bottle 18 | 0.81 | 4.84 | 1.21 | 1.61 | 0.81 |
| Bottle 19 | 0.81 | 4.84 | 0.81 | 1.61 | 0.81 |
| Bottle 20 | 0.81 | 4.84 | 0.81 | 1.61 | 0.81 |
| Bottle 21 | 0.81 | 4.84 | 0.81 | 1.61 | 0.81 |
| Bottle 22 | 0.81 | 4.84 | 0.81 | 1.61 | 0.81 |
| Bottle 23 | 0.81 | 4.84 | 1.61 | 1.61 | 0.81 |
| Bottle 24 | 0.81 | 4.84 | 0.81 | 1.61 | 0.81 |
| Bottle 25 | 0.81 | 4.84 | 1.21 | 1.61 | 0.81 |
| Bottle 26 | 0.81 | 4.84 | 0.81 | 1.61 | 0.81 |
| Bottle 27 | 0.81 | 4.84 | 0.81 | 1.61 | 1.21 |
| Bottle 28 | 0.81 | 4.84 | 0.81 | 1.61 | 1.01 |
| Bottle 29 | 0.81 | 4.84 | 0.81 | 2.42 | 1.01 |
| Bottle 30 | 0.81 | 4.84 | 0.81 | 1.61 | 1.01 |
| Constant light | 0.81 | 4.84 | 0.83 | 1.72 | 0.90 |
| Decreasing light | 0.81 | 4.84 | 0.85 | 1.61 | 0.94 |
| Planned richness: 07 | 0.81 | 4.84 | 0.97 | 1.61 | 0.95 |
| Planned richness: 10 | 0.81 | 4.84 | 0.81 | 1.69 | 0.97 |
| Planned richness: 14 | 0.81 | 4.84 | 0.81 | 1.69 | 0.85 |
| **Experiment** | **0.81** | **4.84** | **0.84** | **1.67** | **0.92** |

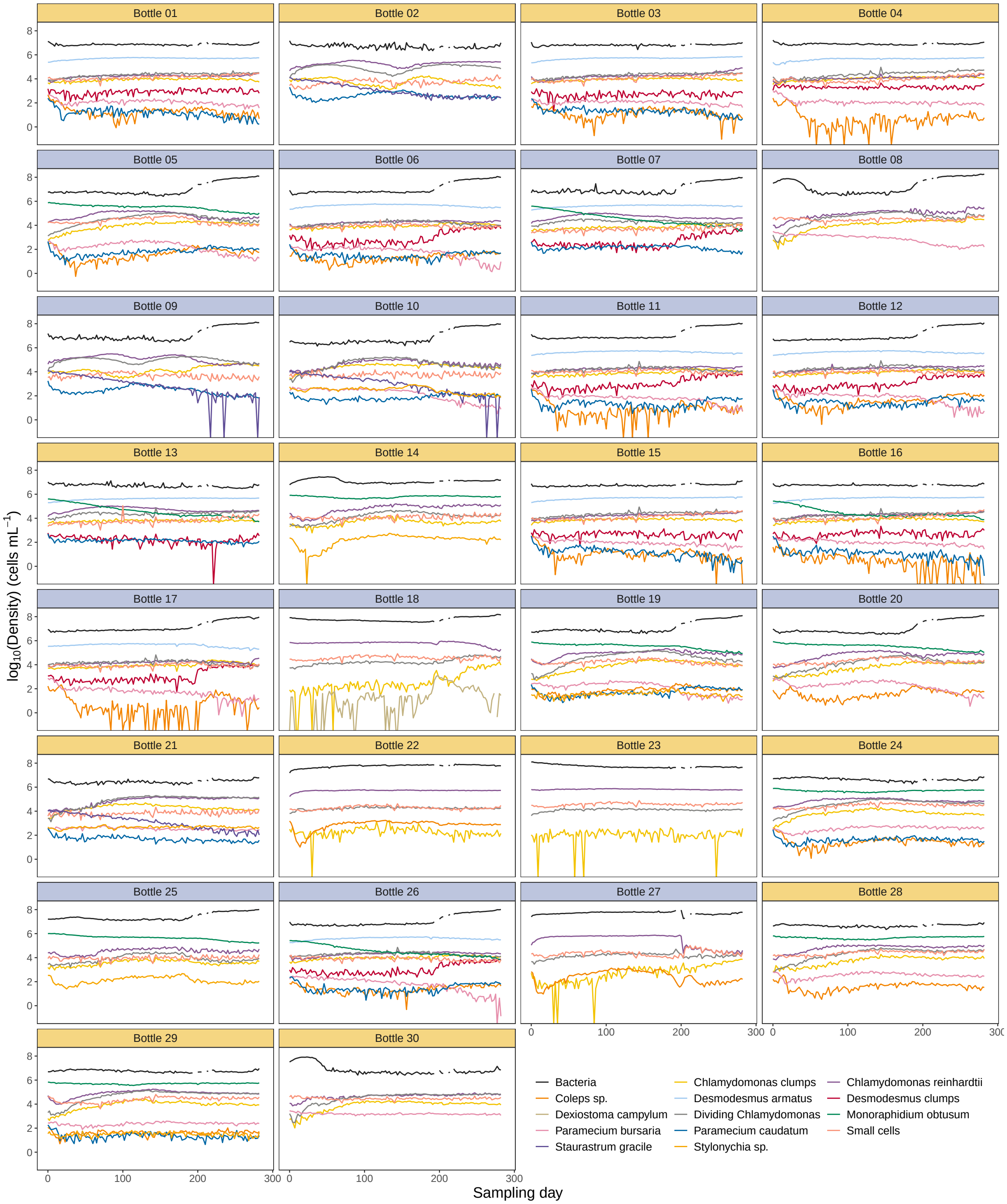

Figure S7: The ***untransformed*** biotic density time series that were used as forecast targets, shown for the different bottles (i.e. the panels). The time series are coloured by forecast target. The background colour of the subpanel strips indicates whether the bottles were in a constant (orange) or in a decreasing (blue) light environment.

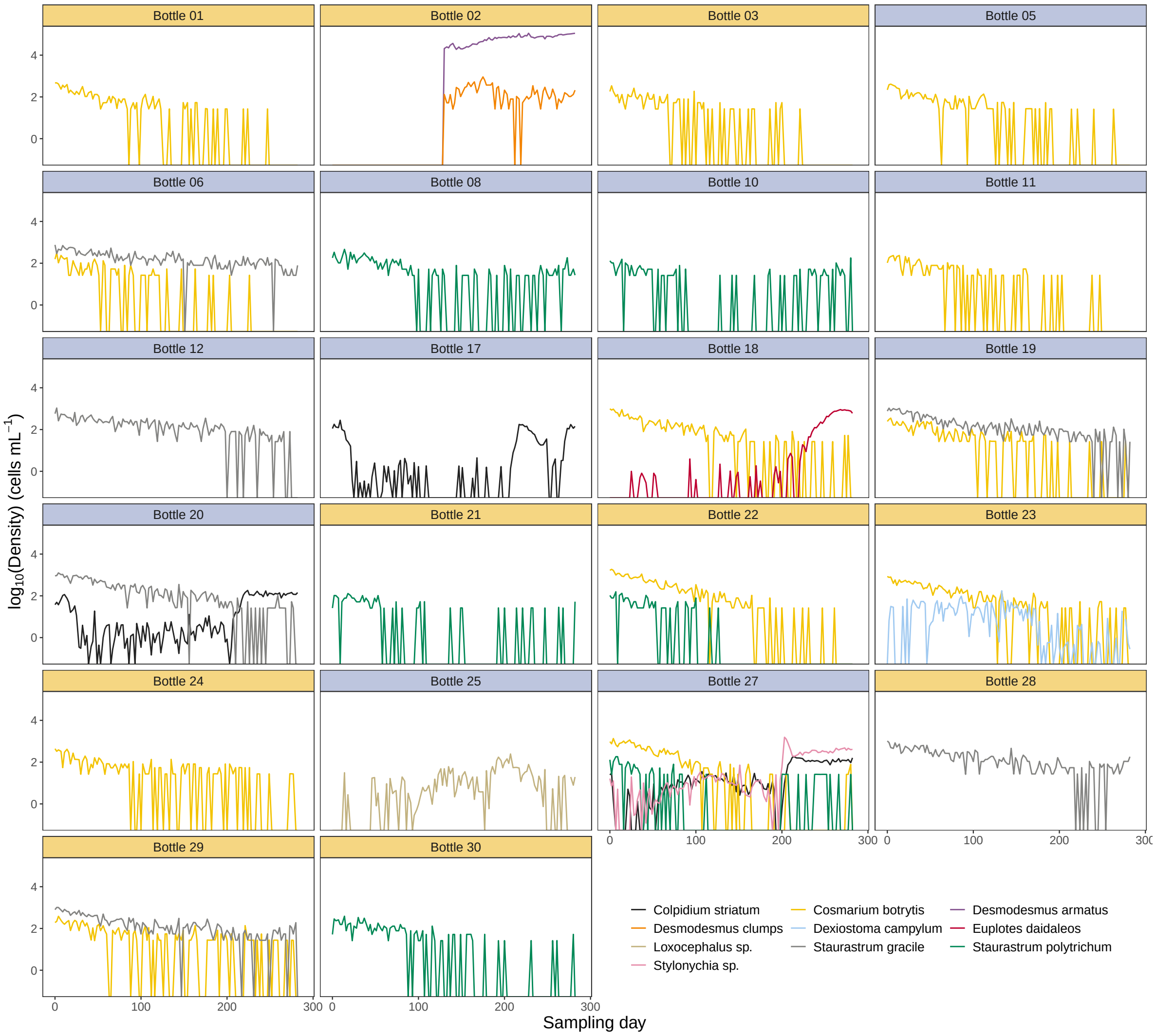

Figure S8: The ***untransformed*** biotic density time series that were ***not*** used as forecast targets, shown for the different bottles (i.e. the panels). The time series are coloured by taxon. The background colour of the subpanel strips indicates whether the bottles were in a constant (orange) or in a decreasing (blue) light environment.

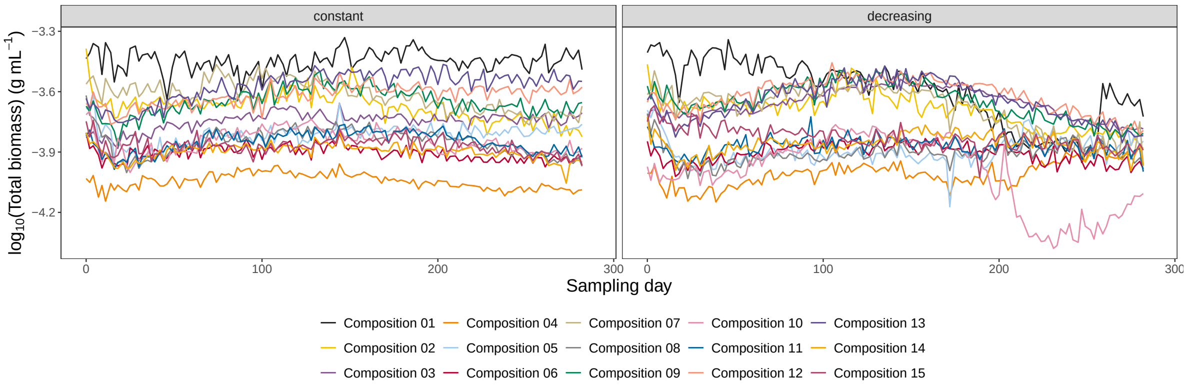

Figure S9: The ***untransformed*** total biomass time series of the communities, shown for the different light conditions (i.e. the panels). The time series are coloured by community composition.

### S4 Calculation of realized richness, biomass and time series properties

#### S4.1 The realized species richness

As described in a previous section, not all species persisted during the experiment. As a consequence, the planned species richness (see for instance Table S4) did not necessarily adequately represent the realized species richness in the bottles. As alternative measures of biodiversity, we also computed the minimal species richness (i.e. the number of species that were present throughout the entire study in the respective bottle) and the mean and the median species richness. The latter two were calculated by first counting the number of species present on each sampling day and then by taking respectively the mean and the median of them.

Figure S15 illustrates the distribution of these measures of richness as well as the correlation among them. The realized richness was generally smaller than the planned richness, but they were correlated and a diversity gradient remained despite the extinctions. Further, we checked if the experimental light conditions (i.e. constant versus gradual decline) affected the realized richness. Figure S16 shows the relation between the light conditions and realized species richness. Visually it does not appear that there is a significant difference in richness between the two light conditions, regardless of the species richness variable. We carried out a paired $t$-test for the mean species richness and confirmed that it was not affected by the light conditions ($t$ = -0.626, df = 14, $p$-value = 0.541, mean of the difference = -0.272).

#### S4.2 Time series metrics: autocorrelation, coefficient of variation and permutation entropy

We calculated several time series properties, as described in this section. The coefficient of variation (CV) is a measure of temporal variability. It is calculated as the ratio of the standard deviation and the mean, and

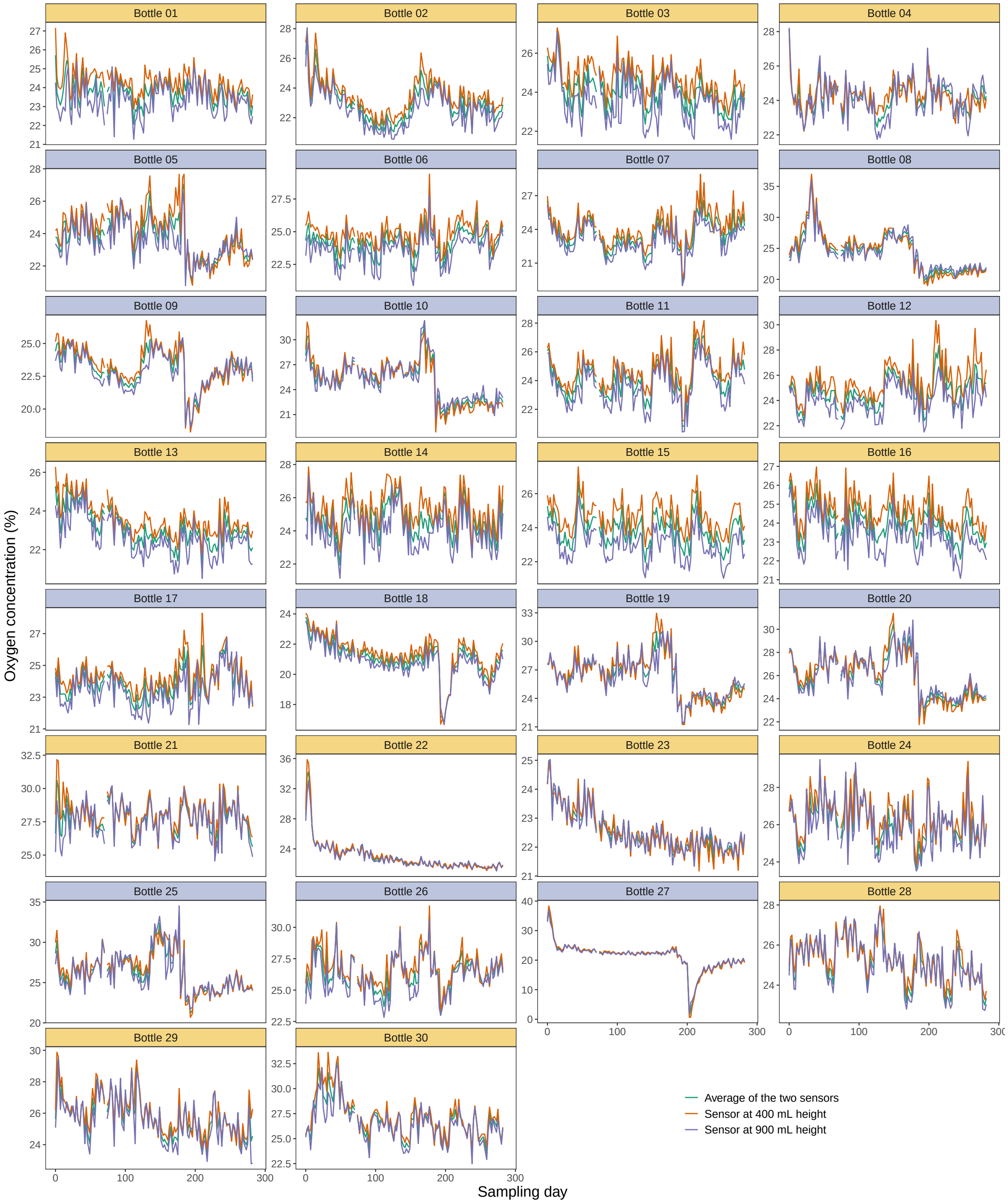

Figure S10: The ***untransformed*** oxygen concentration time series, shown for the different bottles (i.e. the panels). The time series are coloured by sensor height (with the average of the two time series also shown). The background colour of the subpanel strips indicates whether the bottles were in a constant (orange) or in a decreasing (blue) light environment.

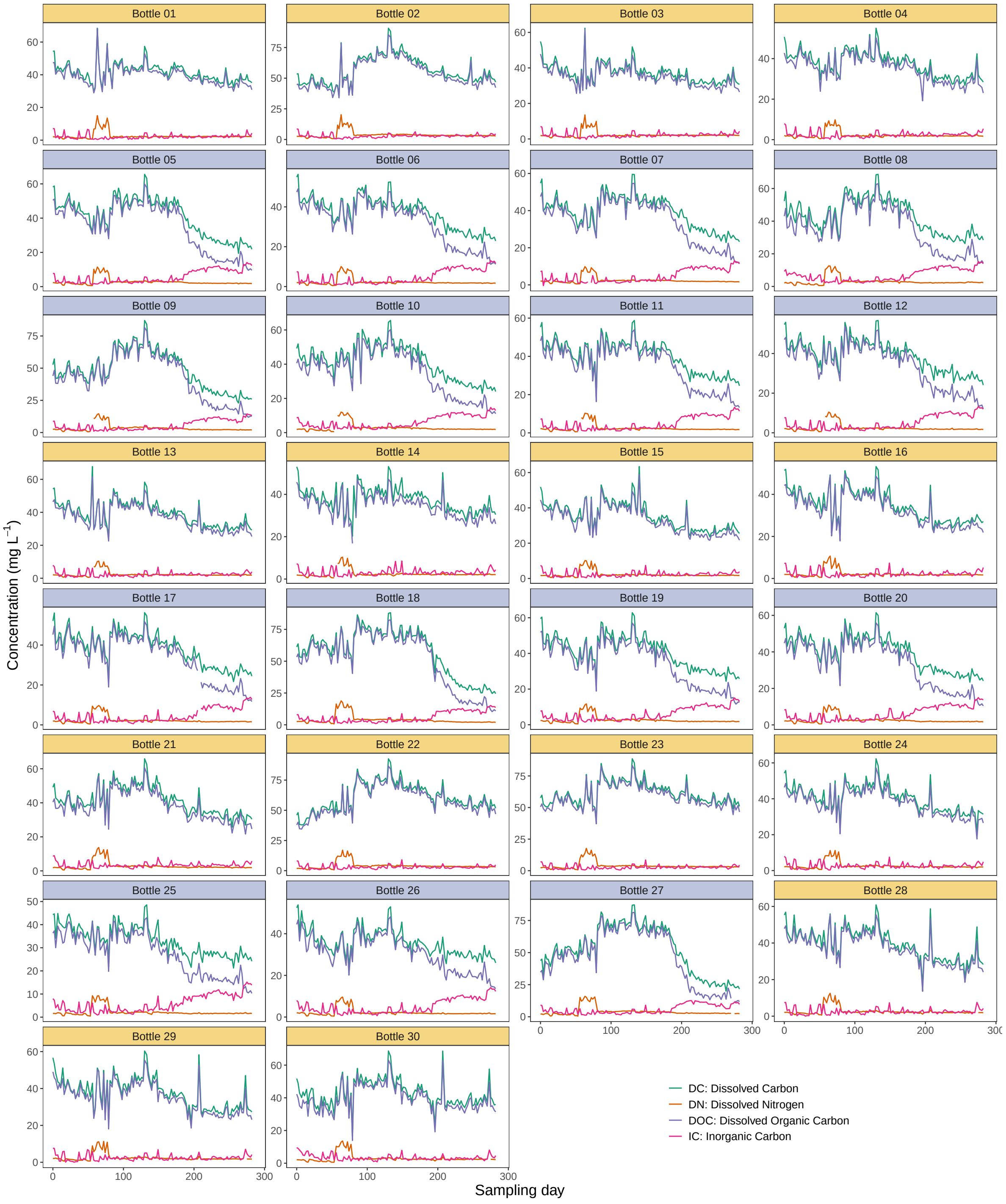

Figure S11: The ***untransformed*** carbon and nitrogen concentration time series, shown for the different bottles (i.e. the panels). The time series are coloured by the type of carbon and nitrogen concentration. The background colour of the subpanel strips indicates whether the bottles were in a constant (orange) or in a decreasing (blue) light environment.

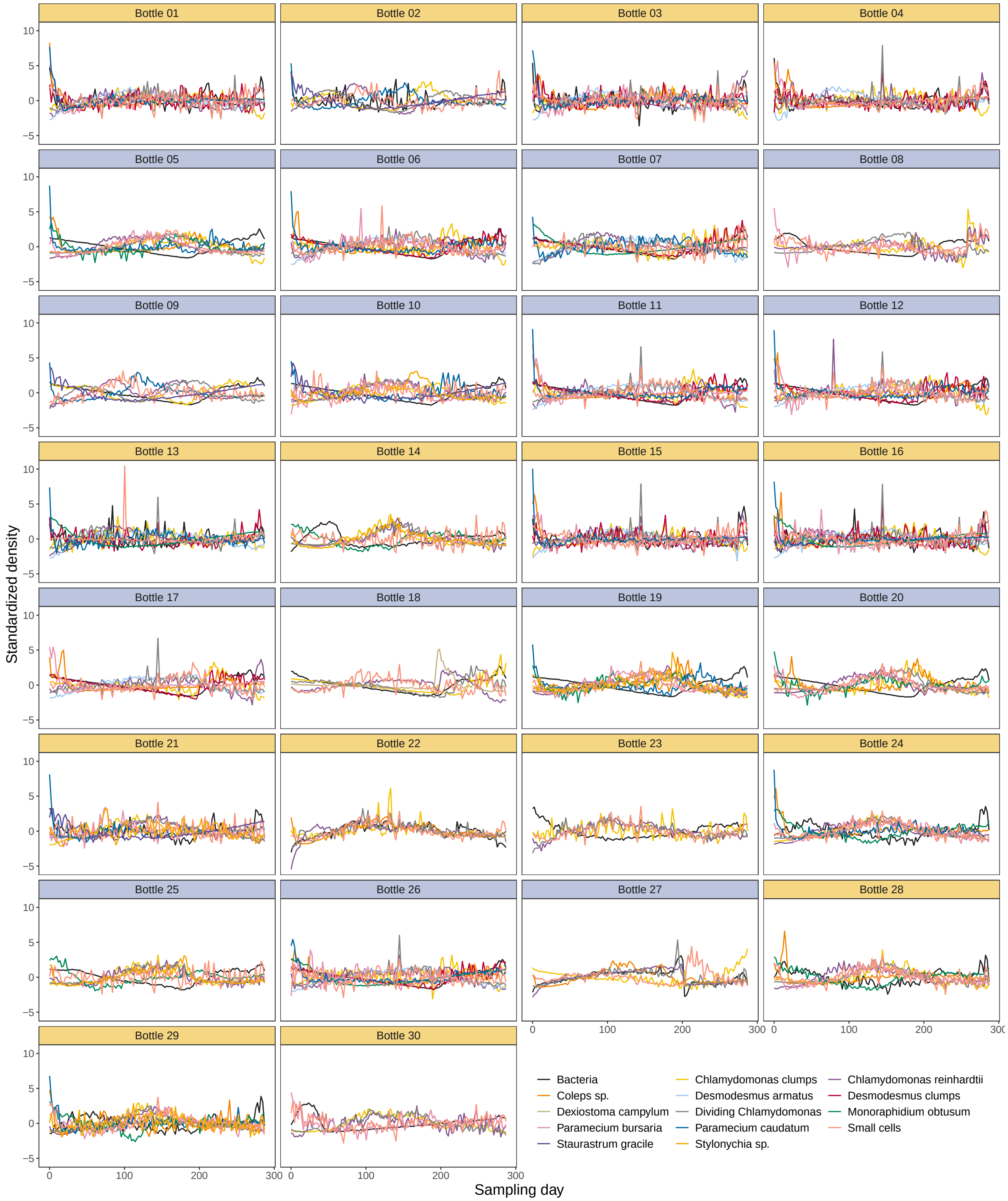

Figure S12: The ***transformed*** biotic density time series that were used as forecast targets, shown for the different bottles (i.e. the panels). The time series are coloured by forecast target. The background colour of the subpanel strips indicates whether the bottles were in a constant (orange) or in a decreasing (blue) light environment.

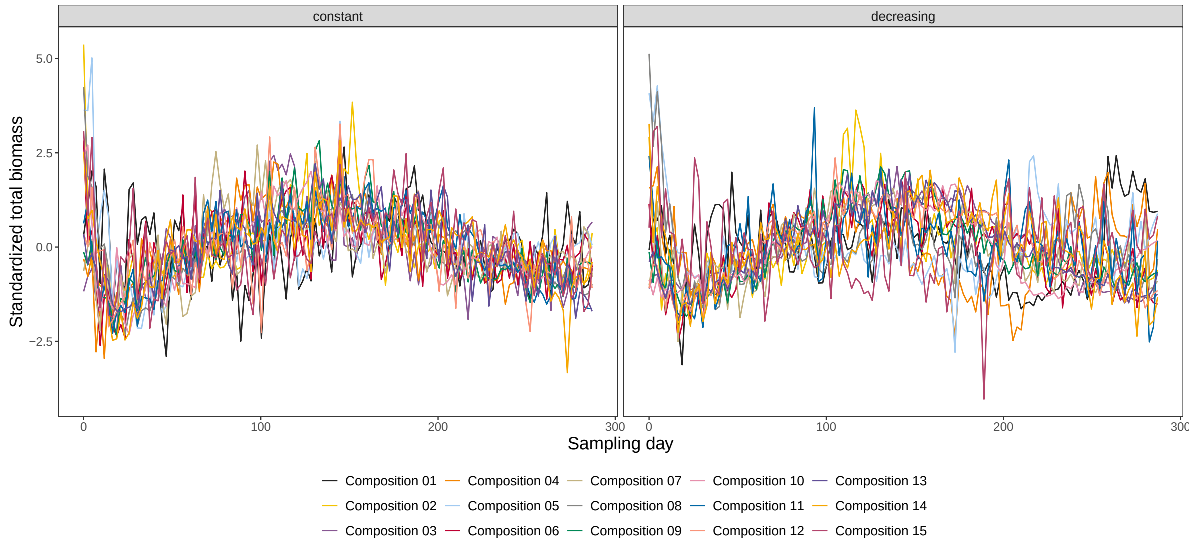

Figure S13: The ***transformed*** total biomass time series of the communities, shown for the different light conditions (i.e. the panels). The time series are coloured by community composition.

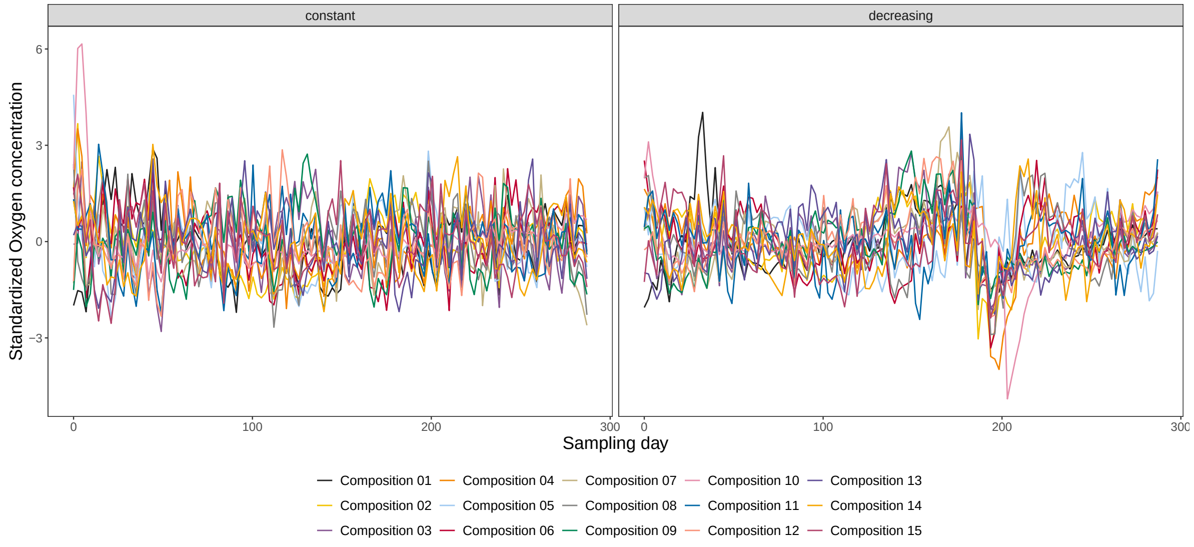

Figure S14: The ***transformed*** oxygen concentration time series, shown for the different light conditions (i.e. the panels). The time series are coloured by community composition.

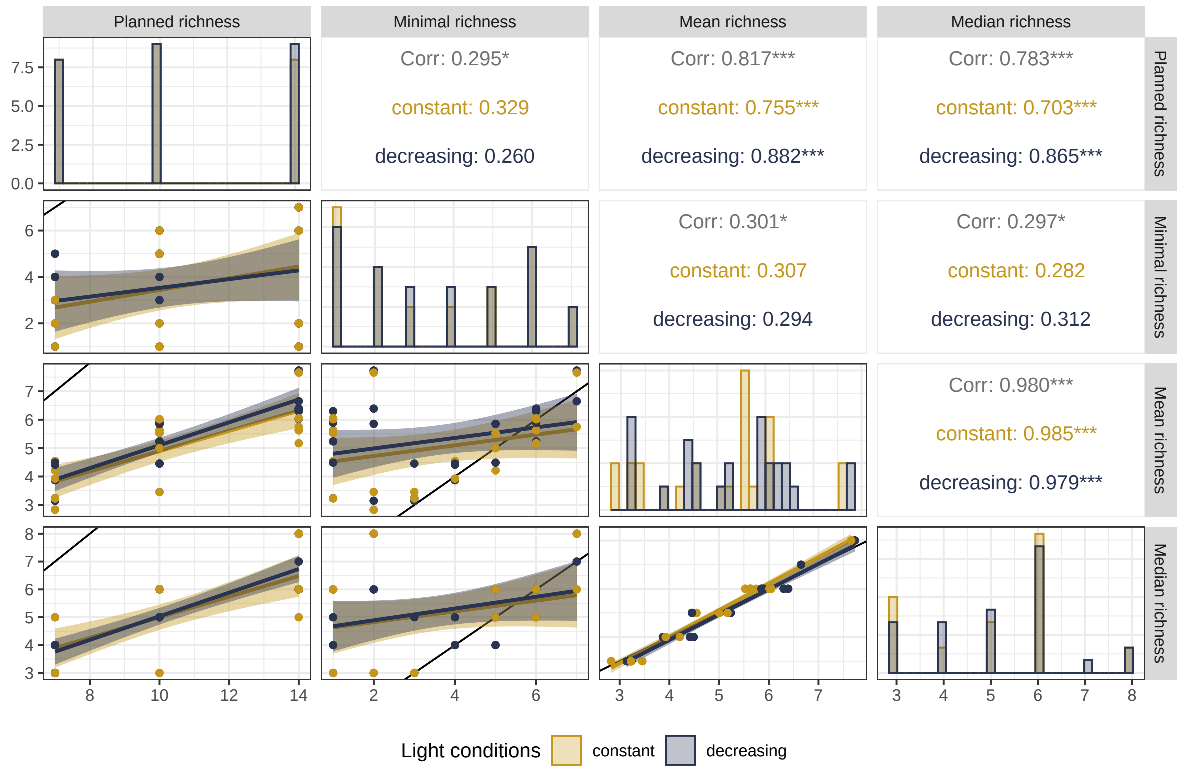

Figure S15: Comparison of the different species richness metrics (i.e. the planned and the realized median, mean and minimal richness), colour-coded for the light conditions. The panels in the diagonal show the barplots of the richness variables. The panels below the diagonal show the pairwise scatter-plots as indicated by the facet labels (the black line is the identity line, the coloured lines and the corresponding shaded ares show the fit and the 95% CI of a linear regression). The panels above the diagonal show the corresponding correlation estimates.

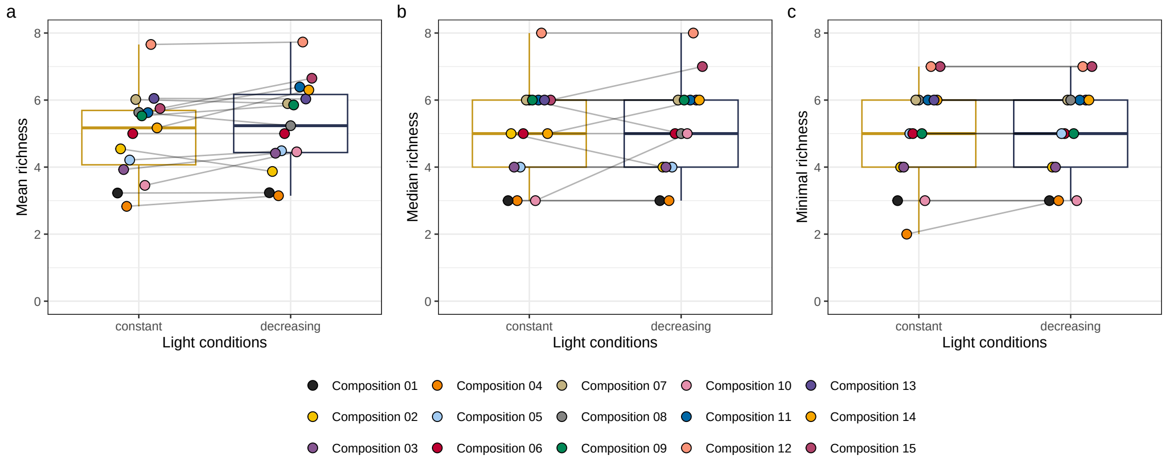

Figure S16: Scatter- and box-plot to investigate the effects of the light conditions on the realized species richness measures. (a) Mean richness; (b) Median richness; (c) Minimal richness. The points are coloured based on species composition and bottles with the same starting species composition are connected by a line. The boxplots are coloured based on light condition (orange = constant, blue = decreasing).

therefore greater values indicate greater variability (i.e. instability) in relation to the average value. Despite the simple formula for its calculation, care is needed if the time series is not stationary. For instance, if the time series is not trendless, the mean value might not be representative and thus the CV not meaningful. The usual way to deal with this is to not use the standard deviation of the time series itself but instead to use the one of the residuals of the regression of the time series values against time, while the mean remains the one of the original time series (see e.g. Donohue et al. 2013). Hence, we used this approach here as well.

The autocorrelation function (ACF) of a time series indicates how much the current value of a time series correlates with its past values. The ACF is usually calculated for each possible time delay (i.e. for all lags). In the main text we focus on the results at lag 5 (i.e. the correlation between current values and the value of 5 time points ago). However, the chosen lag value did not influence the findings. The calculation of the ACF was based on the transformed time series.

Lastly, we calculated the permutation entropy (PE) of the time series. The PE is used to measure the complexity of a time series and thus also a measure of its stochasticity (Pennekamp et al. 2019). Larger values of PE indicate a greater complexity and smaller values a better predictability. PE is therefore expected to correlate positively with forecast error. For its calculation, a time series is cut into snippets of a given length (i.e. the number of time points, called the word length). The snippets are then categorized into words based on time-ordered ranks of the values that they contain (i.e it is an ordinal analysis). Next, the frequencies of the different words are computed and the permutation entropy is then calculated based on their distribution (for more details see Pennekamp et al. 2019). For this study, we tried different word lengths and found that word length did not influence the results. For the results reported in this study we used a word length of five. The calculation of PE was based on the transformed time series.

#### S4.3 Calculation of species and community biomass

We calculated the biomass of the algae and ciliate species as follows. For the algae and ciliate species we used the extracted two-dimensional morphological traits (*width* and *length*) to extrapolate the volume of the tracked cells. We selected the three-dimensional shapes of the species based on Hillebrand et al. (1999) and on personal observations. We approximated the shape as an ellipsoid ($V = \frac{4}{3}\pi\frac{1}{2}(length \cdot height \cdot width)$, with $height = width$ if not specified otherwise) for the following species: *Chlamydomonas*, *Desmodesmus* ($height = width/4$), *Monoraphidium*, "Small cells", *Coleps* sp., *Colpidium*, *Paramecium caudatum*, *Dexiostoma*, *Euplotes* ($height = width/3$), *Loxocephalus*, *Paramecium bursaria* ($height = width/1.5$), *Stylonychia* sp. ($height = width/3$) and *Tetrahymena*. For *Monoraphidium*, because of its shape (i.e. long and potentially bent) we used the trait "geodesic thickness" as its width and height and the trait "geodesic length" as its length. For *Staurastrum gracile* we used two truncated cones ($V = (1/3) \cdot \pi \cdot h \cdot (r^2 + r \cdot R + R^2)$, with $R$ being the radius of the base of the cone, $r$ being the radius of the circle where the cone is truncated and $h$ being the distance between the base of the cone and where the cone is truncated). For this we used

$h = width/2$, $R = length/2$ and $r = length/6$. Further, for *Staurastrum polytrichum*, *Cosmarium* and "Dividing *Chlamydomonas*" we used two Ellipsoids as the shape ($V = 2\frac{4}{3}\pi\frac{1}{2}(a \cdot b \cdot c)$, with $a = width/2$, and $b = c = length/4$). Finally, for "*Chlamydomonas* clumps" and "*Desmodesmus* clumps" we used the extracted FlowCAM trait "Area_ABD" and multiplied it with the height of the clumps. We approximated the height of the clumps with the median width of the *Chlamydomonas* and *Desmodesmus* individuals in the monocultures (that we used for the classifier training). In a last step, we transformed the estimated volumes (in $\mu m^3$) into mass (in grams) by multiplying them with density of water ($10^{-12} g\ \mu m^{-3}$), i.e. we assumed the cells to have the density of water.

To calculate bacterial biomass, we first determined the relationship between forward scatter and size by measuring calibration beads of known size (Spherotech Nano Fluorescent Size Standard Kit: 0.45, 0.88, 1.25 µm; Sysmex Beads Mix: 1 µm). We used the resulting calibration curve ($size(\mu m) = -0.626007 + 0.336922 * log10(FSC)$) to estimate the length of the measured bacteria from their forward scatter. We calculated the biovolume of each particle assuming an ellipsoidal shape ($width = length/3$) and converted to biomass assuming their density to be that of water.

We estimated the community biomass, i.e. total biomass per bottle, by summing the biomass of the species present.

### S5 Forecasting

#### S5.1 The forecast methods

In the following subsection greater detail is given regarding the used forecasting methods and their implementation and parametrization. Of note is that because we were primarily interested in achieving the best possible forecasts we did allow the various models to search a wide parameter space. In other words, we did not want the models to be limited by the chosen parametrization. The trade-off of this is that the interpretability of the model parameters is reduced (this mostly applies to the ARIMA forecasts).

##### S5.1.1 Empirical Dynamic Modeling EDM

Empirical Dynamic Modeling (EDM, Ye and Sugihara 2016) is a nonlinear time series forecasting method in which future values are predicted based on how the forecasted system progressed when it was in a similar state at other times (which are, therefore, not necessarily close in time). The states of the system are determined by a state-space reconstruction of the (shadow) attractor manifold for which the relevant state variables are used (e.g. species abundance time series). Based on Takens' theorem that interacting state variables leave imprints of each other in their time series (Takens 1981), the state variables can also be lagged for this reconstruction. The number of (lagged and non-lagged) time series used for the attractor manifold reconstruction is called the embedding dimension $E$. The EDM functions are available in the

R-package rEDM (Park et al. 2021).

Simplex EDM (Sugihara and May 1990) is a version of EDM in which the attractor manifold is reconstructed by using only the forecasted time series itself and the appropriate lagged versions of it (i.e. it is a univariate approach). The forecasting is then done based on averaging the progression of the nearest neighbors in the reconstructed state-space. Because this approach only considers a single time series and thus has a limited dimensionality, the optimal value of the embedding dimension $E$ (which, in this case, is determined by the number of different lags used) can easily be determined (function `EmbedDimension()`) and then used in the function `Simplex()`. In this study, we allowed the embedding dimension $E$ to be as high as 40. The draw-back of this method is that it does only consider a small part of the system and is thus limited.

Multiview EDM (Ye and Sugihara 2016) is one of the techniques that extend Simplex EDM to multivariate data. More importantly, it is specifically built to deal with systems with a high-dimensionality (i.e. many state variables, e.g. many species and variable abiotic system components present). It does so by reducing the dimensionality of the system by fitting many low-dimension models, which are then ranked by in-sample forecast skill. A certain number ($k$) are then selected from the best models to carry out out-of-sample forecasts which are then averaged (i.e. an ensemble forecast). To fit these low-dimensional models it is crucial that the embedding dimension $E$ and the maximal lag $l$ are restrained to relatively small numbers, as otherwise the number of models to be fitted increases dramatically and the computational power and time that are needed quickly become too big. Here, we set the embedding dimension to $E = 3$ and the maximum lag to $l = 4$. Separately for every forecast target and bottle, we varied the number of retained low-dimensional models by trying out 30 values of $k$ logarithmically spaced between 1 and 111 (the number of time points used for the training of the models) and then kept the models that resulted in the lowest forecast errors. We used the function `Multiview()`.

##### S5.1.2 Recurrent Neural Network RNN

A Recurrent Neural Network (RNN) is a type of artificial neural network. This machine learning approach makes minimal assumptions on the data and works by creating and weighing connections between artificial neurons across different layers based on current and previous variable (i.e. predictor) values. Starting from an input layer, in RNN a network is created in which the input variables (the predictor variables) determine the node values in this layer, and these nodes then determine the values of the nodes in the next layer. The complexity of the model increases the more layers and neurons it is allowed to have. In the final layer, all previous nodes and connections converge to desired output dimension (e.g. the forecasted abundance), again with the associated connection weights determining by how much the various combinations of the variables (i.e. the various "neural pathways") influence the final output. In an iterative procedure (i.e. by repeatedly fitting the network by building on the previous version), the error in the training data (i.e. the error between observed and fitted values) is reduced by adjusting the weights associated to the neuron connections. For more information see Hewamalage, Bergmeir, and Bandara (2021). We used the function `keras_model_sequential()` (and related functions) implemented in the R-package `keras` (Allaire and Chollet 2022). We set our network up to have three layers (with the respective dimensions 7, 49 and 1) and used 30 epochs (i.e. we iteratively fitted the whole data 30 times). Please see our accompanying R-code to this publication for further details regarding our RNN implementation.

###### S5.1.3 Random Forest

Random Forest (RF) is a linear model and machine learning approach based on randomly generating regression trees, also known as decision trees. Among its most common uses is classification, but as in our case, it can also be used to other ends such as forecasting (Cutler, Cutler, and Stevens 2012). In this method, a certain number of regression trees are generated by randomly selecting the features (i.e. the variables, in this case the predictors) at each node, and in this way a random forest is "grown" (and thus the name "Random Forest"). The forecasts are then based on a summary of all the trees, and usually the average is used (i.e. RF is also an ensemble forecast method). In our implementation, in addition to the predictors as described in the main text, we also used them in their lagged fashion (as in the EDM case) with the maximal again being $l = 4$. Further, we used the function `randomForest()` from the R-package `randomForest` (Liaw and Wiener 2002), and used its default parameter values.

###### S5.1.4 ARIMA

ARIMA (Auto-Regressive Integrated Moving Average) is a well-established time series forecasting method in which the forecasts are linearly based on the values closest in time. For this study, we used the function `auto.arima()` from the R-package `forecast` (Rob J. Hyndman and Khandakar 2008; R. Hyndman et al. 2022). As we were interested in achieving the best possible forecasts regardless of interpretability, we allowed the function to search a wide parameter space (the maximum values for the parameters $p$, $q$ and $d$ were respectively 10, 10 and 3, with the order not exceeding 15, see the above citations for more information).

##### S5.2 Correlation of forecasts across methods

As described, we used multiple approaches to forecast the targets. In the main text we focus on the results of the Simplex EDM forecasts, but in this section we show that the different forecasts methods agreed with each other (see also the robustness analyses in a later section). The distribution of forecast errors from the different methods is shown in the diagonal panels in Figure S17. In the same figure, in the off-diagonal panels, the correlation among the forecast errors from the different methods is shown (including scatter-plots and regression results, see caption). As can be seen, the forecast errors were all strongly, significantly and positively correlated, suggesting that the results did not depend on the chosen forecast approach.

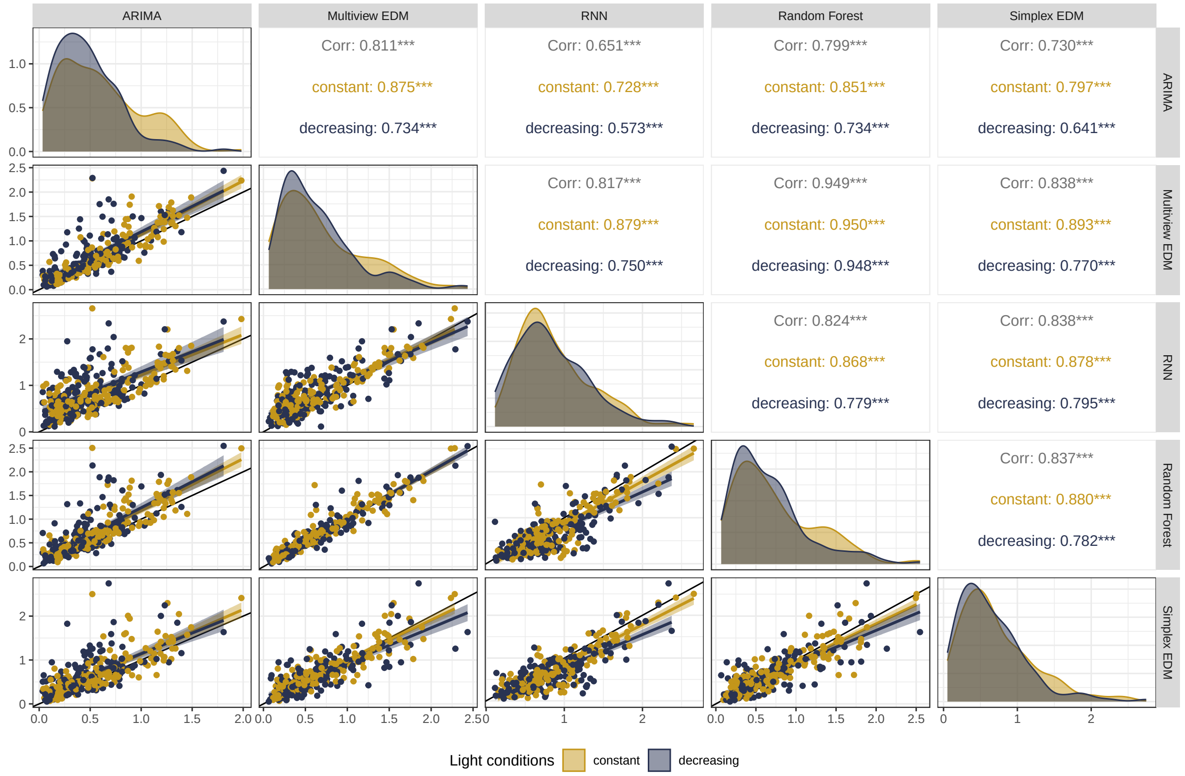

Figure S17: Comparison of forecast errors values (i.e. RMSE) produced by the different forecast methods (i.e. ARIMA, Multiview EDM, RNN, Random Forest and Simplex EDM), colour-coded for the light conditions. The panels in the diagonal show the density plots of the forecast errors. The panels below the diagonal show the pairwise scatter-plots as indicated by the facet labels (the black line is the identity line, the coloured lines and the corresponding shaded ares show the fit and the 95% CI of a linear regression). The panels above the diagonal showing the corresponsing correlation estimates.

### S6 Additional figures and tables

In the main text, we describe the effects of the light conditions and of the realized species richness on the taxon abundance forecasts and the community biomass forecasts. We also describe the effects of the light conditions and of the realized species richness on the time series metrics (autocorrelation, permutation entropy and coefficient of variation) of the taxa abundance time series, as well as the relations between the time series metrics and the taxa abundance forecast errors. We used linear mixed models in which we included all pairwise interactions between the fixed effects and assessed the significance of these interactions with Type III ANOVAs. If the interactions were significant or marginally significant (i.e. $p$-values below 0.1) we kept them in the respective models and used Type III ANOVAs. Otherwise, we removed them from the models and used Type II ANOVAs to investigate the importance of the main fixed effects. Note that: 1) The choice of ANOVA type (Type II or III) does not matter for determining the importance of the interaction; 2) If no interaction is included in the model, then Type II and Type III ANOVAs will produce the same results for the main effects (i.e. the Type III is reduced to a Type II ANOVA); 3) A significant interaction already indicates that the corresponding variables are important in the model, regardless of whether the main effects are estimated to be significant or not (which, in the presence of a significant interaction, after all also depends on the observed values of these variables). The figures and tables of these and other models are reported in the following subsections.

#### S6.1 Effects of experimental conditions on forecasts

Table S6 is an extended version of Table 1 (main text) and provides more information regarding the fitted models. As the interaction between the light conditions and the median richness was significant in Table S6 for the forecast error of the taxa abundances, we additionally also investigated the effects of the richness variable on this response variable separately for the two light conditions (Table S7, for Simplex EDM only). We again found some evidence that the forecast errors increased with increasing richness when the light conditions were constant and that they decreased with increasing richness when the light decreased (the profile-likelihood based 95% confidence intervals did not overlap with the null effect, i.e. with zero, and the corresponding $p$-values were weakly significant, Table S7). Note that for this analysis we log10-transformed the median richness to meet the model assumptions. This was not necessary in the full model (i.e. the non-partitioned model). Further, Table S8 and Figure S18 report the results of the mixed models investigating the relation between the experimental treatments and the taxa abundance forecasts for not only Simplex EDM but for all forecast methods used.

There are several ways how the explained variance (i.e. the coefficient of determination $R^2$) of a mixed model can be calculated, with, to our knowledge, no clear agreed upon standard approach. In fact, Rights and Sterba (2019) explain 12 different approaches to partition the variance. Further, please also note that for mixed effects models the partial $R^2$ values not necessarily sum up to the model $R^2$ and therefore need to be interpreted with care. As a representative example of the other forecast models, in this paragraph we report the (partial) $R^2$ values of the mixed model investigating the relation between the experimental treatments and the taxa abundance forecasts based on Simplex EDM forecasting. For this, we used two options implemented in the function `r2beta` in the R-package `r2glmm` (Jaeger 2017), namely the Kenward-Roger and the Nakagawa-Schielzeth approaches (for details please see the documentation of this package). The achieved total and partial $R^2$ values with the Kenward-Roger approach are: 0.255 (fixed effects of full model), 0.168 (light conditions), 0.002 (median richness), 0.204 (interaction). Likewise, with the Nakagawa-Schielzeth approach: 0.023 (fixed effects of full model), 0.009 (light conditions), 0.010 (median richness), 0.018 (interaction). Alternatively, the effect sizes in terms of variance explained can also be calculated with the function `eta_squared` implemented in the R-package `effectsize` (Ben-Shachar, Lüdecke, and Makowski 2020). In this case, the achieved values are the same as with the Kenward-Roger approach above: 0.17 (light conditions), 0.002 (median richness), 0.20 (interaction). As can be seen, the estimated proportions of variances explained vary a lot based on the method used, and in the case of the Nakagawa-Schielzeth approach they are very low. However, even small values can be useful in explaining relations between quantities. Further, the amount of variation explained might be small not necessarily because effects sizes are small (although clearly that is a possibility) but also because there might be a considerable amount of variation or noise present in the data (typical for biological data). Care is therefore needed in defining a threshold of variation explained that needs to be achieved so that a model is considered useful. Similarly, because of the random effects it not straightforward to visually see the variance explained in a plot of the data and the model fit (such as Figure 1 in the main text).

Further, Tables S10 and S11 and Figures S19 and S20 respectively report the regression results for the models that investigated how the forecast errors of community biomass and mean oxygen concentration depended on the light conditions and the species richness, separately for the different forecasting methods (See Supplementary Section S7 for the robustness analyses regarding the forecast errors of the different taxa). As mentioned in the main text, while generally the trends matched the ones found for the taxa forecasts, (i.e. forecasts appeared to be better in the decreased light conditions at high richness than in the in constant light conditions), in most cases neither of the experimental treatments had a significant effect on the forecast errors of community biomass and oxygen concentration, respectively.

To test whether community level measures (i.e. community biomass, oxygen concentration) are forecasted better than the taxa abundances, we also fitted two mixed models in which we compared the forecast error of taxa abundances with the forecast error of community biomass and oxygen concentration, respectively. The Tables S12 and S13 report the results and the specifics of these models. To this end, we carried out a post-hoc analysis to investigate whether there was a significant difference in forecast error of respectively the community biomass and the mean oxygen concentration when compared to the mean forecast error of taxa abundances. We used the function `emmeans()` and `pairs()` from R-package `emmeans` (Lenth 2023)

Table S6: Results of linear mixed models investigating the effects of species richness and light on forecast errors of taxa and community biomass. This table is an extended version of Table 1 in the main text and includes both the regression tables and the analysis of variance tables of the models (top part of the table: Type III ANOVA; bottom part of the table Type II ANOVA). The models included median centered richness, light treatment and their interaction as fixed effects. In the first model, random intercepts were included for the bottles, incubators and the targets (i.e. taxa). In the second model, random intercepts were included for composition and incubators. Forecast error is based on forecasts computed with Simplex EDM. The estimates for the random intercepts are standard deviations.

| Response | Covariate | Type | Estimate | DF | SE | Test | Value | *p*-value |
|---|---|---|---|---|---|---|---|---|
| Forecast error of target abundances | Intercept | Fixed effect | 0.685 | 16 | 0.093 | *t* | 7.371 | <0.0001 |
| | Richness | | 0.056 | 22 | 0.033 | *t* | 1.695 | 0.1044 |
| | Light (decreasing) | | -0.095 | 6 | 0.086 | *t* | -1.109 | 0.3092 |
| | Richness:Light | | -0.102 | 20 | 0.046 | *t* | -2.246 | 0.0363 |
| | Target (taxa) | Random intercept | 0.074 | 1 | - | $\chi^2$ | 47.396 | <0.0001 |
| | Bottle | | 0.250 | 1 | - | $\chi^2$ | 0.410 | 0.5219 |
| | Incubator | | 0.085 | 1 | - | $\chi^2$ | 1.198 | 0.2737 |
| | Richness | Sum of squares | 0.008 | 1, 21.1 | - | *F* | 0.046 | 0.8328 |
| | Light (decreasing) | | 0.225 | 1, 6.1 | - | *F* | 1.230 | 0.3092 |
| | Richness:Light | | 0.922 | 1, 19.7 | - | *F* | 5.046 | 0.0363 |
| Forecast error of community biomass | Intercept | Fixed effect | 0.657 | 20 | 0.108 | *t* | 6.065 | <0.0001 |
| | Richness | | 0.015 | 20 | 0.062 | *t* | 0.241 | 0.8118 |
| | Light (decreasing) | | -0.017 | 14 | 0.099 | *t* | -0.167 | 0.8698 |
| | Composition | Random intercept | 0.302 | 1 | - | $\chi^2$ | 5.108 | 0.0238 |
| | Incubator | | 0.001 | 1 | - | $\chi^2$ | 0.000 | 1.0000 |
| | Richness | Sum of squares | 0.004 | 1, 19.8 | - | *F* | 0.058 | 0.8118 |
| | Light (decreasing) | | 0.002 | 1, 14.1 | - | *F* | 0.028 | 0.8698 |

which calculates the specified marginal estimated means and their differences and tests whether the latter significantly differ from zero. As written in the main text, in almost all cases there was no significant difference (see Table S14).

Table S7: Relation between log10-transformed median richness and the forecast errors of taxa abundances, with the model carried out separately for the constant and the decreasing light conditions (i.e. split analysis). The table includes the estimated coefficients and corresponding 95% confidence intervals, as well as the analysis of variance (Type II) table. The random intercepts were included for the bottles, incubators and the targets (i.e. taxa), but are not shown in the table. Note that the richness variable was log10-transformed to meet the model assumptions. This was not necessary in the full model (i.e. the non-partitioned model). Forecast error is based on forecasts computed with Simplex EDM.

| Light | Covariate | Estimate | 95% CI | SS | DF | *F*-value | *p*-value |
|---|---|---|---|---|---|---|---|
| Constant | Intercept | 0.158 | -0.371 - 0.697 | | | | |
| | log10(Richness) | 0.716 | 0.033 - 1.39 | 0.768 | 1, 12.6 | 4.118 | 0.0642 |
| Decreasing | Intercept | 1.169 | 0.609 - 1.753 | | | | |
| | log10(Richness) | -0.782 | -1.576 - -0.023 | 0.617 | 1, 12.5 | 4.064 | 0.0659 |

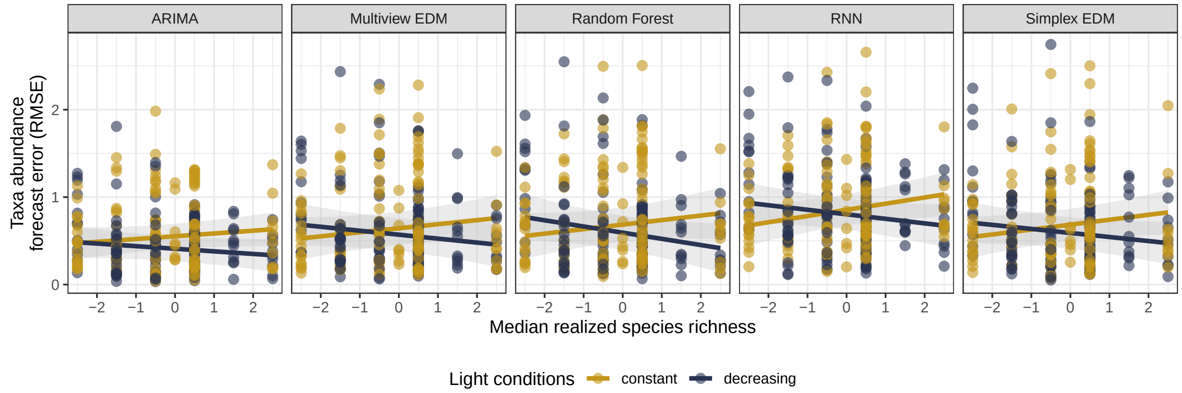

Figure S18: Effects of species richness and light on the forecast error of taxa abundances. Forecasts are for 30 bottles (yellow: constant light, blue: decreasing light) with varying number of taya per bottle (see Figure 1 in the main text). The panels show the results for five different forecasting methods, i.e. ARIMA (Auto-Regressive Integrated Moving Average), Multiview EDM (Empirical Dynamic Modeling), Random Forest, RNN (Recurrent Neural Networks), and Simplex EDM. Lines display the fit of the corresponding linear mixed model, shaded areas denote the 95% confidence intervals. Composition and incubator were included as random intercepts in the linear mixed model.

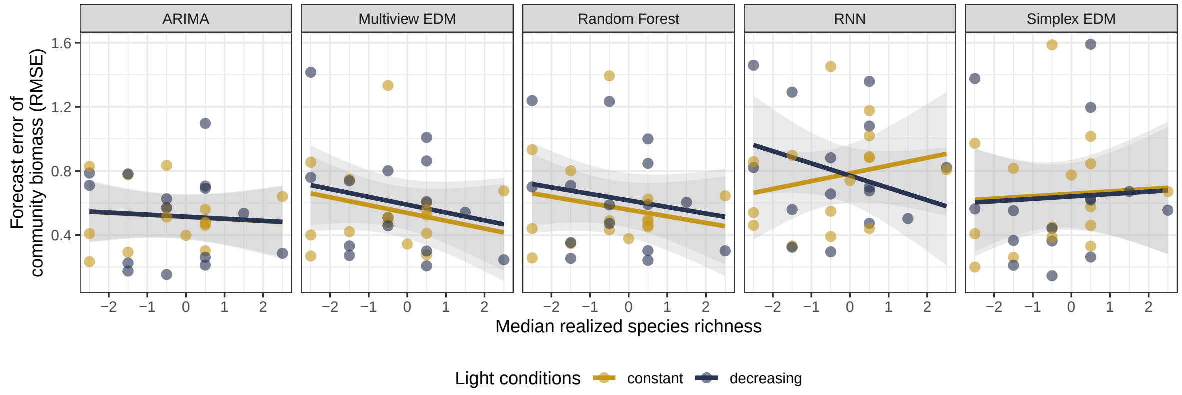

Figure S19: Effects of species richness and light on the forecast error of community biomass. Forecasts are for 30 bottles (yellow: constant light, blue: decreasing light). The panels show the results for five different forecasting methods, i.e. ARIMA (Auto-Regressive Integrated Moving Average), Multiview EDM (Empirical Dynamic Modeling), Random Forest, RNN (Recurrent Neural Networks), and Simplex EDM. Lines display the fit of the corresponding linear mixed model, shaded areas denote the 95% confidence intervals. Composition and incubator were included as random intercepts in the linear mixed model.

Table S8: Results of the regressions investigating the effects of species richness and light on the forecast errors of the taxa abundances (for different forecast methods). The estimates for the random intercepts are standard deviations.

| Response | Method | Covariate | Type | Estimate | DF | SE | Test | Value | $p$-value |
|---|---|---|---|---|---|---|---|---|---|
| Forecast error of taxa abundances | Simplex EDM | Intercept | Fixed effect | 0.685 | 16 | 0.093 | $t$ | 7.371 | <0.0001 |
| | | Richness | | 0.056 | 22 | 0.033 | $t$ | 1.695 | 0.1044 |
| | | Light (decreasing) | | -0.095 | 6 | 0.086 | $t$ | -1.109 | 0.3092 |
| | | Richness:Light | | -0.102 | 20 | 0.046 | $t$ | -2.246 | 0.0363 |
| | | Target (taxa) | Random intercept | 0.074 | 1 | - | $\chi^2$ | 47.396 | <0.0001 |
| | | Bottle | | 0.250 | 1 | - | $\chi^2$ | 0.410 | 0.5219 |
| | | Incubator | | 0.085 | 1 | - | $\chi^2$ | 1.198 | 0.2737 |
| | Multiview EDM | Intercept | Fixed effect | 0.644 | 16 | 0.097 | $t$ | 6.649 | <0.0001 |
| | | Richness | | 0.046 | 22 | 0.033 | $t$ | 1.387 | 0.1792 |
| | | Light (decreasing) | | -0.074 | 6 | 0.090 | $t$ | -0.821 | 0.4436 |
| | | Richness:Light | | -0.091 | 20 | 0.046 | $t$ | -1.991 | 0.0602 |
| | | Target (taxa) | Random intercept | 0.090 | 1 | - | $\chi^2$ | 59.107 | <0.0001 |
| | | Bottle | | 0.260 | 1 | - | $\chi^2$ | 1.086 | 0.2973 |
| | | Incubator | | 0.091 | 1 | - | $\chi^2$ | 1.385 | 0.2393 |
| | ARIMA | Intercept | Fixed effect | 0.552 | 16 | 0.073 | $t$ | 7.531 | <0.0001 |
| | | Richness | | 0.032 | 24 | 0.024 | $t$ | 1.335 | 0.1945 |
| | | Light (decreasing) | | -0.143 | 6 | 0.053 | $t$ | -2.675 | 0.0365 |
| | | Richness:Light | | -0.061 | 22 | 0.033 | $t$ | -1.872 | 0.0747 |
| | | Target (taxa) | Random intercept | 0.055 | 1 | - | $\chi^2$ | 60.559 | <0.0001 |
| | | Bottle | | 0.225 | 1 | - | $\chi^2$ | 0.534 | 0.4648 |
| | | Incubator | | 0.042 | 1 | - | $\chi^2$ | 0.388 | 0.5333 |
| | RNN | Intercept | Fixed effect | 0.854 | 17 | 0.092 | $t$ | 9.309 | <0.0001 |
| | | Richness | | 0.071 | 23 | 0.033 | $t$ | 2.166 | 0.0409 |
| | | Light (decreasing) | | -0.051 | 6 | 0.071 | $t$ | -0.717 | 0.5004 |
| | | Richness:Light | | -0.123 | 21 | 0.045 | $t$ | -2.710 | 0.0131 |
| | | Target (taxa) | Random intercept | 0.095 | 1 | - | $\chi^2$ | 73.686 | <0.0001 |
| | | Bottle | | 0.275 | 1 | - | $\chi^2$ | 1.531 | 0.2159 |
| | | Incubator | | 0.049 | 1 | - | $\chi^2$ | 0.207 | 0.6495 |
| | Random Forest | Intercept | Fixed effect | 0.685 | 15 | 0.102 | $t$ | 6.751 | <0.0001 |
| | | Richness | | 0.052 | 22 | 0.039 | $t$ | 1.311 | 0.2034 |
| | | Light (decreasing) | | -0.092 | 6 | 0.099 | $t$ | -0.931 | 0.3880 |
| | | Richness:Light | | -0.122 | 20 | 0.055 | $t$ | -2.242 | 0.0362 |
| | | Target (taxa) | Random intercept | 0.136 | 1 | - | $\chi^2$ | 51.747 | <0.0001 |
| | | Bottle | | 0.261 | 1 | - | $\chi^2$ | 3.753 | 0.0527 |
| | | Incubator | | 0.092 | 1 | - | $\chi^2$ | 0.831 | 0.3619 |

Table S9: Contrast table comparing the achieved forecast errors of taxa abundances between the two light levels (decreasing and constant) at the highest and lowest achieved (centered) median species richness. The estimate is the difference between the achieved RMSE in the decreasing light setting and the one achieved in the constant light setting.

| Contrast | Centered median richness | Method | Estimate | SE | DF | $t$-value | $p$-value |
|---|---|---|---|---|---|---|---|
| Const. light vs. Decr. light | -2.5 | Simplex EDM | 0.161 | 0.137 | 24.225 | 1.180 | 0.2496 |
| | | Multiview EDM | 0.155 | 0.139 | 22.903 | 1.117 | 0.2757 |
| | | ARIMA | 0.011 | 0.093 | 27.560 | 0.114 | 0.9098 |
| | | Random Forest | 0.214 | 0.159 | 23.909 | 1.349 | 0.1899 |
| | | RNN | 0.256 | 0.126 | 27.791 | 2.030 | 0.0520 |
| | 2.5 | Simplex EDM | -0.351 | 0.151 | 21.970 | -2.333 | 0.0292 |
| | | Multiview EDM | -0.302 | 0.154 | 22.010 | -1.961 | 0.0626 |
| | | ARIMA | -0.297 | 0.104 | 22.572 | -2.848 | 0.0092 |
| | | Random Forest | -0.398 | 0.181 | 23.186 | -2.205 | 0.0376 |
| | | RNN | -0.357 | 0.143 | 23.096 | -2.497 | 0.0201 |

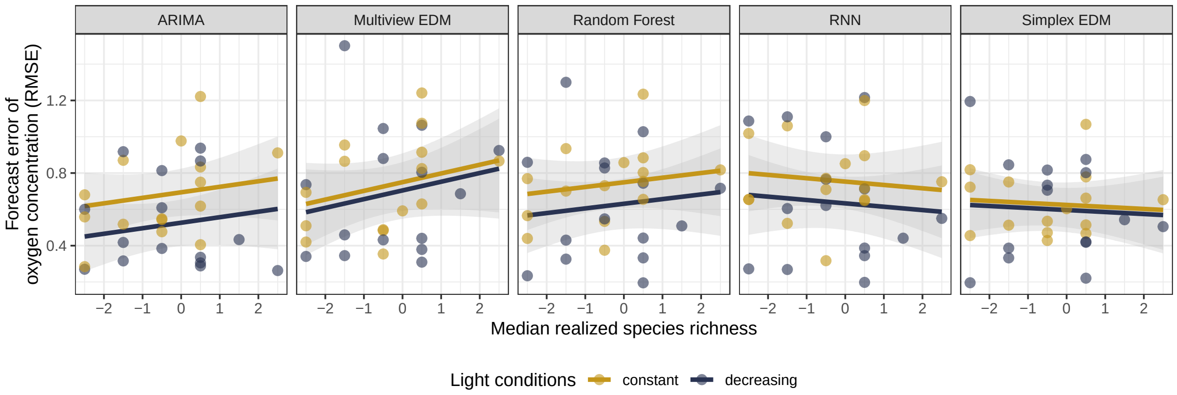

Figure S20: Effects of species richness and light on the forecast error of oxygen concentration. Forecasts are for 30 bottles (yellow: constant light, blue: decreasing light). The panels show the results for five different forecasting methods, i.e. ARIMA (Auto-Regressive Integrated Moving Average), Multiview EDM (Empirical Dynamic Modeling), Random Forest, RNN (Recurrent Neural Networks), and Simplex EDM. Lines display the fit of the corresponding linear mixed model, shaded areas denote the 95% confidence intervals. Composition and incubator were included as random intercepts in the linear mixed model.

Table S10: Results of the regressions investigating the effects of species richness and light on the forecast errors of the community biomass (for different forecast methods). The estimates for the random intercepts are standard deviations.

| Response | Method | Covariate | Type | Estimate | DF | SE | Test | Value | *p*-value |
|---|---|---|---|---|---|---|---|---|---|
| Forecast error of community biomass | Simplex EDM | Intercept | Fixed effect | 0.657 | 20 | 0.108 | - | - | - |
| | | Richness | | 0.015 | 20 | 0.062 | *F* | 0.058 | 0.8118 |
| | | Light (decreasing) | | -0.017 | 14 | 0.099 | *F* | 0.028 | 0.8698 |
| | | Composition | Random intercept | 0.302 | 1 | - | $\chi^2$ | 5.108 | 0.0238 |
| | | Incubator | | 0.001 | 1 | - | $\chi^2$ | 0.000 | 1.0000 |
| | Multiview EDM | Intercept | Fixed effect | 0.538 | 8 | 0.082 | - | - | - |
| | | Richness | | -0.049 | 15 | 0.045 | *F* | 1.191 | 0.2921 |
| | | Light (decreasing) | | 0.051 | 3 | 0.094 | *F* | 0.298 | 0.6197 |
| | | Composition | Random intercept | 0.170 | 1 | - | $\chi^2$ | 1.418 | 0.2337 |
| | | Incubator | | 0.027 | 1 | - | $\chi^2$ | 0.003 | 0.9574 |
| | ARIMA | Intercept | Fixed effect | 0.513 | 6 | 0.071 | - | - | - |
| | | Richness | | -0.013 | 22 | 0.032 | *F* | 0.165 | 0.6883 |
| | | Light (decreasing) | | 0.001 | 5 | 0.099 | *F* | 0.000 | 0.9890 |
| | | Composition | Random intercept | 0.000 | 1 | - | $\chi^2$ | 0.000 | 1.0000 |
| | | Incubator | | 0.066 | 1 | - | $\chi^2$ | 0.144 | 0.7047 |
| | RNN | Intercept | Fixed effect | 0.785 | 19 | 0.093 | - | - | - |
| | | Richness | | 0.049 | 25 | 0.059 | *F* | 0.071 | 0.7924 |
| | | Light (decreasing) | | -0.015 | 13 | 0.079 | *F* | 0.035 | 0.8554 |
| | | Richness:Light | | -0.125 | 13 | 0.056 | *F* | 4.963 | 0.0442 |
| | | Composition | Random intercept | 0.274 | 1 | - | $\chi^2$ | 6.706 | 0.0096 |
| | | Incubator | | 0.000 | 1 | - | $\chi^2$ | 0.000 | 1.0000 |
| | Random Forest | Intercept | Fixed effect | 0.558 | 8 | 0.088 | - | - | - |
| | | Richness | | -0.041 | 15 | 0.045 | *F* | 0.821 | 0.3792 |
| | | Light (decreasing) | | 0.058 | 4 | 0.106 | *F* | 0.305 | 0.6106 |
| | | Composition | Random intercept | 0.166 | 1 | - | $\chi^2$ | 1.182 | 0.2769 |
| | | Incubator | | 0.068 | 1 | - | $\chi^2$ | 0.094 | 0.7594 |

Table S11: Results of the regressions investigating the effects of species richness and light on the forecast errors of mean oxygen (for different forecast methods). The estimates for the random intercepts are standard deviations.

| Response | Method | Covariate | Type | Estimate | DF | SE | Test | Value | $p$-value |
|---|---|---|---|---|---|---|---|---|---|
| Forecast error of mean oxygen | Simplex EDM | Intercept | Fixed effect | 0.624 | 27 | 0.063 | - | - | - |
| | | Richness | | -0.011 | 27 | 0.032 | $F$ | 0.121 | 0.7304 |
| | | Light (decreasing) | | -0.028 | 27 | 0.087 | $F$ | 0.103 | 0.7502 |
| | | Composition | Random intercept | 0.000 | 1 | - | $\chi^2$ | 0.000 | 1.0000 |
| | | Incubator | | 0.000 | 1 | - | $\chi^2$ | 0.000 | 1.0000 |
| | Multiview EDM | Intercept | Fixed effect | 0.750 | 27 | 0.081 | - | - | - |
| | | Richness | | 0.048 | 15 | 0.042 | $F$ | 1.334 | 0.2659 |
| | | Light (decreasing) | | -0.046 | 14 | 0.108 | $F$ | 0.178 | 0.6799 |
| | | Composition | Random intercept | 0.075 | 1 | - | $\chi^2$ | 0.050 | 0.8237 |
| | | Incubator | | 0.000 | 1 | - | $\chi^2$ | 0.000 | 1.0000 |
| | ARIMA | Intercept | Fixed effect | 0.694 | 27 | 0.067 | - | - | - |
| | | Richness | | 0.030 | 27 | 0.033 | $F$ | 0.834 | 0.3692 |
| | | Light (decreasing) | | -0.168 | 27 | 0.092 | $F$ | 3.342 | 0.0786 |
| | | Composition | Random intercept | 0.000 | 1 | - | $\chi^2$ | 0.000 | 1.0000 |
| | | Incubator | | 0.000 | 1 | - | $\chi^2$ | 0.000 | 1.0000 |
| | RNN | Intercept | Fixed effect | 0.753 | 27 | 0.076 | - | - | - |
| | | Richness | | -0.018 | 27 | 0.038 | $F$ | 0.234 | 0.6322 |
| | | Light (decreasing) | | -0.120 | 27 | 0.105 | $F$ | 1.300 | 0.2643 |
| | | Composition | Random intercept | 0.000 | 1 | - | $\chi^2$ | 0.000 | 1.0000 |
| | | Incubator | | 0.000 | 1 | - | $\chi^2$ | 0.000 | 1.0000 |
| | Random Forest | Intercept | Fixed effect | 0.749 | 27 | 0.072 | - | - | - |
| | | Richness | | 0.026 | 27 | 0.036 | $F$ | 0.506 | 0.4829 |
| | | Light (decreasing) | | -0.118 | 27 | 0.099 | $F$ | 1.414 | 0.2448 |
| | | Composition | Random intercept | 0.000 | 1 | - | $\chi^2$ | 0.000 | 1.0000 |
| | | Incubator | | 0.000 | 1 | - | $\chi^2$ | 0.000 | 1.0000 |

Table S12: Results of the regressions investigating whether the community level measure total biomass is forecasted better than taxa based forecasts (here: mean forecast error across all forecasted taxa). As explanatory variables, the model included median centered richness, light treatment and measure (i.e. total biomass forecasts versus taxa abundance forecasts), as well as all pairwise interactions. The table reports the results for all considered forecast methods. The estimates for the random intercepts are standard deviations.

| Response | Method | Covariate | Type | Estimate | DF | SE | Test | Value | $p$-value |
|---|---|---|---|---|---|---|---|---|---|
| Mean forecast error of abundances Versus Forecast error of community biomass | Simplex EDM | Intercept | Fixed effect | 0.715 | 42 | 0.075 | - | - | - |
| | | Richness | | -0.001 | 27 | 0.033 | $F$ | 0.000 | 0.9868 |
| | | Light (decreasing) | | -0.032 | 27 | 0.090 | $F$ | 0.130 | 0.7214 |
| | | Measure (biomass) | | -0.056 | 29 | 0.072 | $F$ | 0.607 | 0.4422 |
| | | Bottle | Random intercept | 0.148 | 1 | - | $\chi^2$ | 1.322 | 0.2503 |
| | | Incubator | | 0.000 | 1 | - | $\chi^2$ | 0.000 | 1.0000 |
| | Multiview EDM | Intercept | Fixed effect | 0.669 | 34 | 0.062 | - | - | - |
| | | Richness | | 0.013 | 26 | 0.040 | $F$ | 1.724 | 0.2006 |
| | | Light (decreasing) | | -0.029 | 26 | 0.080 | $F$ | 0.129 | 0.7228 |
| | | Measure (biomass) | | -0.082 | 29 | 0.045 | $F$ | 3.293 | 0.0799 |
| | | Richness:Light | | -0.100 | 26 | 0.056 | $F$ | 3.210 | 0.0848 |
| | | Bottle | Random intercept | 0.171 | 1 | - | $\chi^2$ | 7.074 | 0.0078 |
| | | Incubator | | 0.000 | 1 | - | $\chi^2$ | 0.000 | 1.0000 |
| | ARIMA | Intercept | Fixed effect | 0.549 | 9 | 0.054 | - | - | - |
| | | Richness | | -0.015 | 22 | 0.021 | $F$ | 0.487 | 0.4925 |
| | | Light (decreasing) | | -0.057 | 6 | 0.069 | $F$ | 0.684 | 0.4404 |
| | | Measure (biomass) | | -0.007 | 29 | 0.041 | $F$ | 0.031 | 0.8623 |
| | | Bottle | Random intercept | 0.115 | 1 | - | $\chi^2$ | 3.311 | 0.0688 |
| | | Incubator | | 0.051 | 1 | - | $\chi^2$ | 0.261 | 0.6095 |
| | RNN | Intercept | Fixed effect | 0.894 | 39 | 0.065 | - | - | - |
| | | Richness | | 0.053 | 26 | 0.040 | $F$ | 0.111 | 0.7421 |
| | | Light (decreasing) | | -0.030 | 26 | 0.080 | $F$ | 0.137 | 0.7142 |
| | | Measure (biomass) | | -0.099 | 29 | 0.059 | $F$ | 2.831 | 0.1032 |
| | | Richness:Light | | -0.124 | 26 | 0.056 | $F$ | 4.888 | 0.0360 |
| | | Bottle | Random intercept | 0.137 | 1 | - | $\chi^2$ | 1.829 | 0.1763 |
| | | Incubator | | 0.000 | 1 | - | $\chi^2$ | 0.000 | 1.0000 |
| | Random Forest | Intercept | Fixed effect | 0.692 | 36 | 0.066 | - | - | - |
| | | Richness | | -0.038 | 27 | 0.030 | $F$ | 1.562 | 0.2221 |
| | | Light (decreasing) | | 0.011 | 27 | 0.084 | $F$ | 0.016 | 0.8997 |
| | | Measure (biomass) | | -0.109 | 29 | 0.051 | $F$ | 4.561 | 0.0413 |
| | | Bottle | Random intercept | 0.181 | 1 | - | $\chi^2$ | 6.084 | 0.0136 |
| | | Incubator | | 0.000 | 1 | - | $\chi^2$ | 0.000 | 1.0000 |

Table S13: Results of the regressions investigating whether the community level measure mean oxygen concentration is forecasted better than taxa based forecasts (here: mean forecast error across all forecasted taxa). As explanatory variables, the model included median centered richness, light treatment and measure (i.e. oxygen concentration forecasts versus taxa abundance forecasts), as well as all pairwise interactions. The table reports the results for all considered forecast methods. The estimates for the random intercepts are standard deviations.

| Response | Method | Covariate | Type | Estimate | DF | SE | Test | Value | $p$-value |
|---|---|---|---|---|---|---|---|---|---|
| Mean forecast error of abundances Versus Forecast error of mean oxygen concentration | Simplex EDM | Intercept | Fixed effect | 0.711 | 39 | 0.052 | - | - | - |
| | | Richness | | -0.019 | 27 | 0.024 | $F$ | 0.639 | 0.4309 |
| | | Light (decreasing) | | -0.037 | 27 | 0.065 | $F$ | 0.331 | 0.5701 |
| | | Measure (biomass) | | -0.085 | 29 | 0.046 | $F$ | 3.415 | 0.0748 |
| | | Bottle | Random intercept | 0.125 | 1 | - | $\chi^2$ | 3.295 | 0.0695 |
| | | Incubator | | 0.000 | 1 | - | $\chi^2$ | 0.000 | 1.0000 |
| | Multiview EDM | Intercept | Fixed effect | 0.673 | 42 | 0.061 | - | - | - |
| | | Richness | | -0.027 | 51 | 0.034 | $F$ | 0.160 | 0.6926 |
| | | Light (decreasing) | | -0.039 | 27 | 0.073 | $F$ | 0.279 | 0.6014 |
| | | Measure (biomass) | | 0.074 | 28 | 0.058 | $F$ | 1.582 | 0.2189 |
| | | Richness:Measure | | 0.076 | 28 | 0.041 | $F$ | 3.454 | 0.0736 |
| | | Bottle | Random intercept | 0.127 | 1 | - | $\chi^2$ | 1.824 | 0.1769 |
| | | Incubator | | 0.000 | 1 | - | $\chi^2$ | 0.000 | 1.0000 |
| | ARIMA | Intercept | Fixed effect | 0.620 | 42 | 0.047 | - | - | - |
| | | Richness | | 0.050 | 26 | 0.028 | $F$ | 0.147 | 0.7050 |
| | | Light (decreasing) | | -0.173 | 26 | 0.056 | $F$ | 9.514 | 0.0048 |
| | | Measure (biomass) | | 0.072 | 29 | 0.047 | $F$ | 2.341 | 0.1369 |
| | | Richness:Light | | -0.085 | 26 | 0.039 | $F$ | 4.639 | 0.0407 |
| | | Bottle | Random intercept | 0.074 | 1 | - | $\chi^2$ | 0.559 | 0.4545 |
| | | Incubator | | 0.000 | 1 | - | $\chi^2$ | 0.000 | 1.0000 |
| | RNN | Intercept | Fixed effect | 0.918 | 42 | 0.057 | - | - | - |
| | | Richness | | 0.023 | 26 | 0.034 | $F$ | 0.940 | 0.3411 |
| | | Light (decreasing) | | -0.092 | 26 | 0.068 | $F$ | 1.812 | 0.1898 |
| | | Measure (biomass) | | -0.177 | 29 | 0.056 | $F$ | 9.921 | 0.0038 |
| | | Richness:Light | | -0.092 | 26 | 0.048 | $F$ | 3.701 | 0.0654 |
| | | Bottle | Random intercept | 0.093 | 1 | - | $\chi^2$ | 0.669 | 0.4134 |
| | | Incubator | | 0.000 | 1 | - | $\chi^2$ | 0.000 | 1.0000 |
| | Random Forest | Intercept | Fixed effect | 0.771 | 41 | 0.058 | - | - | - |
| | | Richness | | 0.042 | 26 | 0.035 | $F$ | 0.080 | 0.7792 |
| | | Light (decreasing) | | -0.116 | 26 | 0.071 | $F$ | 2.695 | 0.1127 |
| | | Measure (biomass) | | -0.032 | 29 | 0.056 | $F$ | 0.323 | 0.5740 |
| | | Richness:Light | | -0.099 | 26 | 0.049 | $F$ | 3.977 | 0.0567 |
| | | Bottle | Random intercept | 0.104 | 1 | - | $\chi^2$ | 0.961 | 0.3269 |
| | | Incubator | | 0.000 | 1 | - | $\chi^2$ | 0.000 | 1.0000 |

Table S14: Table reporting the differences (i.e. the contrasts) between the marginal estimated mean of the forecast errors of the community level variables (total biomas and mean oxygen concentration) and of the mean taxa abundance forecast error. Also reported are the results of the tests of whether the estimated differences are different from zero.

| Contrast | Method | Light | Estimate | SE | DF | $t$-Value | $p$-value |
|---|---|---|---|---|---|---|---|
| Species - Biomass | Simplex EDM | constant | 0.056 | 0.072 | 29 | 0.779 | 0.4422 |
| | | decreasing | 0.056 | 0.072 | 29 | 0.779 | 0.4422 |
| | Multiview EDM | constant | 0.082 | 0.045 | 29 | 1.815 | 0.0799 |
| | | decreasing | 0.082 | 0.045 | 29 | 1.815 | 0.0799 |
| | ARIMA | constant | 0.007 | 0.041 | 29 | 0.175 | 0.8623 |
| | | decreasing | 0.007 | 0.041 | 29 | 0.175 | 0.8623 |
| | RNN | constant | 0.099 | 0.059 | 29 | 1.683 | 0.1032 |
| | | decreasing | 0.099 | 0.059 | 29 | 1.683 | 0.1032 |
| | Random Forest | constant | 0.109 | 0.051 | 29 | 2.136 | 0.0413 |
| | | decreasing | 0.109 | 0.051 | 29 | 2.136 | 0.0413 |
| Species - Oxygen | Simplex EDM | constant | 0.085 | 0.046 | 29 | 1.848 | 0.0748 |
| | | decreasing | 0.085 | 0.046 | 29 | 1.848 | 0.0748 |
| | Multiview EDM | constant | -0.044 | 0.056 | 28 | -0.788 | 0.4374 |
| | | decreasing | -0.044 | 0.056 | 28 | -0.788 | 0.4374 |
| | ARIMA | constant | -0.072 | 0.047 | 29 | -1.530 | 0.1369 |
| | | decreasing | -0.072 | 0.047 | 29 | -1.530 | 0.1369 |
| | RNN | constant | 0.177 | 0.056 | 29 | 3.150 | 0.0038 |
| | | decreasing | 0.177 | 0.056 | 29 | 3.150 | 0.0038 |
| | Random Forest | constant | 0.032 | 0.056 | 29 | 0.569 | 0.5740 |
| | | decreasing | 0.032 | 0.056 | 29 | 0.569 | 0.5740 |

#### S6.2 Relations of time series properties with forecast errors and with experimental conditions

As described in the main text, we investigated the relation between time series properties and the experimental conditions and the forecast errors (see Figure 2 in the main text). The corresponding tables with the model fit information are provided in this section. Table S15 lists the regression and ANOVA tables regarding the models in which we regressed the forecasts errors (response variable) against the time series metrics (coefficient of variation, autocorrelation at lag 5 and permutation entropy). Table S16 lists the regression and ANOVA tables for the models in which the three time series metrics were the response variables and regressed against the experimental conditions (i.e. light and species richness).

#### S6.3 Structural equation model

To carry out the structural equation model (SEM) we used the R-package `lavaan.survey` (Oberski 2014). We respected our experimental design in the SEM analysis by including target, incubator and bottle as a combined random effect variable. We further included covariances between the errors of endogenous model variables if they improved the model goodness-of-fit. Table S17 reports the detailed output of the model (i.e. model coefficients, uncertainties and statistical tests), while Table S18 reports the achieved R-squared values.

Table S15: Results of linear mixed models investigating the relationship between forecast error (response variable) and time series metrics. The models were fitted separately for the explanatory variables "autocorrelation" (at lag 5), "log10-transformed coefficient of variation", and "permutation entropy". This table includes both the regression tables and the analysis of variance (Type II) tables of the models. The models included the respective time series metrics as a covariate, alongside the light conditions and their interaction as fixed effects. The random intercepts were included for the bottles, incubators and the targets (i.e. forecasted taxa). Forecast error is based on forecasts computed with Simplex EDM. The estimates for the random intercepts are standard deviations.

| Main covariate | Covariate | Type | Estimate | DF | SE | Test | Value | *p*-value |
|---|---|---|---|---|---|---|---|---|
| Autocorrelation (lag = 5) | Intercept | Fixed effect | 0.972 | 26 | 0.101 | *t* | 9.650 | <0.0001 |
| | ACF (lag 5) | | -0.690 | 249 | 0.111 | *t* | -6.239 | <0.0001 |
| | Light (decreasing) | | 0.009 | 7 | 0.069 | *t* | 0.126 | 0.9038 |
| | Target (taxa) | Random intercept | 0.000 | 1 | - | $\chi^2$ | 65.009 | <0.0001 |
| | Bottle | | 0.269 | 1 | - | $\chi^2$ | 0.000 | 1.0000 |
| | Incubator | | 0.061 | 1 | - | $\chi^2$ | 1.035 | 0.3089 |
| | ACF (lag 5) | Sum of squares | 6.399 | 1, 248.5 | - | *F* | 38.923 | <0.0001 |
| | Light (decreasing) | | 0.003 | 1, 6.5 | - | *F* | 0.016 | 0.9038 |
| log10-transformed Cofficient of variation) | Intercept | Fixed effect | 0.439 | 25 | 0.110 | *t* | 3.977 | 0.0005 |
| | log10(CV) | | -0.519 | 227 | 0.117 | *t* | -4.424 | <0.0001 |
| | Light (decreasing) | | -0.001 | 6 | 0.094 | *t* | -0.014 | 0.9893 |
| | Target (taxa) | Random intercept | 0.083 | 1 | - | $\chi^2$ | 50.055 | <0.0001 |
| | Bottle | | 0.257 | 1 | - | $\chi^2$ | 0.740 | 0.3896 |
| | Incubator | | 0.099 | 1 | - | $\chi^2$ | 2.075 | 0.1497 |
| | log10(CV) | Sum of squares | 3.313 | 1, 227.3 | - | *F* | 19.573 | <0.0001 |
| | Light (decreasing) | | 0.000 | 1, 6.3 | - | *F* | 0.000 | 0.9893 |
| Permutation entropy | Intercept | Fixed effect | -0.079 | 147 | 0.219 | *t* | -0.360 | 0.7193 |
| | PE | | 1.162 | 245 | 0.307 | *t* | 3.789 | 0.0002 |
| | Light (decreasing) | | -0.029 | 6 | 0.101 | *t* | -0.288 | 0.7829 |
| | Target (taxa) | Random intercept | 0.134 | 1 | - | $\chi^2$ | 47.132 | <0.0001 |
| | Bottle | | 0.234 | 1 | - | $\chi^2$ | 3.509 | 0.0610 |
| | Incubator | | 0.100 | 1 | - | $\chi^2$ | 1.281 | 0.2577 |
| | PE | Sum of squares | 2.423 | 1, 245.2 | - | *F* | 14.357 | 0.0002 |
| | Light (decreasing) | | 0.014 | 1, 6 | - | *F* | 0.083 | 0.7829 |

Table S16: Results of linear mixed models investigating the the effect of species richness and light on time series metrics (response variables). The models were fitted separately for the response variables "autocorrelation" (at lag 5), "log10-transformed coefficient of variation", and "permutation entropy". This table includes both the regression tables and the analysis of variance (Type II) tables of the models. The models included median centered richness, light treatment and their interaction as fixed effects. The random intercepts were included for the bottles, incubators and the targets (i.e. forecasted taxa). Forecast error is based on forecasts computed with Simplex EDM. The estimates for the random intercepts are standard deviations.

| Response | Covariate | Type | Estimate | DF | SE | Test | Value | $p$-value |
|---|---|---|---|---|---|---|---|---|
| Autocorrelation (lag = 5) | Intercept | Fixed effect | 0.431 | 16 | 0.054 | $t$ | 8.017 | <0.0001 |
| | Richness | | 0.004 | 22 | 0.016 | $t$ | 0.281 | 0.7811 |
| | Light (decreasing) | | 0.127 | 5 | 0.046 | $t$ | 2.769 | 0.0363 |
| | Target (taxa) | Random intercept | 0.087 | 1 | - | $\chi^2$ | 54.562 | <0.0001 |
| | Bottle | | 0.152 | 1 | - | $\chi^2$ | 7.027 | 0.0080 |
| | Incubator | | 0.025 | 1 | - | $\chi^2$ | 0.050 | 0.8228 |
| | Richness | Sum of squares | 0.004 | 1, 21.8 | - | $F$ | 0.079 | 0.7811 |
| | Light (decreasing) | | 0.367 | 1, 5.4 | - | $F$ | 7.669 | 0.0363 |
| log10-transformed Cofficient of variation) | Intercept | Fixed effect | -0.429 | 12 | 0.064 | $t$ | -6.723 | <0.0001 |
| | Richness | | -0.007 | 239 | 0.011 | $t$ | -0.680 | 0.4972 |
| | Light (decreasing) | | 0.133 | 236 | 0.027 | $t$ | 4.841 | <0.0001 |
| | Target (taxa) | Random intercept | 0.000 | 1 | - | $\chi^2$ | 90.879 | <0.0001 |
| | Bottle | | 0.221 | 1 | - | $\chi^2$ | 0.000 | 1.0000 |
| | Incubator | | 0.000 | 1 | - | $\chi^2$ | 0.000 | 1.0000 |
| | Richness | Sum of squares | 0.022 | 1, 238.8 | - | $F$ | 0.462 | 0.4972 |
| | Light (decreasing) | | 1.107 | 1, 236 | - | $F$ | 23.440 | <0.0001 |
| Permutation entropy | Intercept | Fixed effect | 0.642 | 14 | 0.023 | $t$ | 27.923 | <0.0001 |
| | Richness | | 0.016 | 238 | 0.004 | $t$ | 3.782 | 0.0002 |
| | Light (decreasing) | | -0.033 | 6 | 0.019 | $t$ | -1.756 | 0.1290 |
| | Target (taxa) | Random intercept | 0.000 | 1 | - | $\chi^2$ | 73.226 | <0.0001 |
| | Bottle | | 0.068 | 1 | - | $\chi^2$ | 0.000 | 1.0000 |
| | Incubator | | 0.022 | 1 | - | $\chi^2$ | 3.945 | 0.0470 |
| | Richness | Sum of squares | 0.100 | 1, 238.1 | - | $F$ | 14.302 | 0.0002 |
| | Light (decreasing) | | 0.022 | 1, 6.1 | - | $F$ | 3.082 | 0.1290 |

Table S17: Table of path coefficients of the structural equation model. The last column reports the standardized coefficients.

| Light conditions | Term | Estimate | SE | $z$-value | $p$-value | Stand. est. |
|---|---|---|---|---|---|---|
| Constant light | Forecast error∼Species richness | 0.015 | 0.031 | 0.469 | 0.6391 | 0.035 |
| | Forecast error∼Coefficient of variation | -0.602 | 0.177 | -3.412 | 0.0006 | -0.416 |
| | Forecast error∼Permutation entropy | 0.085 | 0.659 | 0.128 | 0.8978 | 0.017 |
| | Forecast error∼Autocorrelation (lag 5) | -0.695 | 0.112 | -6.230 | <0.0001 | -0.368 |
| | Coefficient of variation∼Species richness | 0.005 | 0.022 | 0.243 | 0.8084 | 0.018 |
| | Permutation entropy∼Species richness | 0.010 | 0.007 | 1.459 | 0.1446 | 0.117 |
| | Autocorrelation (lag 5)∼Species richness | -0.018 | 0.019 | -0.948 | 0.3429 | -0.081 |
| | Coefficient of variation∼∼Permutation entropy | -0.032 | 0.008 | -4.110 | <0.0001 | -0.784 |
| Decreasing light | Forecast error∼Species richness | -0.077 | 0.035 | -2.224 | 0.0261 | -0.220 |
| | Forecast error∼Coefficient of variation | -0.095 | 0.173 | -0.548 | 0.5834 | -0.072 |
| | Forecast error∼Permutation entropy | 0.430 | 0.569 | 0.756 | 0.4495 | 0.092 |
| | Forecast error∼Autocorrelation (lag 5) | -0.350 | 0.143 | -2.455 | 0.0141 | -0.185 |
| | Coefficient of variation∼Species richness | -0.030 | 0.026 | -1.150 | 0.2500 | -0.111 |
| | Permutation entropy∼Species richness | 0.014 | 0.007 | 2.186 | 0.0288 | 0.191 |
| | Autocorrelation (lag 5)∼Species richness | 0.010 | 0.017 | 0.592 | 0.5536 | 0.053 |
| | Coefficient of variation∼∼Permutation entropy | -0.022 | 0.006 | -3.968 | <0.0001 | -0.628 |

Table S18: Achieved coefficients of determination (R-squared) for the structural equation model.

| Light conditions | Variable | R-squared |
|---|---|---|
| Constant light | Forecast error | 0.3222 |
| | Coefficient of variation | 0.0003 |
| | Permutation entropy | 0.0137 |
| | Autocorrelation (lag 5) | 0.0066 |
| Decreasing light | Forecast error | 0.0972 |
| | Coefficient of variation | 0.0124 |
| | Permutation entropy | 0.0365 |
| | Autocorrelation (lag 5) | 0.0028 |

### S7 Robustness analyses

#### S7.1 Effects of chosen richness variable and forecast method

We investigated if the main finding presented in the main text (i.e. interactive effect of richness and light on taxa abundance forecast error) depended on the forecasting method and the richness measure. Hence we redid the mixed model that investigates the treatment effects on taxa abundance forecast error but using different richness measures and forecasting methods. Figure S21 shows coefficients and corresponding confidence intervals of the various models, and thus the effects of the chosen richness variable and forecast method on the results. The main panel of interest is the bottom-right one, i.e. the one that shows the interaction between light conditions and the species richness. As can be seen, the coefficient estimates were very similar in all cases and the 95% confidence intervals did not overlap with the null effect line (i.e. with zero) in 14 out of 20 cases and only just overlapped in the remaining cases (thus still suggesting some evidence of significance). Therefore, the results seem to be robust regarding the choice of forecast method and richness variable (including the planned species richness).

#### S7.2 Effect of used detrending method

The analysis presented in the main text is based on time series that were detrended with a simple temporal regression. However, arguably this detrending was potentially less adequate for some time series if more than one trend was present over the course of the experiment. Therefore we also did the detrending with an alternative approach, in which we used a segmented temporal regression for which we allowed up to two break points (i.e. up to three different trends) for all time series. The resulting transformed time series are shown in Figure S22 (note: only the biotic time series used as forecast targets are shown).

We then repeated the main analysis (i.e. effect of experimental conditions on taxa abundance forecasts), also considering the different forecasting approaches and the different species richness metrics, as before. The results are shown in the model coefficient Figure S23, which has the same layout as Figure S21 from the previous section. Again, the estimated interaction estimates were very similar irrespective of forecasting method and richness variable and were also very similar to the estimated coefficients based on the simple

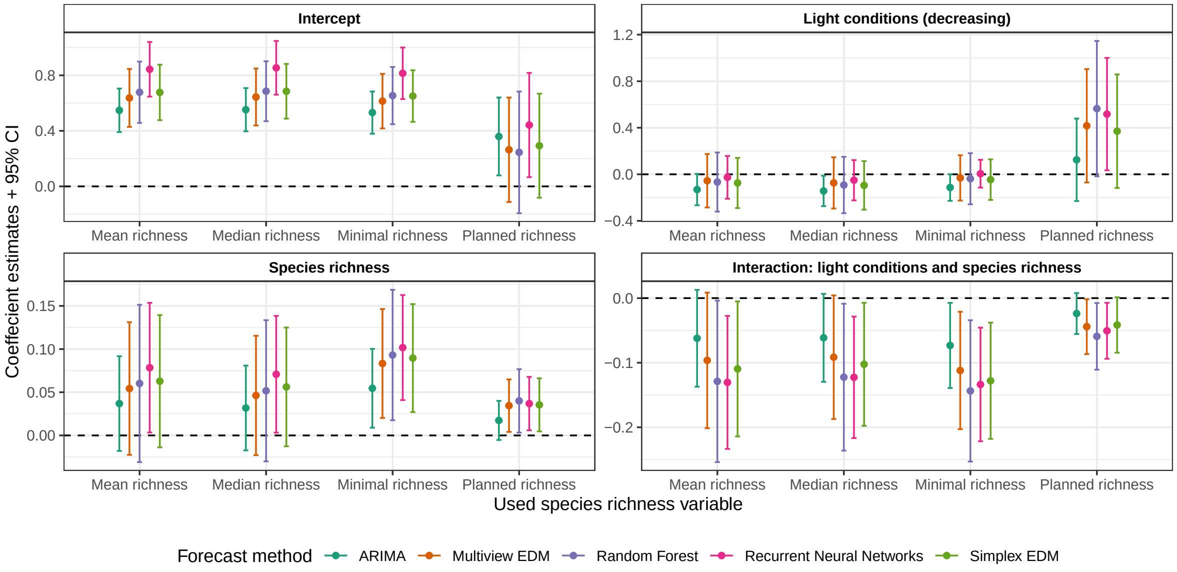

Figure S21: Robustness analysis figure regarding the chosen species richness variable and the forecast method. The figure depicts the coefficient estimates for regressions between the taxa abundance forecast errors (response) and the covariates "species richness" and "light conditions", as specified by the panel labels. The dots are the estimates and the corresponding error-bars are the 95% CI. The colour specifies the used forecast method and the x-axis specifies the used species richness variable.

temporal detrending. We found the interaction to be significant in 11 out of 20 cases, with the $p$-values in the remaining cases being mostly smaller than 0.10 (therefore still suggesting some evidence of significance). Thus, we conclude that the choice of detrending method did not significantly affect the results.

#### S7.3 Species effects on results

We also investigated whether the results were particularly affected by some species. Therefore we repeated the main analysis (i.e. effect of experimental conditions on taxa abundance forecasts) by iteratively excluding single species from the analysis. Figure S24 shows the resulting coefficient plot (which also considered the different richness variables), based on the forecast errors produced by Simplex EDM forecasts. Again, we found that the interaction estimates were very similar regardless of the excluded species and we found it to be significant in 40 out of 56 cases. Out of the remaining 16 cases, the $p$-values were smaller than 0.10 in 13 cases, thus still suggesting some evidence of significance. The taxa that exerted the strongest pull on the results were the Bacteria, but note that even in this case the interaction coefficient estimate was very similar, although with a larger confidence interval.

#### S7.4 Standardization using historical data only

In our main analysis, we standardized time series using the complete dataset to create idealized conditions for investigating treatment effects on forecasting performance. By removing trends and standardizing based

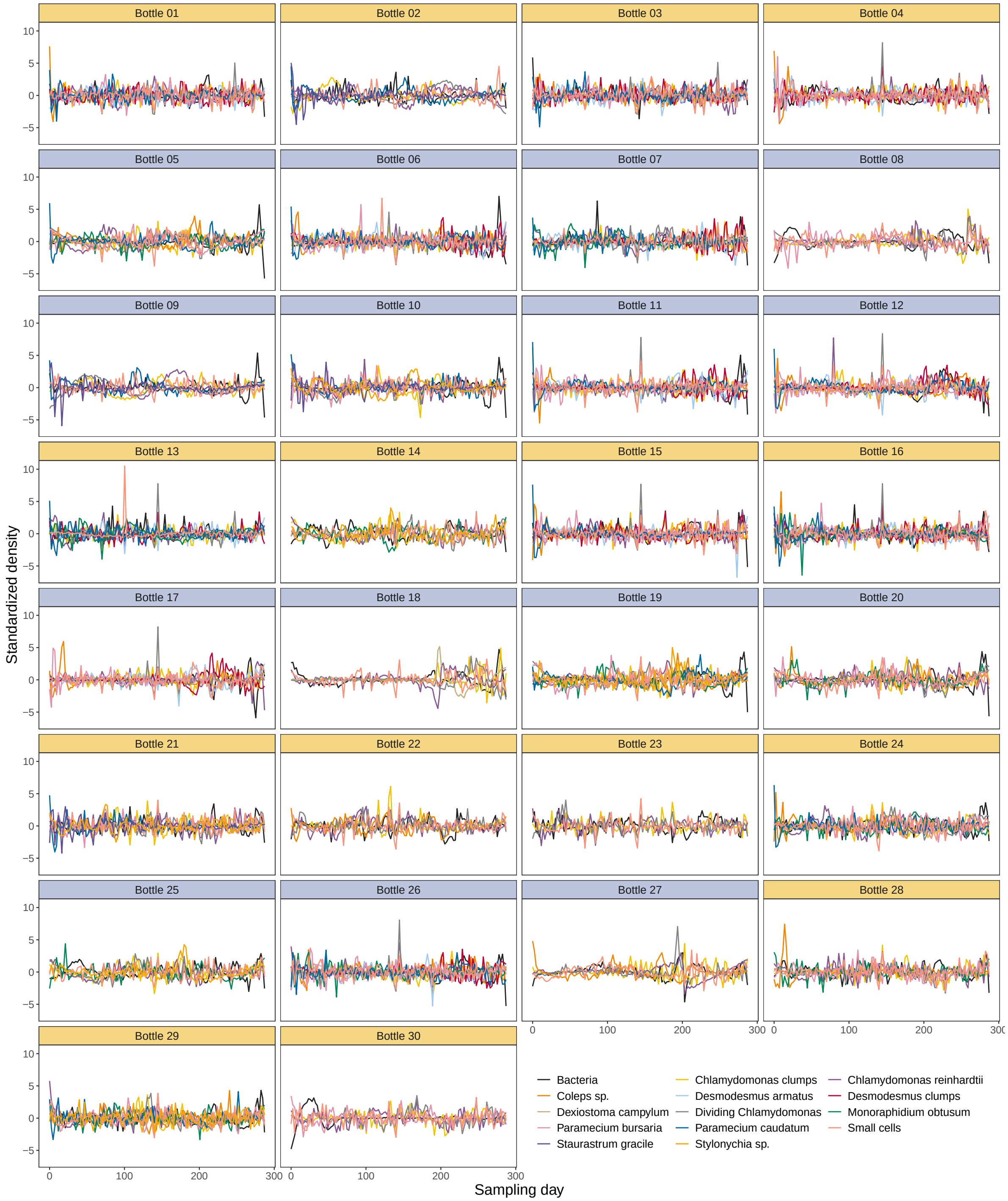

Figure S22: ***Time series detrending based on segmented temporal regression***. The ***transformed*** biotic density time series that were used as forecast targets, shown for the different bottles (i.e. the panels). The time series are coloured by forecast target. The background colour of the subpanel strips indicates whether the bottles were in a constant (orange) or in a decreasing (blue) light environment.

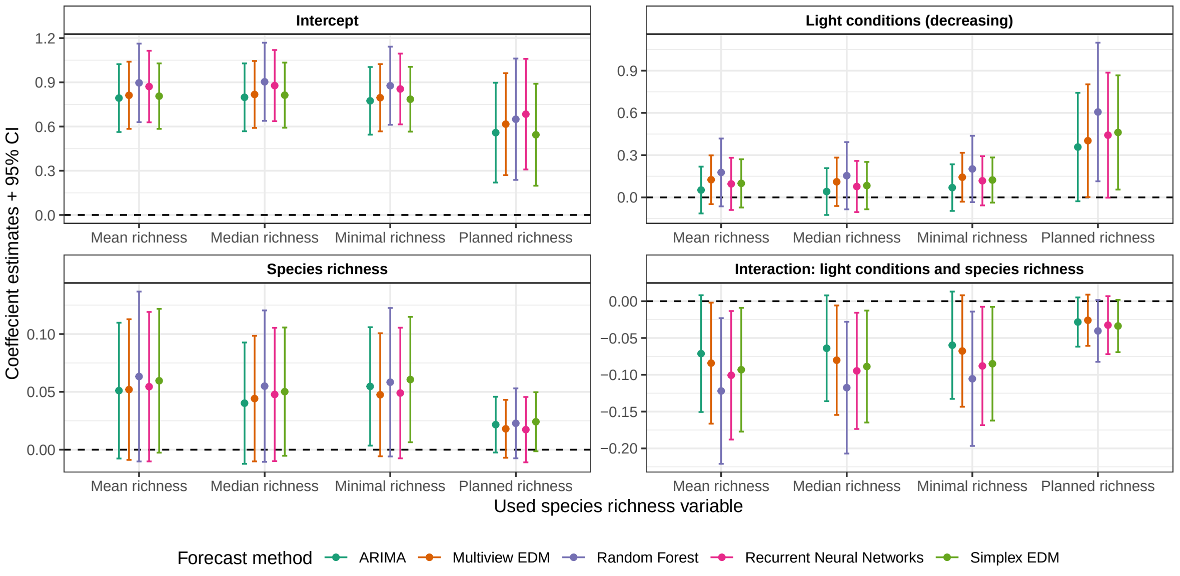

Figure S23: Robustness analysis figure for when the time series were detrended using the **segmented regression** approach. The figure depicts the coefficient estimates for regressions between the taxa abundance forecast errors (response) and the covariates "species richness" and "light conditions", as specified by the panel labels. The dots are the estimates and the corresponding error-bars are the 95% CI. The colour specifies the used forecast method and the x-axis specifies the used species richness variable.

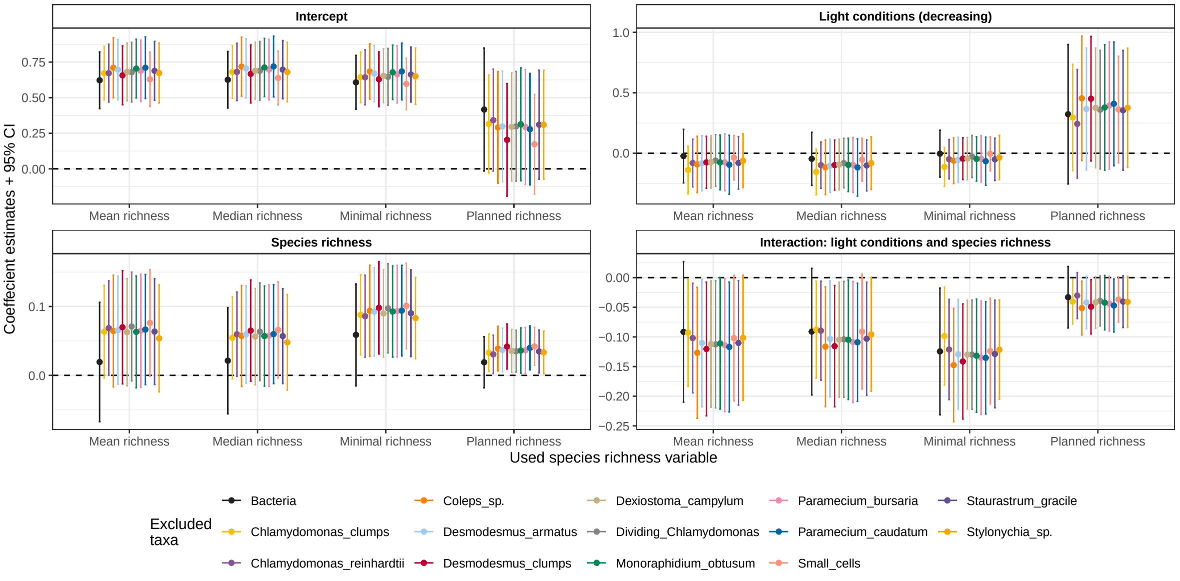

Figure S24: Robustness analysis figure regarding the sensitivity of the results on specific forecasted taxa. The figure depicts the coefficient estimates for regressions between the taxa abundance forecast errors (response) and the covariates "species richness" and "light conditions" (as specified by the panel labels) for when the specified taxa were excluded (i.e. not included in the linear mixed model). The dots are the estimates and the corresponding error-bars are the 95% CI. The colour specifies the taxon that was excluded and the x-axis specifies the used species richness variable. The forecasts are based on the Simplex EDM approach.

on full time series characteristics, we maximized statistical power and minimized variance unrelated to our experimental treatments, creating a controlled environment to test our hypotheses about forecast quality under different conditions. This approach of maximizing statistical power by using the whole time series is often used (see e.g. Munch et al. 2023).

However, using future data for standardization creates information leakage that would be impossible in applied forecasting (Yang, Li, and Jiang 2024). To address this methodological concern, we conducted a robustness analysis using only historical data for detrending and standardization, which better represents real-world forecasting challenges where future patterns are unknown. Figure S25 compares the RMSE values that resulted based on the two approaches, for the different forecast targets across the experimental bottles. As can be seen, there was a strong positive correlation between the RMSE values resulting from the two approaches.

Further, Figure S26 has the same layout as Figure S21, i.e. it shows the coefficients and corresponding confidence intervals of the models investigating the effects of biodiversity (i.e. species richness) and light conditions on forecast errors across different biodiversity measures and forecast methods, but in this case the time series were processed using only historical data (i.e. the 111 time points in the training dataset) rather than the entire dataset. As before, the main panel of interest is the bottom-right one, i.e. the one that shows the interaction between light conditions and the species richness. As can be seen, the results are essentially unchanged and the coefficient estimates were very similar in all cases and the 95% confidence intervals did not overlap with the null effect line (i.e. with zero) in 18 out of 20 cases and only just overlapped in the remaining cases (thus still suggesting some evidence of significance). Therefore, in our case whether we use only historical data for the detrending and standardization or whether we used the entire time series did not impact out results. Thus, the comparison between these two approaches offers insights into how our findings might translate to applied settings and strengthens the validity of our conclusions regarding treatment effects.

#### S7.5 Forecast error evaluation on original and transformed scale

In the main analysis, we evaluated forecast skill using the Root Mean Square Error (RMSE) calculated on detrended and standardized data, which directly assesses the model's ability to capture variations around the trend. This approach aligns with recommendations from forecasting literature suggesting that statistical analysis of forecast performance is often more reliable on transformed scales where data properties are more favorable for statistical inference (Box and Cox 1964; Rob J. Hyndman and Athanasopoulos 2018).

To ensure the robustness of our findings, we carried out an additional analysis by back-transforming forecasts to the original scale before calculating normalized RMSE. The normalization is necessary for the comparison of the achieved forecast errors across time series (of, for example, different species). Because the detrending and standardization consisted in simply using the standardized residuals of the regression of the quantity

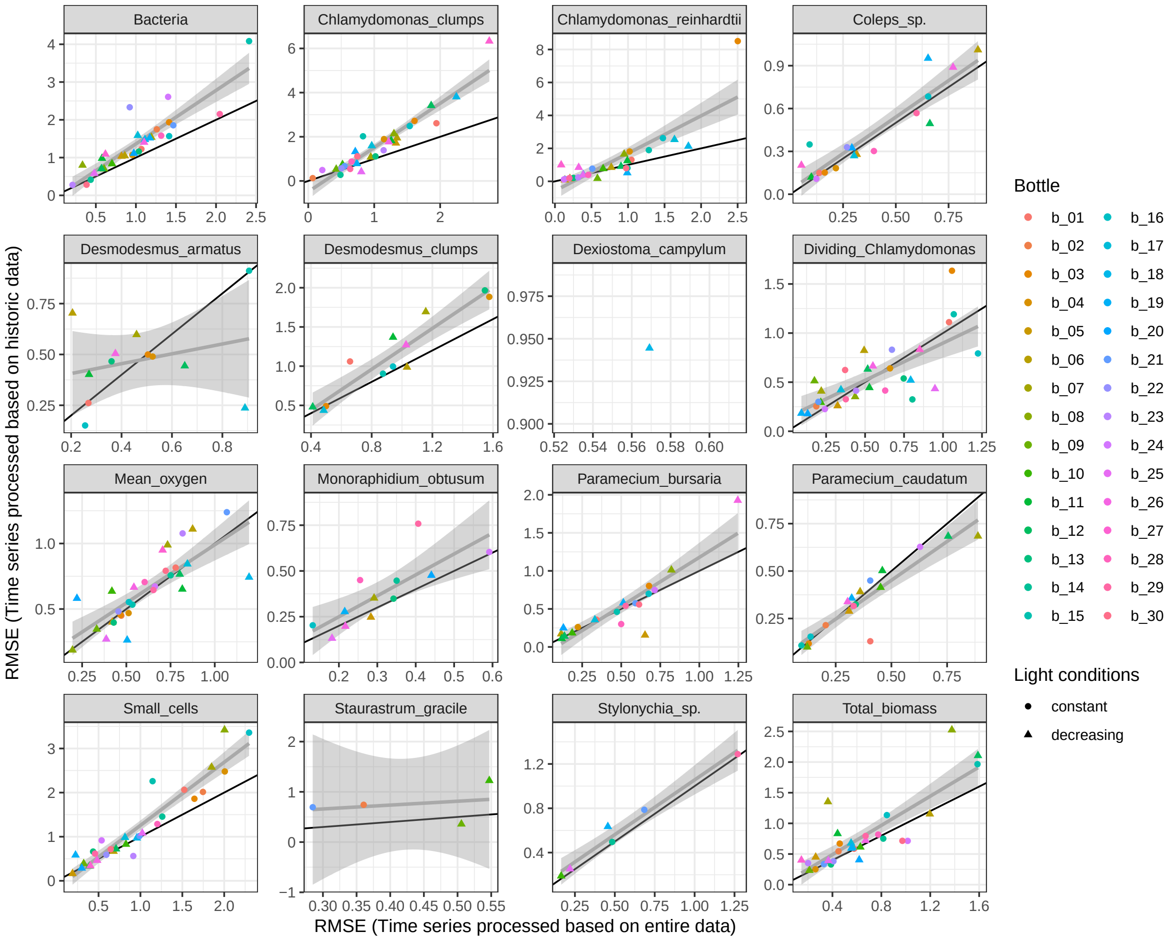

Figure S25: Scatter plot of the RMSE values for when the time series were processed (i.e. detrended, standardized) based on the entire data available against the RMSE values for when the time series were processed using only historic data. RMSE is based on Simplex EDM. Each forecast target is shown in a separate panel, with the colour of the dots indicating the experimental bottle and the shape the light conditions. The black line indicates the identity line, while the grey line (estimate) and the corresponding shaded area (95% CI) are the results of a simple linear regression (fixed effects only).

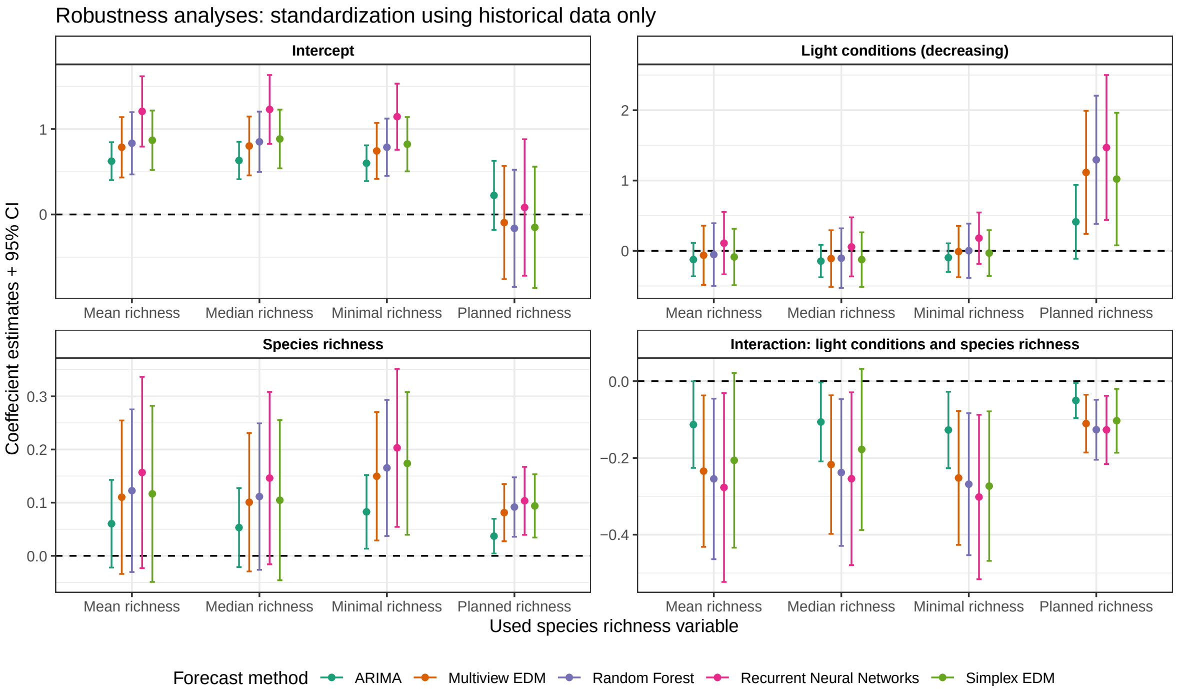

Figure S26: Robustness analysis figure investigating the effects of the standardization using only historical data (instead of the entire time series) on the main results. The relation between biodiversity, light and forecast error is shown for different species richness variables and the different forecast methods. The figure depicts the coefficient estimates for regressions between the taxa abundance forecast errors (response) and the covariates "species richness" and "light conditions", as specified by the panel labels. The dots are the estimates and the corresponding error-bars are the 95% CI. The colour specifies the used forecast method and the x-axis specifies the used species richness variable.

of interest against time (see the main text), the back-transformation is also straightforward: the forecasted values are multiplied by the original standard deviation of the residuals and then the temporal trend is added back. For the subsequent normalization, we divided the RMSE by the standard deviation of the entire original time series (Morley 2019). Note that because the original time series potentially contained trends, their standard deviations are either equal to or larger than those of the above described (trendless) residuals, which means that the calculated normalized RMSE values on the back-transformed scale were either equal or smaller than the ones calculated on the transformed scale. This can be seen in Figure S27, which compares the RMSE values resulting from the two different approaches (i.e. whether forecasts where back-transformed or not before RMSE calculation) for the different forecast targets and across bottles. The Figure also shows that there was a strong positive correlation for the RMSE values coming from the two approaches.

Further, Figure S28 has the same layout as Figure S21, i.e. it shows the coefficients and corresponding confidence intervals of the models investigating the effects of biodiversity (i.e. species richness) and light conditions on forecast errors across different biodiversity measures and forecast methods, but in this case it is for when the normalized RMSE were calculated based on back-transformed forecasts. As before, the main panel of interest is the bottom-right one, i.e. the one that shows the interaction between light conditions and the species richness. As can be seen, the results are essentially unchanged and the coefficient estimates were very similar in all cases and the 95% confidence intervals did not overlap with the null effect line (i.e. with zero) in 17 out of 20 cases and only just overlapped in the remaining cases (thus still suggesting some evidence of significance). Therefore, in our case whether we back-transformed the forecasts or not before calculating the normalized forecast errors did not impact out results.

The relationship between forecast performance metrics on transformed versus original scales is important to consider, as it can affect interpretations of forecast skill drivers. Our dual approach—evaluating on both transformed and original scales—provides complementary perspectives that enhance the robustness of our findings regarding the drivers of forecast skill.

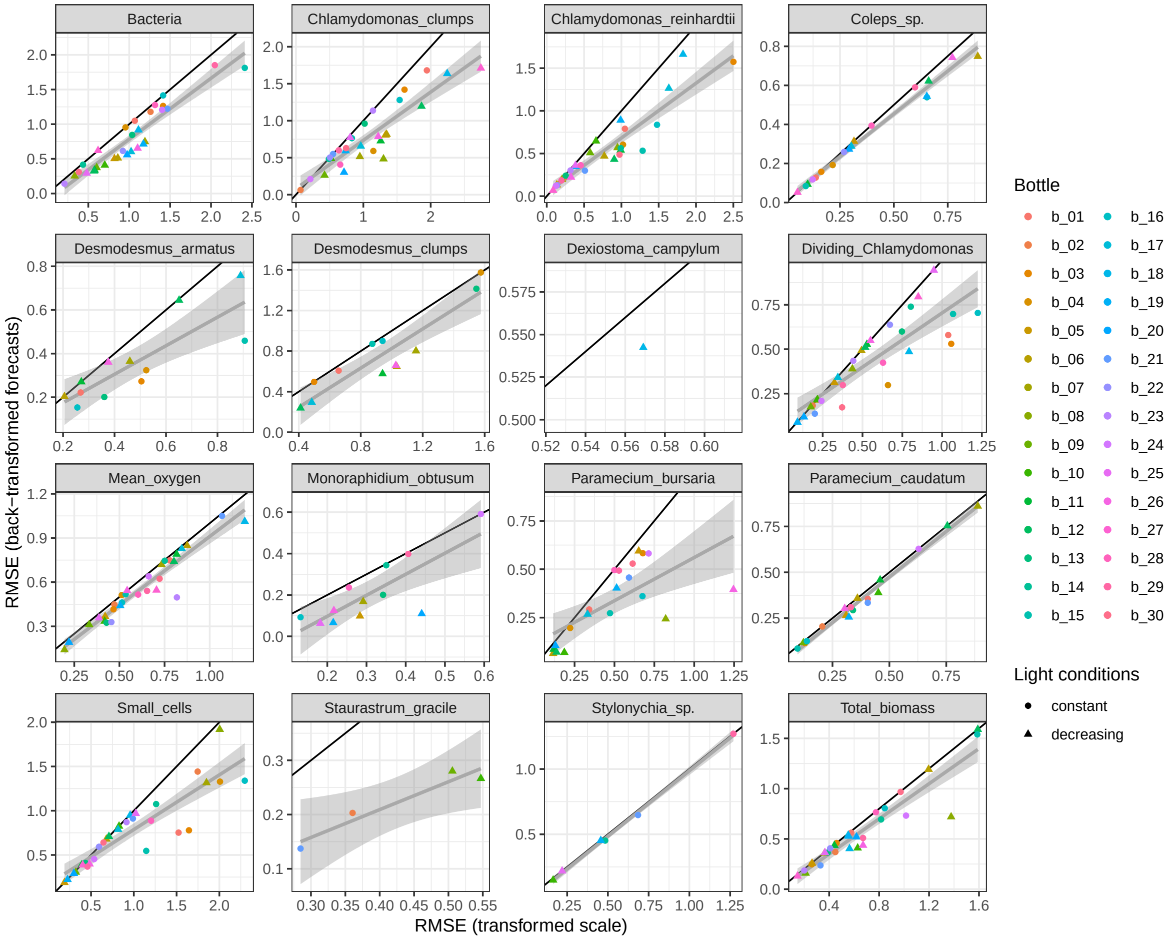

Figure S27: Scatter plot of the RMSE values for the forecasts left on the transformed scale against the RMSE values for the back-transformed forecasts (i.e. the original scale). RMSE is based on Simplex EDM. Each forecast target is shown in a separate panel, with the colour of the dots indicating the experimental bottle and the shape the light conditions. The black line indicates the identity line, while the grey line (estimate) and the corresponding shaded area (95% CI) are the results of a simple linear regression (fixed effects only).

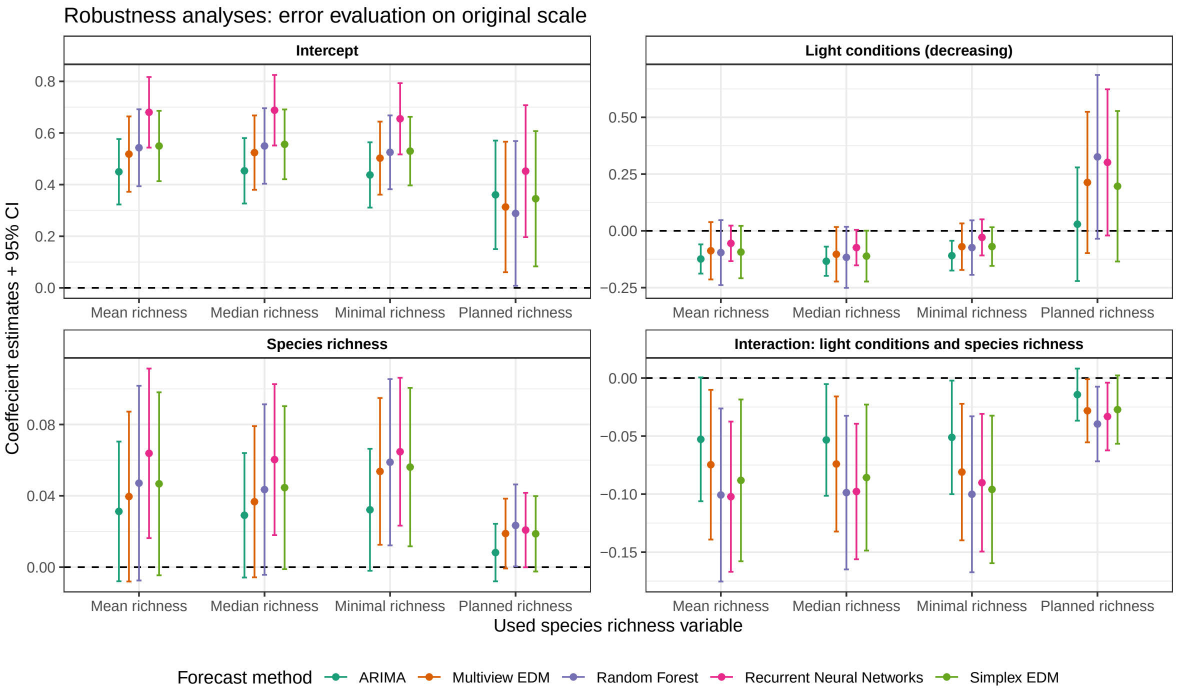

Figure S28: Robustness analysis figure investigating the effects of calculating the RMSE based on back-transformed forecasts (i.e. on the original scale instead of the transformed scale) on the main results. The relation between biodiversity, light and forecast error is shown for different species richness variables and the different forecast methods. The figure depicts the coefficient estimates for regressions between the taxa abundance forecast errors (response) and the covariates "species richness" and "light conditions", as specified by the panel labels. The dots are the estimates and the corresponding error-bars are the 95% CI. The colour specifies the used forecast method and the x-axis specifies the used species richness variable.

### S8 Further figures used in the discussion

#### S8.1 Insurance effect of diversity and possible consequences for forecastability

At the very low end of the diversity gradient, forecast error was higher in decreasing than in constant light. A possible reason was the decline and sudden rise, respectively, of the algae-consumer functional group (i.e. omnivores + mixotrophs) when diversity was low and light decreased. In one of the two compositions with lowest diversity, *Paramecium bursaria* was the only algae-consumer in the community and declined in response to the light decline (Figure S29, bottle 8). This decline in the algae-consumer functional group resulted in pronounced changes in the dynamics of prey organisms (Figure S30, bottle 8). In contrast, in communities with higher diversity, the decline of the dominant algae-consumer (*Paramecium bursaria*) in response to the light decrease was compensated by increases in other algae-consumer species (Figure S29, e.g. bottle 5, bottle 6). These dynamics are in line with the insurance effect of diversity whereby the loss of species due to environmental change is compensated by other species that perform similar functions and tolerate the new environmental conditions (Loreau et al. 2021; Yachi and Loreau 1999). Hence, when diversity was low, lack of insurance might have contributed to the high forecast error by leading to previously unobserved dynamics.

In the second of the two compositions with lowest diversity, the algae-consumer functional group increased suddenly in abundance when light decreased (Figure S29, bottle 18). As long as light intensity was high, algae-consumers were absent in this community. When light declined, the consumer *Euplotes daidaleos* immigrated, probably because the environmental conditions became more favourable for this species (note that all species which were part of a community were allowed to immigrate once every three weeks; see Methods section in the main text for details). Immigration and abundance increase of *Euplotes daidaleos* resulted in pronounced changes in prey abundances (Figure S30, bottle 18), which probably contributed to the low forecast error in this community. In more diverse communities, however, the algae-consumer niche was always filled, irrespective of the light level. Therefore, more-diverse communities were more resistant to the immigration of new algae-consumer species, in line with earlier findings (Beaury et al. 2020; Shurin 2000). Taken together, when diversity was high, the algae-consumer pool contained species that tolerated high-light conditions and species that tolerated low-light conditions. Consequently, algae-consumers were present over the entire light gradient, thereby preventing pronounced changes in lower trophic levels.

#### S8.2 Coefficient of variation of aggregate ecosystem properties

Below we show the relationship between the experimental treatments and the coefficient of variation of total community biomass and oxygen, respectively (Figure S31). We also compare the coefficient of variation at two levels of aggregation, i.e. calculated for community biomass and for population biomass (Figure S32).

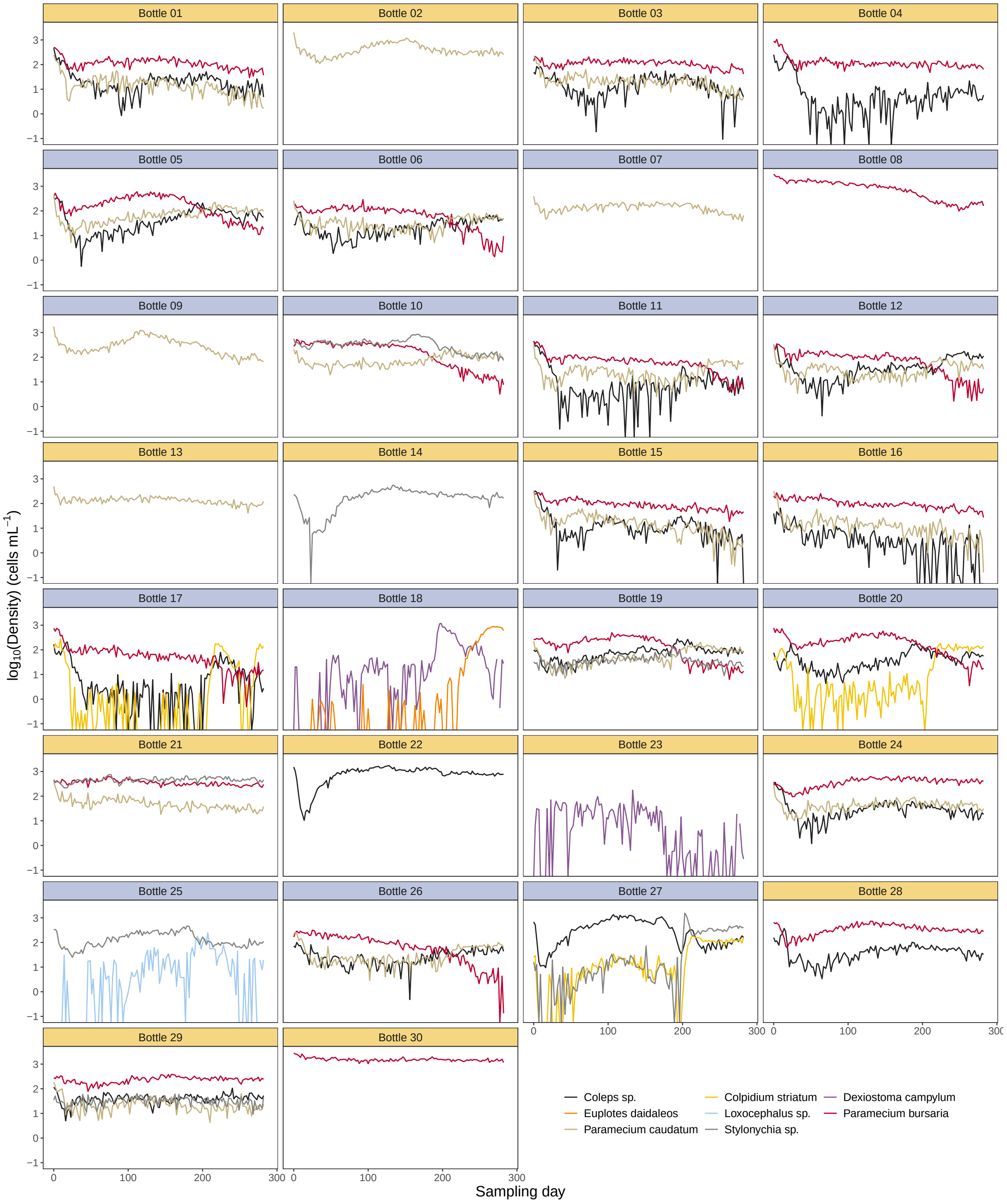

Figure S29: Time series of ciliate densities in the 30 experimental communities. Time series are coloured by ciliate species. The background colour of the subpanel strips indicates whether bottles were maintained in constant light (orange) or in decreasing light (blue).

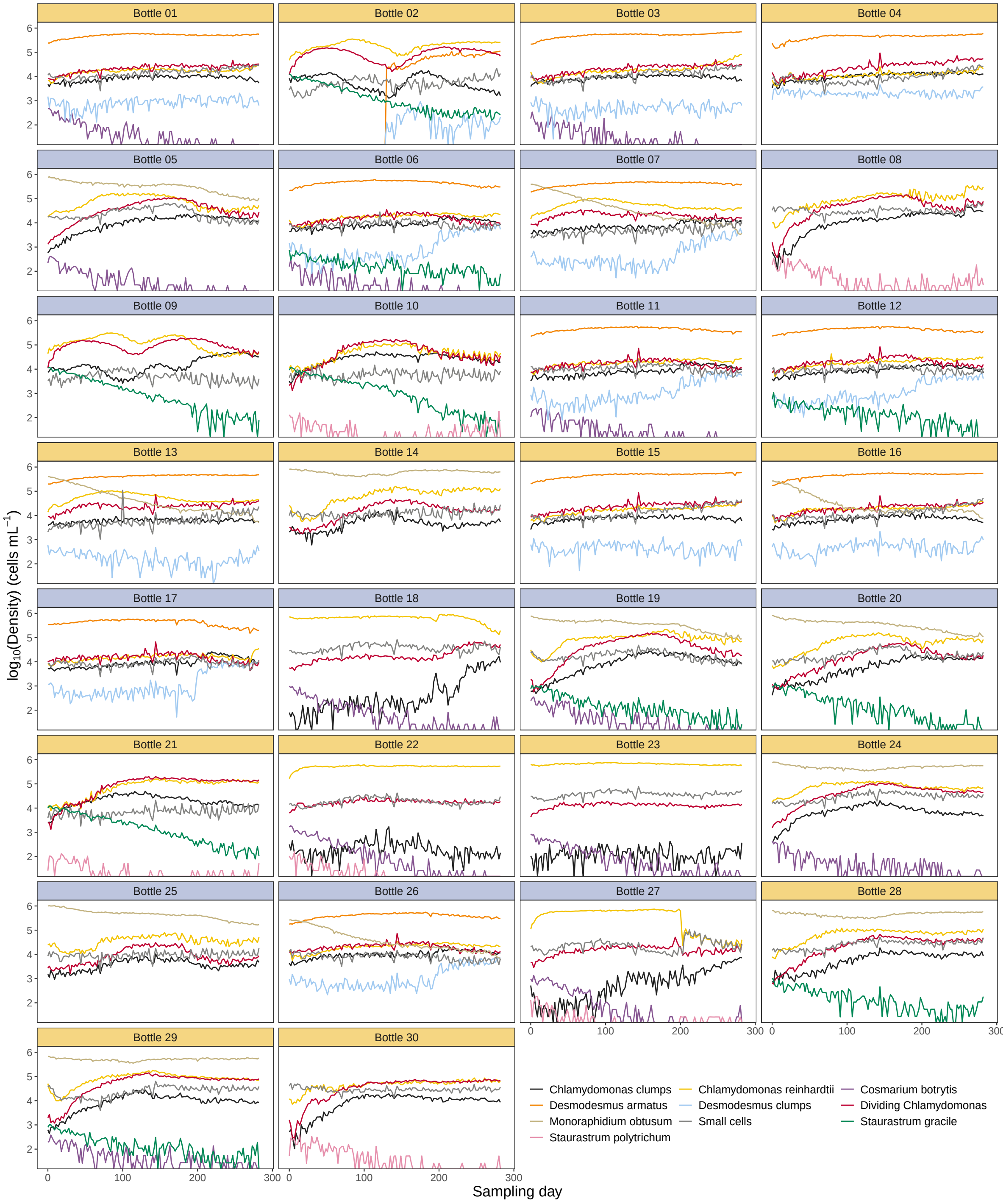

Figure S30: Time series of algae densities in the 30 experimental communities. Time series are coloured by algal species. The background colour of the subpanel strips indicates whether bottles were maintained in constant light (orange) or in decreasing light (blue).

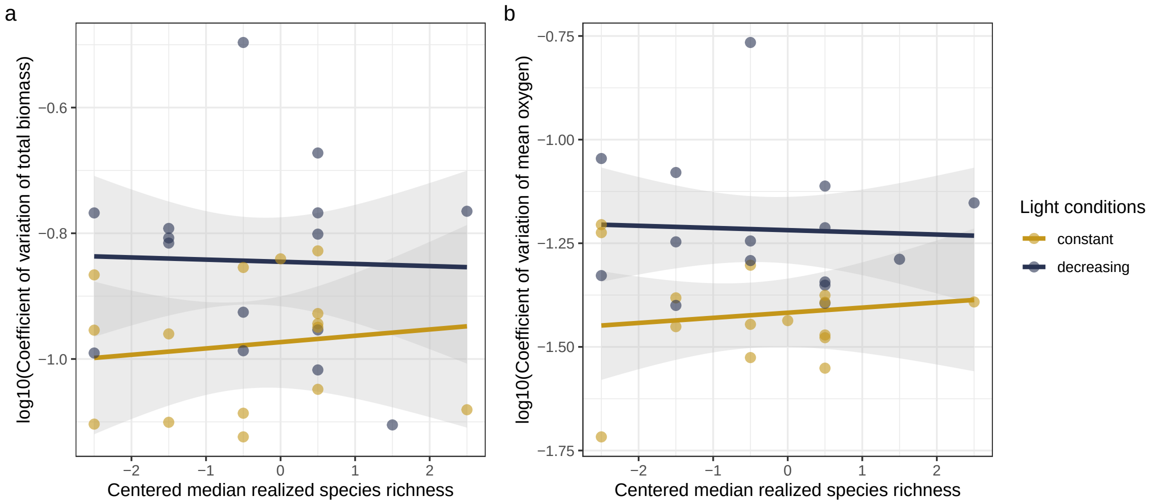

Figure S31: Coefficient of variation of (a) total community biomass, and (b) oxygen concentration. Lines display the fit of the corresponding linear mixed model, shaded areas denote the 95% confidence intervals. Incubator and composition were included as random effects in the model.

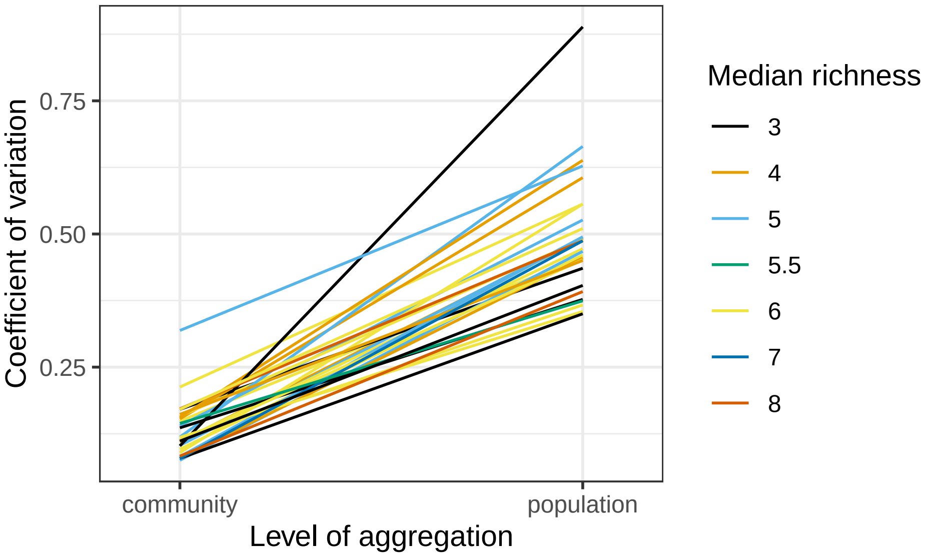

Figure S32: Coefficient of variation (CV) of community and population biomass. For each bottle, we calculated the CV of the time series of total community biomass, and the average CV of the biomass time series of all species in a bottle that were used as forecast targets. Lines connect the two values of a given bottle, and are coloured by median richness.

### S9 Other diversity indices

There are various aspects to biodiversity. In this study, we manipulated species richness while attempting to keep the number of functional groups constant. As previously described and as is common for biological experiments, species did not always persist, and therefore the observed number of functional groups may also deviate from the planned number of functional groups. Indeed, while the number of trophic levels remained the same for all but one bottle throughout the experiment, the median number of functional groups did vary and was correlated with the realized median species richness (Figure S33, note that the light conditions did not influence the number of functional groups). Further, as two additional measures of diversity, we also computed the median Shannon evenness for the taxa abundances and the taxa biomasses (i.e. we computed these measures for each sampling day and then took the median of these calculated values). The correlations and distributions of the abundance and biomass evenness, the median species richness and (as mentioned) the median number of functional groups are all given in Figure S33. As can be seen, these measures were generally positively correlated, although not always significantly so. We also computed the Shannon diversity for the abundance and biomass, but because of their very strong correlation with the respective evenness indices they are not further shown and mentioned here.

Next, we used these additional indices of diversity instead of the median species richness variable in the forecast error analysis to investigate the relations between light conditions and the diversity index with the response variable (taxa abundance forecast error). The results are given in Figure S34. We generally found the same patterns as when we used the median species richness (which is also included in Figure S34 for comparison), but only in the case of the median number of functional groups these were also significant (which was expected based on the correlation between median species richness and median number of functional groups shown in Figure S33).

### S10 Versions of R and of the R-packages used

**R version**: R version 4.4.1 (2024-06-14 ucrt).

**Base packages**: grid, parallel, stats, graphics, grDevices, utils, datasets, methods, base.

**Other packages**: r2glmm 0.1.2, effectsize 1.0.0, lavaan.survey 1.1.3, lavaan 0.6-19, survey 4.4-2, survival 3.8-3, bemovi 1.0, flowViz 1.70.0, flowCore 2.8.0, keras 2.15.0, reticulate 1.42.0, slider 0.3.2, broom.mixed 0.2.9.6, GGally 2.2.1, ggh4x 0.3.0, lmerTest 3.1-3, lme4 1.1-37, Matrix 1.7-3, caret 7.0-1, lattice 0.22-6, here 1.0.1, pals 1.10, kableExtra 1.4.0, rEDM 1.15.4, doParallel 1.0.17, iterators 1.0.14, foreach 1.5.2, forecast 8.23.0, pracma 2.4.4, data.table 1.17.0, patchwork 1.3.0, randomForest 4.7-1.2, lubridate 1.9.4, forcats 1.0.0, stringr 1.5.1, dplyr 1.1.4, purrr 1.0.4, readr 2.1.5, tidyr 1.3.1, tibble 3.2.1, ggplot2 3.5.1, tidyverse 2.0.0, ggstance 0.3.7, emmeans 1.11.0, knitr 1.50.

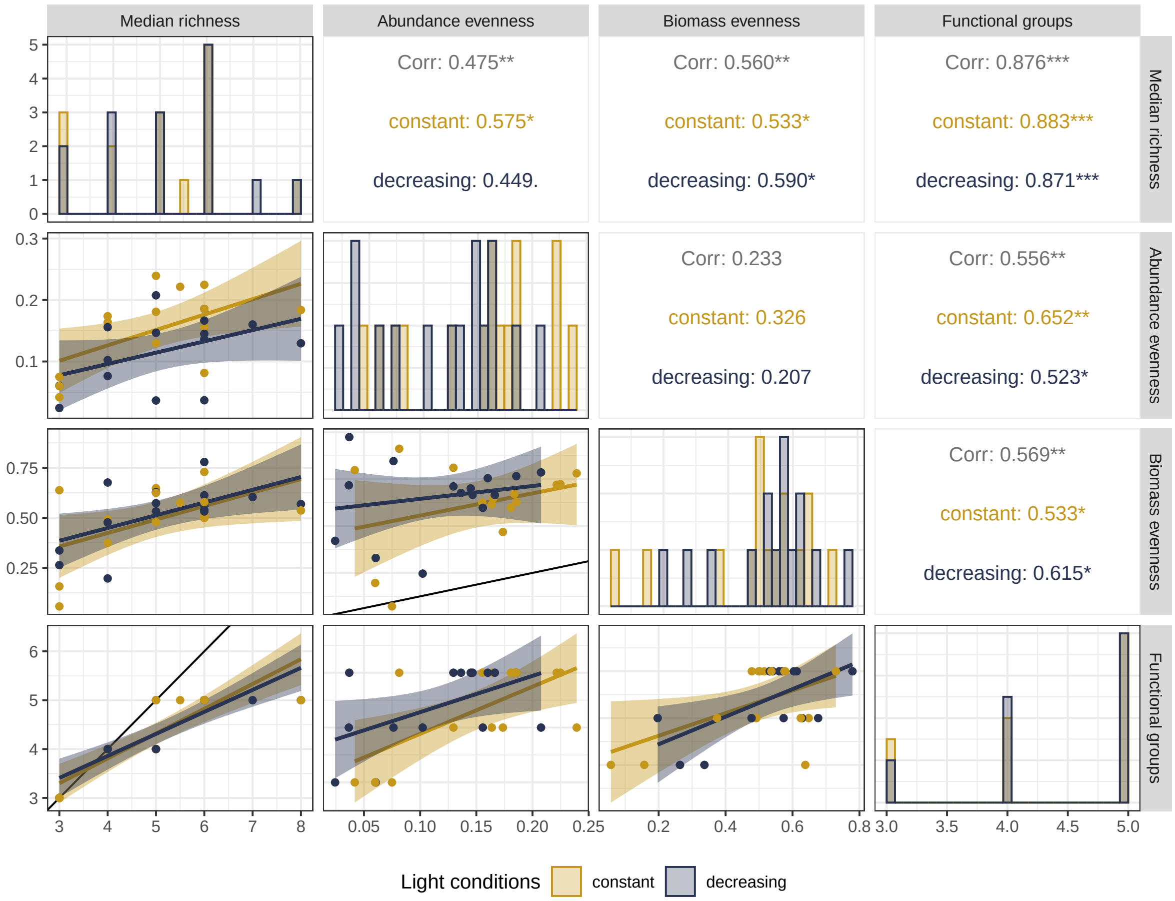

Figure S33: Correlations and distributions of different diversity indices (i.e. the median species richness, the median Shannon evenness of the species abundances and biomasses, and the median number of functional groups), colour-coded for the light conditions. The panels in the diagonal show the bar-plots of the richness variables. The panels below the diagonal show the pairwise scatter-plots as indicated by the facet labels (the black line is the identity line, the coloured lines and the corresponding shaded areas show the fit and the 95% CI of a linear regression). The panels above the diagonal show the corresponding correlation estimates.

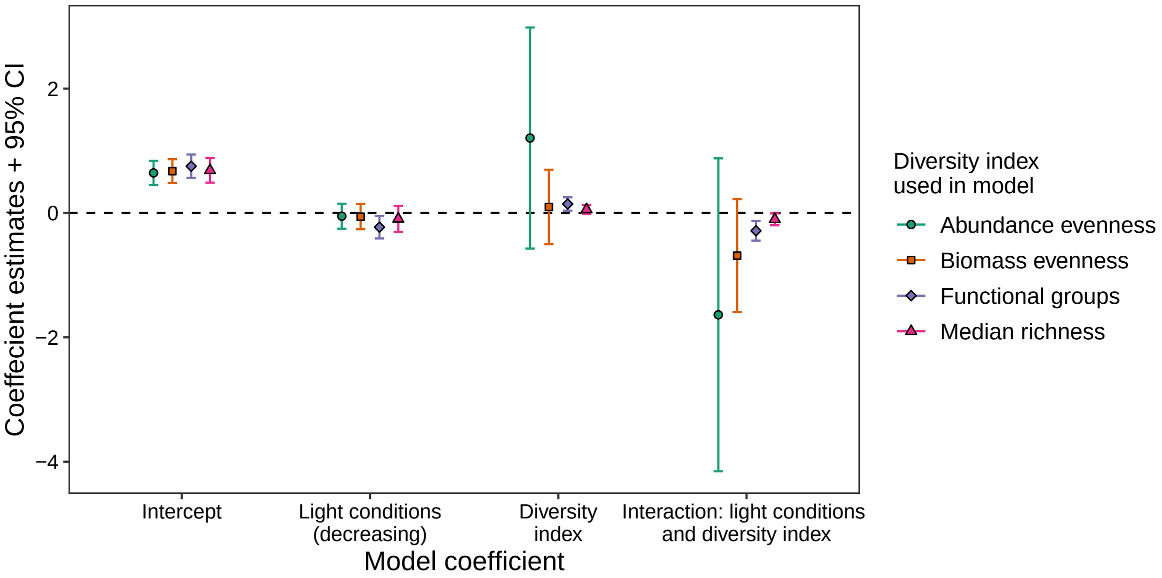

Figure S34: Results of forecast error analyses using different diversity indices (i.e. the median species richness, the median Shannon evenness of the species abundances and biomasses, and the median number of functional groups). The figure depicts the coefficient estimates for regressions between the taxa abundance forecast errors (response) and the covariates "diversity index" and "light conditions". The dots are the estimates and the corresponding error-bars are the 95% CI. The colour and shape of the dots specify the used diversity index and the x-axis specifies the model coefficient.

**This document was generated on April 11, 2025 at 18:30**.

Donohue, Ian, Owen L Petchey, José M Montoya, Andrew L Jackson, Luke McNally, Mafalda Viana, Kevin Healy, Miguel Lurgi, Nessa E O'Connor, and Mark C Emmerson. 2013. "On the Dimensionality of

Ecological Stability." *Ecology Letters* 16 (4): 421–29.

Ellis, B, Perry Haaland, Florian Hahne, Nolwenn Le Meur, Nishant Gopalakrishnan, Josef Spidlen, Mike Jiang, and Greg Finak. 2023. *flowCore: flowCore: Basic Structures for Flow Cytometry Data*. https://doi.org/10.18129/B9.bioc.flowCore.

Hewamalage, Hansika, Christoph Bergmeir, and Kasun Bandara. 2021. "Recurrent Neural Networks for Time Series Forecasting: Current Status and Future Directions." *International Journal of Forecasting* 37 (1): 388–427.

Hillebrand, Helmut, Claus-Dieter Dürselen, David Kirschtel, Utsa Pollingher, and Tamar Zohary. 1999. "Biovolume Calculation for Pelagic and Benthic Microalgae." *Journal of Phycology* 35 (2): 403–24.

Hyndman, R, G Athanasopoulos, C Bergmeir, G Caceres, L Chhay, M O'Hara-Wild, F Petropoulos, S Razbash, E Wang, and F Yasmeen. 2022. *Forecast: Forecasting Functions for Time Series and Linear Models*. https://pkg.robjhyndman.com/forecast.

Hyndman, Rob J., and George Athanasopoulos. 2018. *Forecasting: Principles and Practice*. 2nd ed. Melbourne, Australia: OTexts. https://otexts.com/fpp2/.

Hyndman, Rob J, and Yeasmin Khandakar. 2008. "Automatic Time Series Forecasting: The Forecast Package for r." *Journal of Statistical Software* 27: 1–22.

Jaeger, Byron. 2017. *R2glmm: Computes r Squared for Mixed (Multilevel) Models*. https://CRAN.R-project.org/package=r2glmm.

Krug, Rainer M. 2024. *LEEF.measurement.bemovi: Prepares Movies for Analysis with Bemovi and Extracts Data*. https://github.com/LEEF-UZH/LEEF.measurement.bemovi/releases/tag/v1.1.0LEEF_1.

Lenth, Russell V. 2023. *Emmeans: Estimated Marginal Means, Aka Least-Squares Means*. https://CRAN.R-project.org/package=emmeans.

Liaw, Andy, and Matthew Wiener. 2002. "Classification and Regression by randomForest." *R News* 2 (3): 18–22. https://CRAN.R-project.org/doc/Rnews/.

Loreau, Michel, Matthieu Barbier, Elise Filotas, Dominique Gravel, Forest Isbell, Steve J Miller, Jose M Montoya, et al. 2021. "Biodiversity as Insurance: From Concept to Measurement and Application." *Biological Reviews* 96 (5): 2333–54.

Meyer, David, Evgenia Dimitriadou, Kurt Hornik, Andreas Weingessel, and Friedrich Leisch. 2021. *E1071: Misc Functions of the Department of Statistics, Probability Theory Group (Formerly: E1071), TU Wien*. https://CRAN.R-project.org/package=e1071.

Morley, James. 2019. "Normalizing the RMSE." Marine Data Science blog. https://www.marinedatascience.co/blog/2019/01/07/normalizing-the-rmse/.

Munch, Stephan B., Tanya L. Rogers, Celia C. Symons, David Anderson, and Frank Pennekamp. 2023. "Constraining Nonlinear Time Series Modeling with the Metabolic Theory of Ecology." *Proceedings of the National Academy of Sciences* 120 (12): e2211758120. https://doi.org/10.1073/pnas.2211758120.

Oberski, Daniel. 2014. "Lavaan. Survey: An r Package for Complex Survey Analysis of Structural Equation

Models." *Journal of Statistical Software* 57: 1–27.

Park, Joseph, Cameron Smith, George Sugihara, and Ethan Deyle. 2021. *rEDM: Empirical Dynamic Modeling ('EDM')*. https://CRAN.R-project.org/package=rEDM.

Pennekamp, Frank, Alison C Iles, Joshua Garland, Georgina Brennan, Ulrich Brose, Ursula Gaedke, Ute Jacob, et al. 2019. "The Intrinsic Predictability of Ecological Time Series and Its Potential to Guide Forecasting." *Ecological Monographs* 89 (2): e01359.

Pennekamp, Frank, Nicolas Schtickzelle, and Owen L. Petchey. 2015. "BEMOVI, Software for Extracting Behavior and Morphology from Videos, Illustrated with Analyses of Microbes." *Ecology and Evolution* 5 (13): 2584–95. https://doi.org/10.1002/ece3.1529.

Rights, Jason D., and Sonya K. Sterba. 2019. "Quantifying Explained Variance in Multilevel Models: An Integrative Framework for Defining R-Squared Measures." *Psychological Methods* 24 (3): 309–38. https://doi.org/10.1037/met0000184.

Shurin, Jonathan B. 2000. "Dispersal Limitation, Invasion Resistance, and the Structure of Pond Zooplankton Communities." *Ecology* 81 (11): 3074–86.

Sugihara, George, and Robert M May. 1990. "Nonlinear Forecasting as a Way of Distinguishing Chaos from Measurement Error in Time Series." *Nature* 344 (6268): 734–41.

Takens, Floris. 1981. "Detecting Strange Attractors in Turbulence." In *Dynamical Systems and Turbulence, Warwick 1980*, 366–81. Springer.

Yachi, Shigeo, and Michel Loreau. 1999. "Biodiversity and Ecosystem Productivity in a Fluctuating Environment: The Insurance Hypothesis." *Proceedings of the National Academy of Sciences* 96 (4): 1463–68.

Yang, Xinyi, Jingyi Li, and Xuchu Jiang. 2024. "Research on Information Leakage in Time Series Prediction Based on Empirical Mode Decomposition." *Scientific Reports* 14 (1): 28362. https://doi.org/10.1038/s41598-024-80018-9.

Ye, Hao, and George Sugihara. 2016. "Information Leverage in Interconnected Ecosystems: Overcoming the Curse of Dimensionality." *Science* 353 (6302): 922–25.
